# DIALECT: DNA-Programmable Synthetic Cells Communicate with Mammalian Cells through Synthetic Receptors

**DOI:** 10.64898/2026.11.06.764853

**Authors:** Jesse Lentjes, Anna-Maria Makri Pistikou, Bas W. A. Bögels, Maaruthy Yelleswarapu, Shuo Yang, Alexander Gräwe, Thijs W. van Veldhuisen, Federica Eduati, Maarten Merkx, Jan C. M. van Hest, Stephen Mann, Tom F. A. de Greef

## Abstract

Synthetic cells can regulate mammalian signalling through the controlled release of diffusible protein ligands, but coupling programmable synthetic-cell senders to orthogonal, engineered mammalian receptors remains challenging. Here we present **DIALECT** (DNA-Instructed Assembly of Ligands for Engineered Cellular Transduction), a platform in which proteinosome-based synthetic cells translate sequence-defined DNA inputs into coiled-coil signalling ligand outputs for engineered mammalian receptors. DNA–ligand conjugates are localized within semipermeable proteinosomes through DNA hybridization and released by toehold-mediated DNA strand displacement (TMSD). The released ligands diffuse to neighbouring mammalian cells and activate cognate CC–GEMS receptors, inducing JAK/STAT3 signalling and reporter activation, with a higher response in cognate than non-cognate receiver cells under activation conditions. We further implement distributed Boolean AND logic using two independently programmed proteinosome populations that respond to distinct DNA inputs and release complementary ligand components. Simultaneous addition of both inputs produces substantially higher reporter activity than the respective single-input conditions, consistent with extracellular assembly of an active receptor agonist. **DIALECT** thus establishes a modular interface linking DNA-based molecular computation in synthetic cells to orthogonal signalling pathways in mammalian cells, providing a foundation for hybrid artificial–biological communication networks.

## Introduction

Engineered synthetic cells increasingly emulate the molecular communication strategies through which living cells coordinate sensing, information processing and collective behaviour. Communication has been demonstrated using diverse compartment architectures, including lipid vesicles, polymer-based capsules, coacervates and proteinosomes, with small molecules, nucleic acids and proteins serving as signalling species.^[1–8]^ Such systems have enabled controlled signal release and molecular exchange,^[9,10]^ while DNA offers sequence-specific recognition and programmable reaction networks for processing distinct molecular inputs. In semipermeable proteinosomes,^[11]^ DNA hybridization and strand displacement have enabled sequence-selective signalling between synthetic-cell populations.^[12–15]^ Early interfaces with living cells predominantly involved bacterial receivers, typically using quorum-sensing molecules or other diffusible signals.^[16–21]^

Synthetic–living cell interfaces have since expanded to mammalian systems through protein production, nucleic-acid delivery and diffusible or surface-presented molecular signals.^[22–27]^ Notably, ATP-fuelled DNA reaction networks coupled to artificial-cell microgels enable controlled release of TNFα–ssDNA conjugates and activation of NF-κB signalling in mammalian reporter cells,^[26]^ while stimulus-responsive DNA artificial cells use contact-dependent presentation of aptameric ligands to regulate native mammalian receptors.^[27]^ Although these systems demonstrate programmable artificial–mammalian signalling, both rely on endogenous receptor recognition, and the latter additionally requires direct contact. Related platforms have also been explored for localized therapeutic production and immune-cell activation.^[28–31]^ In parallel, engineered receptor systems enable mammalian cells to recognize defined extracellular cues and convert them into programmable intracellular responses.^[32–40]^ Soluble-ligand-responsive receptors include Generalized Extracellular Molecule Sensors (GEMS) ^[32]^, coiled-coil GEMS (CC–GEMS)^[33]^ and Synthetic Intramembrane Proteolysis Receptors (SNIPRs)^[35]^, whereas synthetic Notch (synNotch) receptors predominantly mediate contact-dependent signalling through ligands presented on neighbouring cells or synthetic materials.^[34,36]^ Additional receptor-engineering strategies have further expanded programmable extracellular sensing and cellular control.^[37–40]^ Combining DNA-addressable, diffusible ligand release from bottom-up synthetic cells with orthogonal receptor recognition would allow sender programming and receiver specificity to be engineered independently, providing a basis for modular, multi-input communication.

Here we introduce **DIALECT** (DNA-Instructed Assembly of Ligands for Engineered Cellular Transduction), a communication system that couples DNA-programmable proteinosome senders to mammalian cells expressing orthogonal CC–GEMS receptors (Fig. 1a). CC–GEMS receptors couple programmable coiled-coil (CC) recognition to customizable GEMS-mediated cellular signalling. Sequence-defined DNA inputs trigger toehold-mediated DNA strand displacement (TMSD), releasing soluble bifunctional ssDNA–coiled-coil signalling ligands that diffuse to neighbouring mammalian cells and activate cognate CC–GEMS receptors, initiating Janus kinase/signal transducer and activator of transcription 3 (JAK/STAT3) signalling and downstream gene expression. We further distribute signal processing across two independently programmed proteinosome populations, each releasing a complementary ligand component in response to a distinct DNA input.

**Figure 1.**
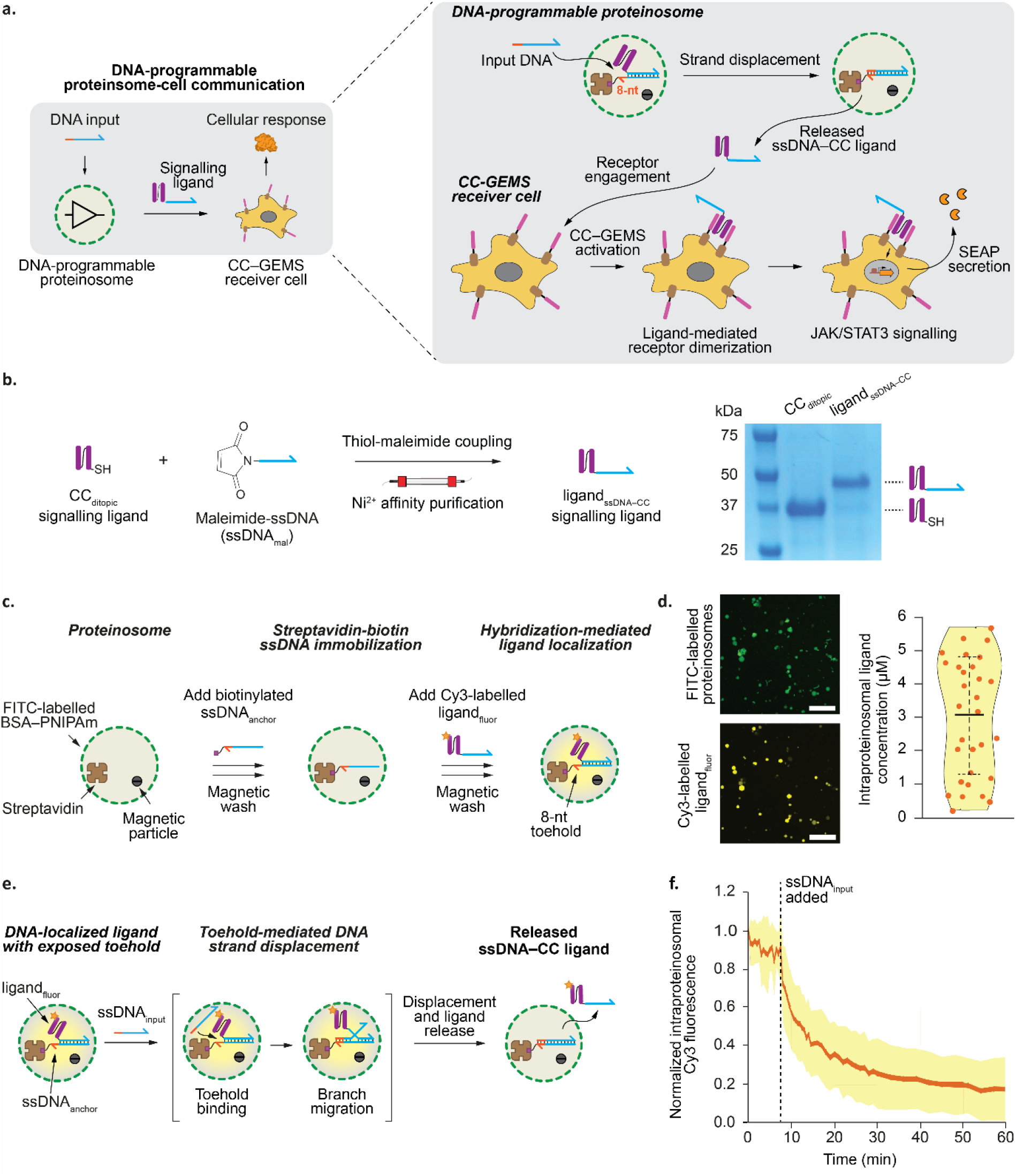
DNA-programmable localization and release of synthetic signalling ligands from proteinosomes. (a) Schematic of proteinosome–mammalian cell communication. BSA-NH_2_/PNIPAm proteinosomes containing streptavidin and magnetic microparticles immobilize biotinylated 34-nt DNA anchors (**ssDNA**_**anchor**_). An ssDNA–coiled-coil (CC) ligand (**ligand**_**ssDNA–CC**_) hybridizes through a complementary 26-nt domain, leaving an exposed 8-nt toehold. Cognate 34-nt input DNA (**ssDNA**_**input**_) triggers TMSD-mediated release, allowing the ligand to diffuse to receiver cells and activate CC–GEMS receptor dimerization, JAK/STAT3 signalling and SEAP secretion. (b) Cysteine-modified ditopic CC ligand (**CC**_**ditopic**_) was conjugated to maleimide-functionalized ssDNA (26 nt) by thiol–maleimide coupling and purified by Ni^2+^-affinity chromatography. SDS–PAGE showed apparent molecular masses of approximately 37 kDa (**CC**_**ditopic**_) and 50 kDa (**ligand**_**ssDNA–CC**_). Uncropped gel, Fig. S22. (c) FITC-labelled proteinosomes encapsulating streptavidin and magnetic microparticles were loaded with Cy3-labelled **ligand**_**fluor**_ through streptavidin–biotin immobilization of **ssDNA**_**anchor**_ followed by ligand hybridization. Magnetic retrieval after each loading step removed unbound DNA and ligand. Cy3 was attached to **CC**_**ditopic**_ using Cy3–NHS ester. (d) Representative fluorescence micrographs and intraproteinosomal ligand concentrations determined by calibrated Cy3 fluorescence (Fig. S4). Violin plots show individual proteinosomes (*n* = 31), with horizontal lines indicating the mean and error bars s.d. Mean ligand concentration, 3.08 ± 1.79 µM (mean ± s.d.). Scale bar, 250 µm. (e) Cognate input DNA binds the exposed 8-nt toehold and displaces the ligand-associated DNA strand from **ssDNA**_**anchor**_ by branch migration, allowing the released ssDNA–CC ligand to diffuse through the semipermeable proteinosome membrane. (f) Ligand-release kinetics of proteinosomes containing **ligand**_**fluor**_ in a microfluidic trapping array (Fig. S6). Complementary input DNA (100 nM) was added at t = 8 min. Intraproteinosomal Cy3 fluorescence was monitored over time and normalized to the steady-state no-input signal. Data represent individual proteinosomes (*n* = 15); solid line, mean; shading, s.d. Rapid fluorescence loss supports TMSD-mediated ligand release. Representative images and individual trajectories, Fig. S7. Representative time-lapse recording, Supplementary Movie 1.

Simultaneous addition of both inputs enables AND-gated mammalian-cell activation, consistent with extracellular assembly of an active receptor agonist. **DIALECT** thus integrates programmable molecular processing in bottom-up synthetic cells with orthogonal mammalian receptor signalling and distributed molecular logic.

## Results and Discussion

### DNA-Programmable Localization and Release of Synthetic Signalling Ligands from Proteinosomes

To establish direct communication between DNA-programmable synthetic cells and mammalian cells, we engineered a bifunctional ssDNA–coiled-coil (ssDNA–CC) signalling ligand that couples sequence-selective DNA strand displacement to CC–GEMS receptor activation (Fig. 1a). The ligand contains a 26-nt ssDNA recognition domain for hybridization to a 34-nt anchor within proteinosomes, leaving an exposed 8-nt toehold for a cognate 34-nt input strand, and an engineered CC domain for receptor engagement.^[33]^ Input binding initiates toehold-mediated DNA strand displacement (TMSD), releasing the ligand from the anchor.^[41–43]^ The released ligand can subsequently diffuse to neighbouring mammalian receiver cells and induce CC–GEMS receptor dimerization and JAK/STAT3-dependent SEAP secretion.^[33]^ All constructs used in this study are listed in Supplementary Tables S1–S5.

To generate the bifunctional ligand, we introduced a single cysteine into the previously reported A′A′ CC construct^[33]^ between its His and SUMO tags near the N-terminus, providing a defined attachment site for DNA conjugation while minimizing interference with the receptor-binding interface (Fig. S1).^[44–49]^ The resulting **CC**_**ditopic**_ protein was expressed in *E. coli* and purified by Ni^2+^-affinity chromatography, with SDS–PAGE showing migration at approximately 37 kDa (Fig. S2; Methods). In parallel, 26-nt amine-modified ssDNA (**ssDNA**_**Am**_) was reacted with sulfo-SMCC to generate maleimide-functionalized ssDNA (**ssDNA**_**mal**_), which was conjugated to **CC**_**ditopic**_ by thiol–maleimide coupling (Fig. 1b).^[50–53]^ Following sequential affinity purification to remove excess ssDNA (Methods), SDS–PAGE showed a shift to approximately 50 kDa, consistent with formation of the bifunctional **ligand**_**ssDNA–CC**_ conjugate (Fig. 1b; Fig. S3). For fluorescence experiments, the CC domain was labelled with Cy3–NHS ester to yield **ligand**_**fluor**_ (DoL 4.5; Methods).

We next examined DNA-mediated ligand localization in hollow BSA-NH_2_/PNIPAm proteinosomes, which provide semipermeable compartments that allow controlled loading and release of molecular cargo, but exhibit size-dependent macromolecular permeability (Fig. 1c).^[10–12,54,55]^ Biotinylated **ssDNA**_**anchor**_ was immobilized through streptavidin–biotin interactions within FITC-labelled proteinosomes containing streptavidin and magnetic microparticles, followed by hybridization-mediated loading of **ligand**_**fluor**_ (Fig. 1c; Methods).^[10,12,13]^ Magnetic retrieval after each loading step removed unbound DNA and ligand species (Methods).^[54]^ Fluorescence microscopy revealed strong Cy3 fluorescence localized within the proteinosomes (Fig. 1d), and calibration against a Cy3 standard (Fig. S4) yielded a mean intraproteinosomal ligand concentration of 3.08 ± 1.79 μM (mean ± s.d.; *n* = 31). Variation in loading may reflect differences in proteinosome size, morphology, membrane permeability and streptavidin content.^[11,12,54,55]^ Low-level ligand localization was detected in controls lacking streptavidin and cognate anchor DNA (Fig. S5), potentially reflecting cysteine-dependent interactions with the BSA-rich matrix, including thiol–disulfide exchange at BSA Cys34.^[56–58]^ Nevertheless, ligand localization was 3.7-fold higher in the presence of both streptavidin and cognate anchor DNA, supporting DNA hybridization as the predominant retention mechanism.

Having established DNA-mediated ligand localization, we tested sequence-specific release from proteinosomes (Fig. 1e). The anchor–ligand duplex exposes an 8-nt toehold that allows cognate **ssDNA**_**input**_ to initiate TMSD, displacing the ligand-bearing strand and enabling the ssDNA–CC conjugate to diffuse into the surrounding medium.^[41–43]^ Because proximity of the CC domain to the toehold could hinder strand displacement,^[59,60]^ we examined ligand release experimentally. FITC-labelled proteinosomes loaded with **ligand**_**fluor**_ were confined in a previously established microfluidic trapping array^[10,12,13]^ (Fig. S6; Methods), enabling real-time fluorescence monitoring following controlled input delivery.^[10,12,13,54]^ Addition of complementary **ssDNA**_**input**_ caused a rapid decrease in intraproteinosomal Cy3 fluorescence (Fig. 1f; Fig. S7; Supplementary Movie 1; Methods), supporting TMSD-mediated ligand displacement and release. The sequence specificity of DNA-triggered ligand release was further supported by subsequent experiments showing ligand retention under non-complementary input conditions during proteinosome–mammalian cell co-incubation (Fig. 3b) and orthogonal release of distinct ligand components in response to their cognate DNA inputs (Fig. S19).

Together, these results establish DNA-programmable localization within and input-triggered release of a synthetic signalling ligand from proteinosomes, providing a mechanism to control the availability of a diffusible CC–GEMS agonist.

### Establishing Conditions for Stable Proteinosome–Cell Communication

To identify cell culture conditions compatible with proteinosome–mammalian cell communication, we assessed the stability of the ssDNA–CC ligand in different media (Fig. 2a,b). Although the CC domain retains receptor activity under mammalian culture conditions,^[32,33]^ serum nucleases can degrade the ssDNA domain.^[61–66]^ Preserving the ssDNA domain is important for hybridization-mediated ligand retention within sender proteinosomes and input-selective TMSD release, independently of the receptor-activating function of the CC domain. We therefore tested media with reduced or absent serum,^[61–64]^ including formulations supplemented with non-cognate ssDNA as a sacrificial nuclease substrate (Fig. 2a).^[63,66]^ Following overnight incubation at 37 °C, Serum-free DMEM, serum-free 293 SFM II (SFM), and low-serum DMEM supplemented with 50% PBS and 1 μM non-cognate ssDNA retained >90% intact ligand (Fig. 2b; Methods). The latter formulation, termed protection medium, contained 1.25% heat-inactivated FBS, 50% PBS and 1 μM **ssDNA**_**scramble**_ in DMEM.

**Figure 2.**
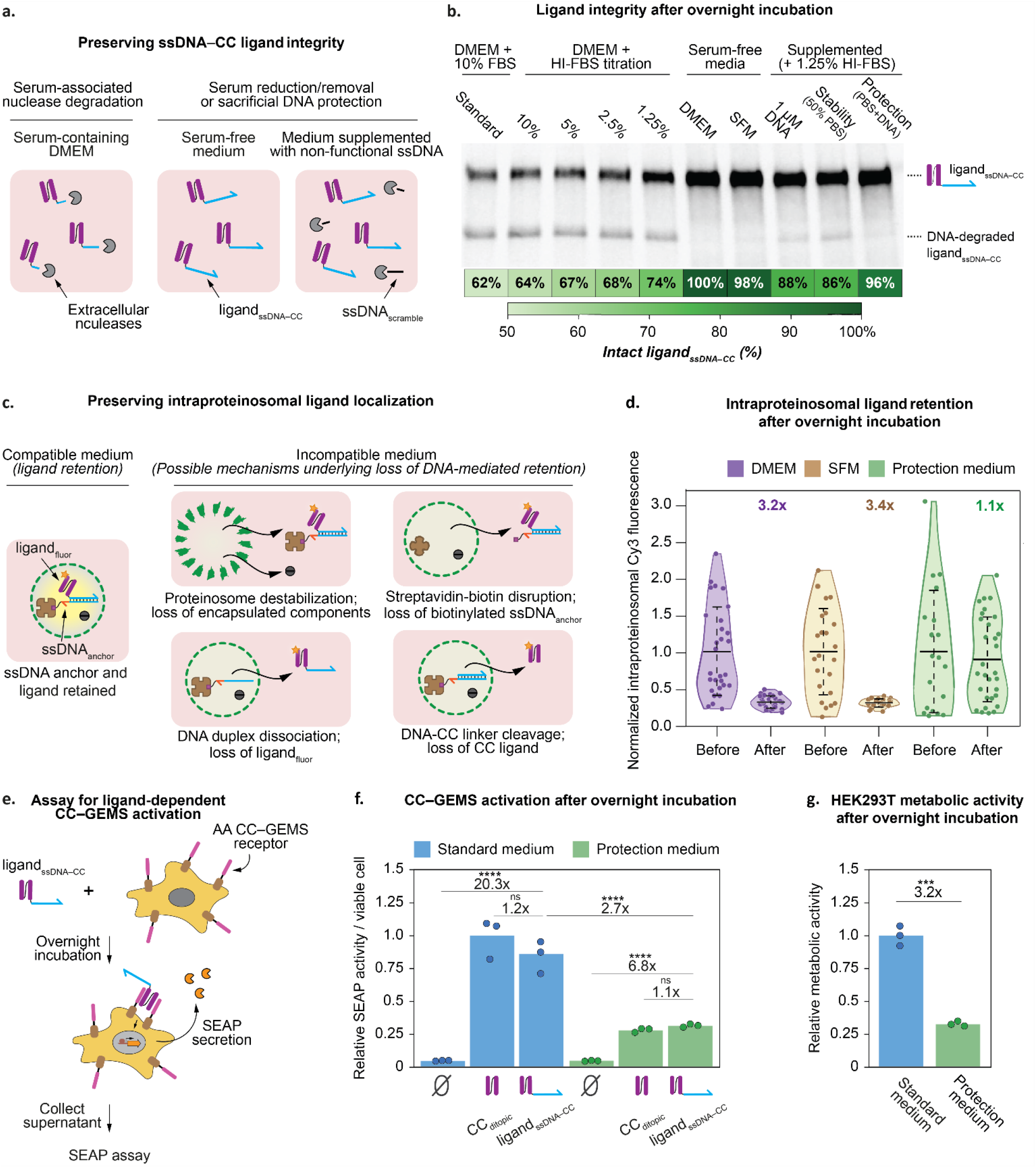
Engineering conditions for DNA-programmable proteinosome–cell communication. (a) Schematic of nuclease-mediated degradation of the ssDNA domain in serum-containing medium and its protection by serum reduction or addition of non-complementary ssDNA (34-nt, **ssDNA**_**scramble**_) as a sacrificial nuclease substrate without triggering TMSD. (b) Anti-SUMO Western blot of **ligand**_**ssDNA–CC**_ (1 µM) following overnight incubation in standard medium (DMEM containing 10% FBS), DMEM with decreasing concentrations of heat-inactivated FBS (HI-FBS), serum-free DMEM, serum-free 293 SFM II (SFM), or modified low-serum DMEM. Stability medium consisted of DMEM containing 1.25% HI-FBS and 50% PBS; protection medium additionally contained 1 µM **ssDNA**_**scramble**_. Media containing individual additives were also tested. Samples were diluted tenfold in PBS before electrophoresis. Upper and lower bands correspond to intact and DNA-degraded **ligand**_**ssDNA–CC**_, respectively. The heat map shows ligand integrity calculated as upper/(upper + lower) × 100 from densitometric analysis. Media preserving >90% intact ligand were selected for further experiments. Uncropped blot, Fig. S24. (c) Schematic of potential loss of intraproteinosomal ligand retention during mammalian cell culture (37 °C, 5% CO_2_) through proteinosome destabilization, disruption of streptavidin–biotin anchoring or DNA hybridization, or cleavage of the ssDNA–CC linkage. These processes may cause premature ligand release and compromise DNA-programmable control over mammalian cell activation. (d) FITC-labelled proteinosomes containing Cy3-labelled **ligand**_**fluor**_ were incubated overnight in serum-free DMEM, SFM or protection medium. Intraproteinosomal Cy3 fluorescence was normalized to its pre-incubation value. Serum-free media caused ligand loss, whereas protection medium maintained localization without substantial TMSD-mediated release by **ssDNA**_**scramble**_. Encapsulated streptavidin was retained but biotinylated cargo was lost in serum-free media, consistent with disruption of streptavidin–biotin anchoring (Fig. S9). Violin plots show individual proteinosomes (DMEM, *n* = 29 before/30 after; SFM, *n* = 23/30; protection medium, *n* = 19/30). Horizontal lines indicate means and error bars denote s.d. Fold reductions were calculated as ratios of mean pre-incubation to post-incubation fluorescence. Representative micrographs, Fig. S8. (e) HEK293T cells transiently expressing CC–GEMS receptors were incubated overnight with cognate ssDNA–CC ligand. Secreted SEAP activity was quantified by measuring absorbance at 405 nm. (f) AA CC–GEMS-expressing HEK293T cells were incubated overnight with 120 nM **CC**_**ditopic**_ or **ligand**_**ssDNA–CC**_ in standard or protection medium. SEAP activity was normalized to independently measured viable-cell numbers (NucleoCounter NC-100; Methods) and expressed relative to **CC**_**ditopic**_ in standard medium. Both ligands elicited comparable responses within each medium, and **ligand**_**ssDNA–CC**_ remained largely intact during activation (Fig. S10). Protection medium reduced ligand-induced SEAP output per viable cell 2.7-fold. Data represent biological triplicates. Statistical significance was determined by one-way ANOVA across all six conditions with Tukey’s multiple-comparison test. (g) HEK293T cells were incubated overnight in standard or protection medium, and MTT activity was measured per well without normalization to viable-cell number. MTT activity was normalized to the mean signal in standard medium. Protection medium reduced MTT activity 3.2-fold. Data represent biological triplicates. Statistical significance was determined using a two-sided Welch’s t-test. ns, P > 0.05; *P ≤ 0.05; **P ≤ 0.01; ***P ≤ 0.001; ****P ≤ 0.0001.

We next examined whether these conditions maintained DNA-mediated ligand retention within proteinosomes during overnight incubation under standard mammalian culture conditions (Fig. 2c,d; Methods). Intraproteinosomal fluorescence decreased 3.2- and 3.4-fold in serum-free DMEM and SFM, respectively, compared with only 1.1-fold in protection medium (Fig. 2d; Fig. S8). To investigate ligand loss, we examined proteinosomes containing fluorescently labelled streptavidin or biotinylated cargo (Fig. S9; Methods). Streptavidin remained encapsulated, whereas biotinylated cargo was lost in both serum-free media, consistent with disruption of streptavidin–biotin anchoring. This may reflect the finite dissociation rate of the streptavidin–biotin interaction,^[67–71]^ with medium composition influencing cargo rebinding.^[67]^ Protection medium best maintained DNA-mediated ligand retention and was selected for subsequent experiments.

Finally, we examined whether DNA conjugation affected CC–GEMS activation. AA CC–GEMS-expressing HEK293T cells were incubated overnight with 120 nM **CC**_**ditopic**_ or **ligand**_**ssDNA–CC**_ in standard or protection medium (Fig. 2e,f; Methods). Both ligands produced comparable SEAP responses within each medium, indicating that DNA conjugation did not substantially impair receptor activation. Anti-SUMO Western blotting confirmed that most DNA-conjugated ligand remained intact during signalling (Fig. S10; Methods). Relative to untreated controls, ssDNA–CC ligand increased SEAP activity per viable cell 20.3-fold in standard medium and 6.8-fold in protection medium. SEAP output per viable cell was 2.7-fold lower in protection medium, while MTT activity per well decreased 3.2-fold (Fig. 2f,g; Methods). These changes broadly paralleled each other, consistent with altered cellular metabolic state contributing to the attenuated signalling under protection-medium conditions of reduced serum concentration and DMEM nutrient content. However, MTT activity also depends on viable-cell number and therefore does not independently establish reduced metabolic activity per cell.

Together, these results identify protection medium as suitable for proteinosome–mammalian cell communication, largely preserving ligand integrity and DNA-mediated retention while supporting CC–GEMS activation, albeit with reduced signalling output. We therefore used protection medium for subsequent DNA-triggered ligand release and proteinosome–mammalian cell communication experiments.

### Programmed Communication between DNA-Programmable Proteinosomes and Synthetic Receptor-Engineered Mammalian Cells

Having established conditions compatible with ligand retention and receptor activation, we next investigated whether DNA-triggered ligand release remained functional during proteinosome–mammalian cell co-incubation (Fig. 3a). HEK293T cells were co-incubated overnight with FITC-labelled proteinosomes containing **ligand**_**fluor**_ in either protection medium or activation medium, the latter comprising stability medium supplemented with 1 μM cognate **ssDNA**_**input**_ (Methods). Intraproteinosomal ligand concentration remained high in protection medium but decreased 8.3-fold in activation medium (Fig. 3b; Fig. S11), supporting ligand retention in the absence of a cognate input and TMSD-mediated release during co-incubation with mammalian cells.

**Figure 3.**
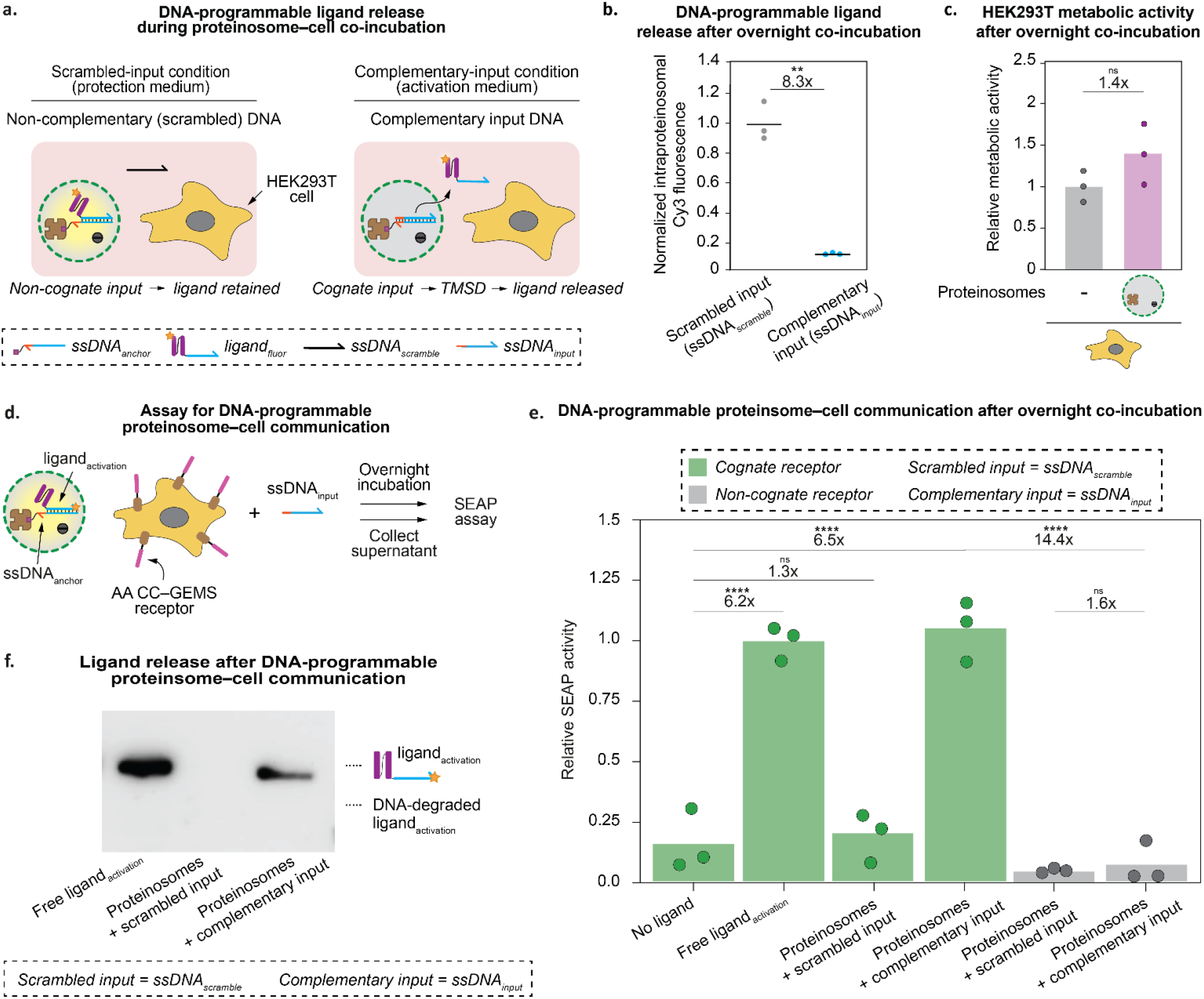
DNA-programmable proteinosome–cell communication by input-triggered ligand release. (a) Schematic of ligand retention and DNA-triggered release during proteinosome–mammalian cell co-incubation. Non-complementary input (protection medium; stability medium supplemented with 1 μM **ssDNA**_**scramble**_) maintains ligand retention, whereas complementary input (activation medium; stability medium supplemented with 1 μM **ssDNA**_**input**_) triggers TMSD-mediated release. Co-incubation with mammalian cells does not induce substantial ligand leakage under scrambled-input conditions. (b) HEK293T cells were co-incubated overnight with FITC-labelled proteinosomes containing **ligand**_**fluor**_ under scrambled- or complementary-input conditions. Intraproteinosomal Cy3 fluorescence was normalized to the scrambled-input condition. Complementary input reduced fluorescence 8.3-fold, consistent with sequence-specific ligand release. Biological triplicates; horizontal lines, means; dots, individual replicates. Two-sided Welch’s t-test. Representative micrographs, Fig. S11. (c) HEK293T cells were incubated overnight in stability medium with or without FITC-labelled proteinosomes. MTT activity was normalized to cells incubated in stability medium alone and remained comparable upon proteinosome co-incubation, consistent with compatibility under the tested conditions. Biological triplicates; bars, means; dots, individual replicates. Two-sided Welch’s t-test. Representative micrographs, Fig. S12. (d) Assay for proteinosome–cell communication. CC–GEMS-expressing HEK293T receiver cells were co-incubated overnight with ligand-containing proteinosomes and input DNA. Secreted SEAP was quantified by absorbance at 405 nm. (e) Cognate AA CC–GEMS receiver cells were incubated without ligand or with 120 nM free Cy3-labelled ssDNA–CC ligand (**ligand**_**activation**_) under scrambled-input conditions, or with FITC-labelled proteinosomes containing **ligand**_**activation**_ under scrambled- or complementary-input conditions. Non-cognate BB CC–GEMS receiver cells were tested with ligand-containing proteinosomes under both input conditions. **Ligand**_**activation**_ was prepared by conjugating **CC**_**ditopic**_ to Cy3-labelled maleimide-functionalized 26-nt ssDNA (**ssDNA**_**activation**_). SEAP activity was normalized to the 120 nM free-ligand condition. Proteinosomes under complementary-input conditions elicited a response in cognate receiver cells comparable to that induced by free ligand, whereas proteinosomes under scrambled-input conditions did not induce activation. Non-cognate receptors showed low activity under both input conditions, without a significant difference. Biological triplicates; bars, means; dots, individual replicates. One-way ANOVA across all six conditions with Tukey’s multiple-comparison test. (f) Representative anti-SUMO Western blot of culture supernatants from cognate CC–GEMS receiver cells following the co-incubations in (e). Ligand was detected under free-ligand and complementary-input conditions, whereas no detectable ligand was observed under scrambled-input conditions. Uncropped blot, Fig. S24. ns: P > 0.05; *P ≤ 0.05; **P ≤ 0.01; ***P ≤ 0.001; ****P ≤ 0.0001.

To assess proteinosome–cell compatibility, we incubated HEK293T cells overnight in stability medium with or without FITC-labelled proteinosomes and measured cellular metabolic activity using an MTT assay (Fig. 3c; Fig. S12; Methods). Proteinosome co-incubation did not substantially alter cellular metabolic activity, consistent with compatibility under the conditions tested.

To estimate the number of proteinosomes required for communication, we selected a target ligand concentration of 120 nM, within the concentration range previously shown to support robust CC–GEMS activation.^[33]^ Using a 100 μL assay volume to calculate the total ligand requirement (Fig. S13), together with the measured intraproteinosomal ligand concentration (3.08 ± 1.79 μM; Fig. 1d) and proteinosome diameter distribution (23.35 ± 6.60 μm; Fig. S13), we estimated the required proteinosome number. Assuming complete ligand release, these parameters yielded an estimated requirement of approximately 4.69 × 10^5^ ligand-containing proteinosomes, which was used in subsequent communication experiments. To test sequence-dependent receptor activation, FITC-labelled proteinosomes containing **ligand**_**activation**_ were co-incubated overnight with HEK293T cells expressing cognate AA or non-cognate BB CC–GEMS receptors, with ligand release controlled by sequence-specific DNA input (Fig. 3d; Methods). In **ligand**_**activation**_, the Cy3 fluorophore was confined to the ssDNA domain rather than the CC domain, enabling fluorescent tracking without modifying the receptor-binding CC domain.

Co-incubation in protection medium did not significantly activate cognate CC–GEMS receptors relative to the no-ligand control, consistent with ligand retention within proteinosomes (Fig. 3e). In contrast, co-incubation in activation medium increased SEAP activity 6.5-fold, comparable to the 6.2-fold increase elicited by 120 nM free ligand. Non-cognate CC–GEMS receptors showed no substantial activation under either condition, with SEAP activity in cognate receiver cells 14.4-fold higher than in non-cognate receiver cells under activation conditions. These results demonstrate that mammalian cell activation depends on both sequence-specific ligand release from the proteinosomes and cognate ligand–receptor recognition, providing orthogonal control at the sender and receiver levels.

To verify ligand release during proteinosome–cell communication, we analysed culture supernatants from cognate CC–GEMS co-incubations by anti-SUMO Western blotting (Fig. 3f; Methods). The signalling ligand was detected following co-incubation in activation medium, but not in protection medium. Although the ligand band was less intense than that of the free-ligand control, heterogeneity in proteinosome size and ligand loading introduced uncertainty into the estimated ligand dose (Fig. 1d; Fig. S13). Nevertheless, receiver-cell activation remained comparable to the free-ligand condition, consistent with previous observations that CC–GEMS receptors can exhibit similar signalling responses across a range of ligand concentrations.^[33]^ Together, these results establish DNA-programmable communication between proteinosome-based synthetic cells and receptor-engineered mammalian cells, with DNA inputs controlling ligand release from the sender and ligand–receptor recognition determining activation of the receiver.

### Distributed DNA-Programmable Proteinosome Populations Enable AND-Gate Control of Mammalian Cell Signalling

Having established DNA-programmable proteinosome–mammalian cell communication using a single signalling ligand, we next extended the system to distributed Boolean AND logic (Fig. 4a). We designed two complementary ligand components, Cy5-labelled **ligand**_**AND1**_ containing A′Γ CC and Cy3-labelled **ligand**_**AND2**_ containing A′Γ’ CC, based on the previously reported AND-gate architecture of Makri Pistikou et al., in which association of complementary A’Γ and A’Γ’ coiled-coil components restored receptor-activating function.^[33]^ Both components were engineered with cysteine residues for cysteine-directed conjugation^[72]^ and coupled to orthogonal ssDNA domains, enabling independent localization and release from separate proteinosome populations. Construct design, protein expression and ssDNA conjugation were validated in Supplementary Figs. S14–S17 (Methods). Each cognate DNA input triggers TMSD-mediated release of its corresponding ligand component. Only when both DNA inputs are present can the released ligand components jointly support CC–GEMS activation, potentially through association in solution or at the receiver-cell surface.

**Figure 4.**
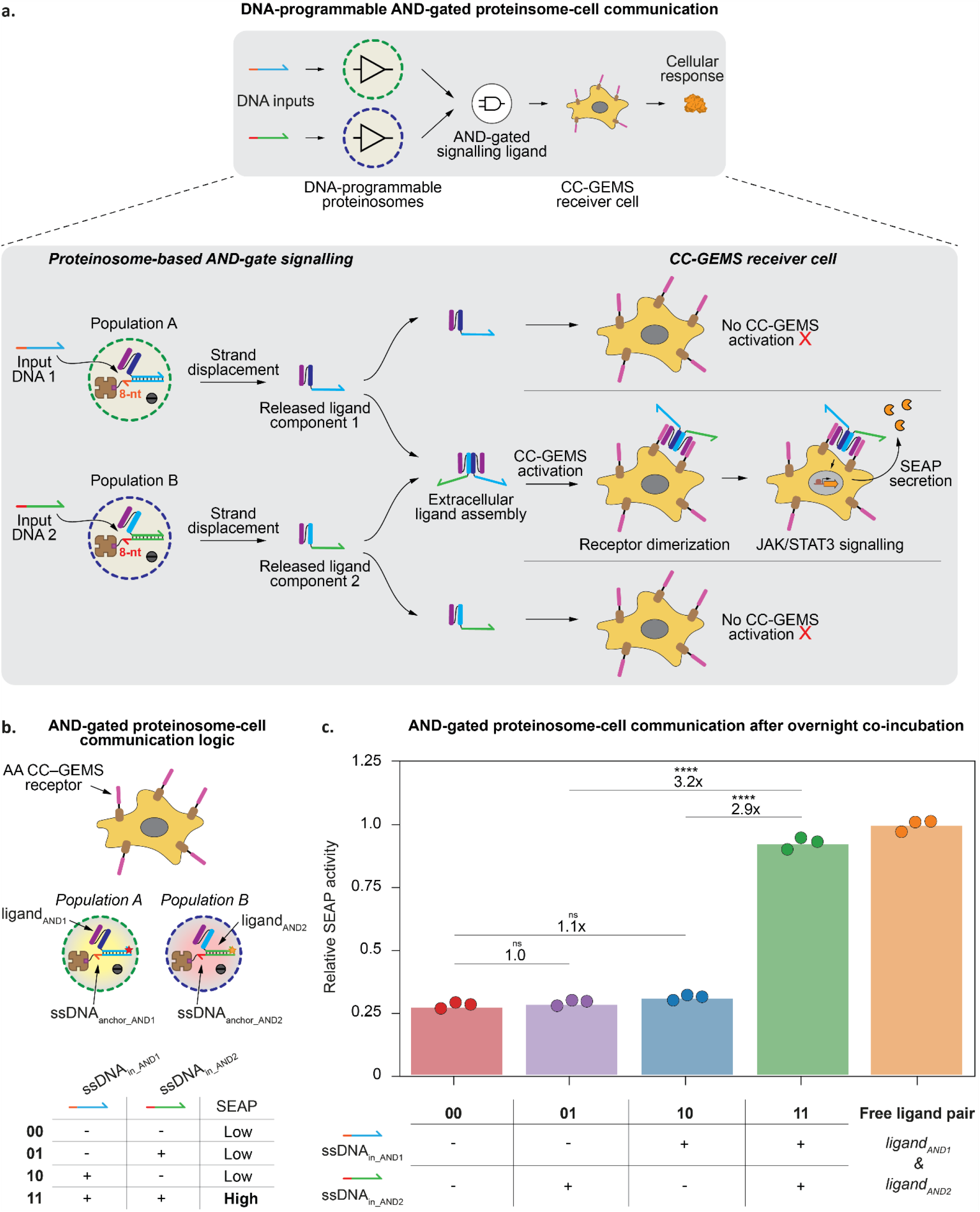
Distributed DNA-programmable proteinosome populations enable AND-gate control of mammalian cell signalling. (a) Schematic of DNA-programmable AND-gated proteinosome–cell communication. Proteinosome populations **A** and **B** contain complementary ligand components (**ligand**_**AND1**_ and **ligand**_**AND2**_) hybridized through 26-nt DNA domains to their respective biotinylated 34-nt anchors (**ssDNA**_**anchor_AND1**_ and **ssDNA**_**anchor_AND2**_), immobilized via streptavidin–biotin binding and leaving exposed 8-nt toeholds. Cognate 34-nt DNA inputs (**ssDNA**_**in_AND1**_ and **ssDNA**_**in_AND2**_) trigger TMSD-mediated release of the corresponding ligands. The released components associate through complementary coiled-coil interactions to form an active signalling ligand, which can engage neighbouring receiver cells and induce CC–GEMS receptor dimerization, JAK/STAT3 signalling and SEAP secretion. Both DNA inputs are required for formation of the active ligand. (b) Schematic of distributed AND-gate communication between two independently programmed proteinosome populations and CC–GEMS-expressing receiver cells. Four DNA-input states (00, 01, 10 and 11) produce low SEAP activity in the absence of one or both inputs and high activity when both inputs are present. (c) AA CC–GEMS-expressing receiver cells were co-incubated overnight with proteinosome populations **A** and **B.** Population **A** was FITC-labelled and contained Cy5-labelled **ligand**_**AND1**_; population **B** was DyLight405-labelled and contained Cy3-labelled **ligand**_**AND2**_. The four input conditions comprised neither input (00, 2 µM **ssDNA**_**scramble**_), input 1 alone (10, 1 µM **ssDNA**_**in_AND1**_), input 2 alone (01, 1 µM **ssDNA**_**in_AND2**_), or both inputs (11, 1 µM each). Scrambled DNA was added to the single-input conditions to maintain a total DNA concentration of 2 µM. A free-ligand positive control contained 120 nM each of **ligand**_**AND1**_ and **ligand**_**AND2**_ under scrambled-input conditions. SEAP activity was normalized to the free-ligand pair. The dual-input condition (11) elicited a response comparable to the free-ligand pair and was 3.2- and 2.9-fold higher than conditions 01 and 10, respectively. Neither single-input condition differed significantly from the no-input condition (01/00, 1.0-fold; 10/00, 1.1-fold). Data represent biological triplicates; bars indicate means and dots individual replicates. Statistical significance was determined by one-way ANOVA across all five conditions with Tukey’s multiple-comparison test. ns: P > 0.05; *P ≤ 0.05; **P ≤ 0.01; ***P ≤ 0.001; ****P ≤ 0.0001.

Before implementing distributed logic across proteinosome populations, we tested whether the complementary ssDNA–CC ligand components functioned as an AND gate independently of proteinosomes (Fig. S18). CC–GEMS-expressing HEK293T cells were treated overnight with neither ligand component (00), **ligand**_**AND1**_ alone (10), **ligand**_**AND2**_ alone (01), or both components together (11) (Methods). SEAP activity remained comparable under the 00, 01 and 10 conditions but increased substantially when both components were present (11). These results confirm that the DNA-conjugated ligand components retain AND-gate functionality, providing the basis for distributed proteinosome–mammalian cell communication.

We next examined whether the two ligand components could be independently localized and released from separate proteinosome populations in response to their respective DNA inputs (Fig. S19). Cy5-labelled **ligand**_**AND1**_ and Cy3-labelled **ligand**_**AND2**_ were localized through their corresponding ssDNA anchors within FITC-labelled proteinosome population **A** and DyLight405-labelled population **B**, respectively (Methods). Both ligands remained localized in the absence of input DNA (00). Addition of **ssDNA**_**in_AND1**_ (10) or **ssDNA**_**in_AND2**_ (01) selectively decreased the intraproteinosomal fluorescence of the corresponding ligand component, without detectable fluorescence loss in the non-cognate proteinosome population. These results support orthogonal, sequence-specific DNA-triggered release of the two ligand components, establishing independently programmable sender populations for distributed AND-gated communication.

To demonstrate distributed AND-gated communication, we co-incubated proteinosome populations **A** and **B** overnight with CC–GEMS-expressing mammalian cells (Fig. 4b,c). The four communication states comprised neither DNA input (00), **ssDNA**_**in_AND1**_ alone (10), **ssDNA**_**in_AND2**_ alone (01), or both inputs (11) (Fig. 4b; Methods). The single-input conditions (01 and 10) produced 1.0- and 1.1-fold the SEAP activity of the no-input condition (00), respectively, with neither differing significantly from 00 (Fig. 4c). In contrast, simultaneous addition of **ssDNA**_**in_AND1**_ and **ssDNA**_**in_AND2**_ increased SEAP activity 3.2-fold and 2.9-fold relative to conditions 01 and 10, respectively. The dual-input response was comparable to that elicited by the free ligand pair, consistent with extracellular assembly of an active signalling ligand from components released by the independently programmed proteinosome populations. Together, these results establish distributed AND-gated communication between synthetic-cell senders and mammalian receivers, in which both DNA inputs are required for substantial receptor activation.

Finally, we tested whether the single-input proteinosome–mammalian cell communication system could regulate an output beyond SEAP by measuring secretion of interleukin-10 (IL-10), an anti-inflammatory cytokine involved in immune regulation (Fig. S20; Methods). DNA-triggered ligand release increased IL-10 secretion relative to the non-triggered condition, demonstrating control over an alternative secreted protein. Together, these results establish distributed AND-gated communication and show that the single-input system can regulate distinct mammalian cell outputs.

## Conclusion

Here we establish **DIALECT**, a DNA-programmable communication interface linking bottom-up synthetic cells to receptor-engineered mammalian cells. Sequence-defined DNA inputs control TMSD-mediated release of coiled-coil signalling ligands from proteinosomes, coupling molecular processing within synthetic cells to CC–GEMS signalling and downstream gene expression in mammalian receiver cells. This architecture establishes orthogonal control at both the sender and receiver levels, with DNA sequence governing ligand release and cognate ligand–receptor recognition determining receiver-cell activation.

Importantly, we demonstrate that this communication architecture can be distributed across independently programmed synthetic-cell populations. By assigning complementary ligand components to two proteinosome populations responsive to distinct DNA inputs, we implemented Boolean AND logic in which substantial mammalian-cell activation requires both inputs. The single-input system controlled IL-10 secretion in addition to SEAP, demonstrating that the communication interface can regulate distinct mammalian outputs. These results establish bottom-up synthetic cells as programmable molecular processors whose outputs can engage genetically encoded signalling pathways in living cells, rather than serving solely as reservoirs of diffusible signals. Future incorporation of larger DNA reaction networks, additional orthogonal ligand–receptor pairs and more complex multicellular logic could enable increasingly sophisticated hybrid artificial–biological communication networks.

## Supporting Information

The authors have cited additional references within the Supporting Information.^[73]^

## Acknowledgements

This work was supported by the European Research Council (ERC; project no. 101000199 AMIGA) and the Dutch Research Council (NWO; project no. 016.176.622).

## Data Availability Statement

The datasets generated and analysed, and the code/scripts used for data analysis, in the present study have been deposited in the Zenodo database at https://zenodo.org/records/24011014 (https://doi.org/10.5281/zenodo.24011014).

## Entry for the Table of Contents

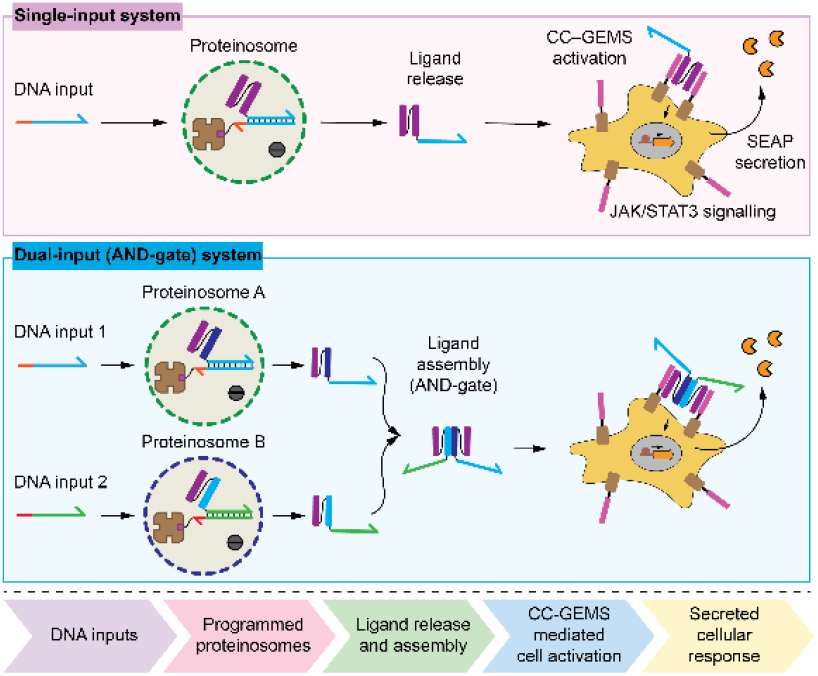

DNA-programmable proteinosomes enable synthetic cell–mammalian cell communication through engineered coiled-coil Generalized Extracellular Molecule Sensors (CC–GEMS). DNA-triggered ligand release and assembly provide single-input and distributed AND-gate control of mammalian signalling, enabling regulated secretion of specified outputs.

## References

[1] V. Mukwaya, S. Mann, and H. Dou, “Chemical communication at the synthetic cell/living cell interface,” Communications Chemistry 4 (2021): 161. 10.1038/s42004-021-00597-w.

[2] H. Moghimianavval, C. Newell, P. Parvizian, M. J. Booth, and A. P. Liu, “Strategies and applications of synthetic cell communication,” Nature Chemical Biology 21 (2025): 1317–1329. 10.1038/s41589-025-02002-2.

[3] H. Karoui, P. S. Patwal, B. V. V. S. Pavan Kumar, and N. Martin, “Chemical communication in artificial cells: Basic concepts, design and challenges,” Frontiers in Molecular Biosciences 9 (2022): 880525. 10.3389/fmolb.2022.880525.

[4] P. J. Grimes, A. Galanti, and P. Gobbo, “Bioinspired networks of communicating synthetic protocells,” Frontiers in Molecular Biosciences 8 (2021): 804717. 10.3389/fmolb.2021.804717.

[5] B. C. Buddingh’ and J. C. M. van Hest, “Artificial cells: Synthetic compartments with life-like functionality and adaptivity,” Accounts of Chemical Research 50 (2017): 769–777. 10.1021/acs.accounts.6b00512.

[6] Y. Elani, “Interfacing living and synthetic cells as an emerging frontier in synthetic biology,” Angewandte Chemie International Edition 60 (2021): 5602–5611. 10.1002/anie.202006941.

[7] K. P. Adamala, M. Dogterom, Y. Elani, et al., “The present and future of synthetic cell development,” Nature Reviews Molecular Cell Biology 25 (2024): 162–167. 10.1038/s41580-023-00686-9.

[8] A. Samanta, L. Baranda Pellejero, M. Masukawa, and A. Walther, “DNA-empowered synthetic cells as minimalistic life forms,” Nature Reviews Chemistry 8 (2024): 454–470. 10.1038/s41570-024-00606-1.

[9] C. E. Hilburger, M. L. Jacobs, K. R. Lewis, J. A. Peruzzi, and N. P. Kamat, “Controlling secretion in artificial cells with a membrane AND gate,” ACS Synthetic Biology 8 (2019): 1224–1230. 10.1021/acssynbio.8b00435.

[10] S. Yang, A. Joesaar, B. W. A. Bögels, S. Mann, and T. F. A. de Greef, “Protocellular CRISPR/Cas-Based Diffusive Communication Using Transcriptional RNA Signaling,” Angewandte Chemie International Edition 61 (2022): e202202436. 10.1002/anie.202202436.

[11] X. Huang, M. Li, D. C. Green, et al., “Interfacial assembly of protein–polymer nano-conjugates into stimulus-responsive biomimetic protocells,” Nature Communications 4 (2013): 2239. 10.1038/ncomms3239.

[12] A. Joesaar, S. Yang, B. Bögels, et al., “DNA-based communication in populations of synthetic protocells,” Nature Nanotechnology 14 (2019): 369–378. 10.1038/s41565-019-0399-9.

[13] S. Yang, P. A. Pieters, A. Joesaar, et al., “Light-Activated Signaling in DNA-Encoded Sender–Receiver Architectures,” ACS Nano 14 (2020): 15992–16002. 10.1021/acsnano.0c07537.

[14] A. Zambrano, G. Fracasso, M. Gao, et al., “Programmable synthetic cell networks regulated by tunable reaction rates,” Nature Communications 13 (2022): 3885. 10.1038/s41467-022-31471-5.

[15] A. Heidari, O. I. Sentürk, S. Yang, et al., “Orthogonal Light-Dependent Membrane Adhesion Induces Social Self-Sorting and Member-Specific DNA Communication in Synthetic Cell Communities,” Small 19 (2023): 2206474. 10.1002/smll.202206474.

[16] R. Lentini, S. P. Santero, F. Chizzolini, et al., “Integrating artificial with natural cells to translate chemical messages that direct Escherichia coli behaviour,” Nature Communications 5 (2014): 4012. 10.1038/ncomms5012.

[17] R. Lentini, N. Y. Martín, M. Forlin, et al., “Two-way chemical communication between artificial and natural cells,” ACS Central Science 3 (2017): 117–123. 10.1021/acscentsci.6b00330.

[18] H. Niederholtmeyer, C. Chaggan, and N. K. Devaraj, “Communication and quorum sensing in non-living mimics of eukaryotic cells,” Nature Communications 9 (2018): 5027. 10.1038/s41467-018-07473-7.

[19] I. Gispert, J. W. Hindley, C. P. Pilkington, et al., “Stimuli-responsive vesicles as distributed artificial organelles for bacterial activation,” Proceedings of the National Academy of Sciences of the United States of America 119 (2022): e2206563119. 10.1073/pnas.2206563119.

[20] J. M. Smith, D. Hartmann, and M. J. Booth, “Engineering cellular communication between light-activated synthetic cells and bacteria,” Nature Chemical Biology 19 (2023): 1138–1146. 10.1038/s41589-023-01374-7.

[21] X. Wang, L. Tian, H. Du, et al., “Chemical communication in spatially organized protocell colonies and protocell/living-cell microarrays,” Chemical Science 10 (2019): 9446–9453. 10.1039/c9sc04522h.

[22] Ö. D. Toparlak, J. Zasso, S. Bridi, et al., “Artificial cells drive neural differentiation,” Science Advances 6 (2020): eabb4920. 10.1126/sciadv.abb4920.

[23] G. Chen, R. Levin, S. Landau, et al., “Implanted synthetic cells trigger tissue angiogenesis through de novo production of recombinant growth factors,” Proceedings of the National Academy of Sciences of the United States of America 119 (2022): e2207525119. 10.1073/pnas.2207525119.

[24] Z. Chen, J. Wang, W. Sun, et al., “Synthetic beta cells for fusion-mediated dynamic insulin secretion,” Nature Chemical Biology 14 (2018): 86–93. 10.1038/nchembio.2511.

[25] A. B. Cook, S. Sun, Y. Li, J. Scheerstra, and J. C. M. van Hest, “Photoactivatable synthetic exosomes for RNA-based communication between artificial cells and living cells,” Angewandte Chemie International Edition 64 (2025): e202514041. 10.1002/anie.202514041.

[26] S. Sethi, C. Sharma, and A. Walther, “ATP-powered signaling between artificial and living cells,” Angewandte Chemie International Edition 64 (2025): e202517843. 10.1002/anie.202517843.

[27] M. Wang, H. Nan, M. Wang, et al., “Responsive DNA artificial cells for contact and behavior regulation of mammalian cells,” Nature Communications 16 (2025): 2410. 10.1038/s41467-025-57770-1.

[28] N. Krinsky, M. Kaduri, A. Zinger, et al., “Synthetic cells synthesize therapeutic proteins inside tumors,” Advanced Healthcare Materials 7 (2018): 1701163. 10.1002/adhm.201701163.

[29] Z. Liu, Y.-R. Li, Y. Yang, et al., “Viscoelastic synthetic antigen-presenting cells for augmenting the potency of cancer therapies,” Nature Biomedical Engineering 8 (2024): 1615–1633. 10.1038/s41551-024-01272-w.

[30] D. Burgstaller, N. Piernitzki, N. Küchler, et al., “Soft synthetic cells with mobile membrane ligands for ex vivo expansion of therapy-relevant T-cell phenotypes,” Small 20 (2024): e2401844. 10.1002/smll.202401844.

[31] D. K. Y. Zhang, A. S. Cheung, and D. J. Mooney, “Activation and expansion of human T cells using artificial antigen-presenting cell scaffolds,” Nature Protocols 15 (2020): 773–798. 10.1038/s41596-019-0249-0.

[32] L. Scheller, T. Strittmatter, D. Fuchs, D. Bojar, and M. Fussenegger, “Generalized extracellular molecule sensor platform for programming cellular behavior,” Nature Chemical Biology 14 (2018): 723–729. 10.1038/s41589-018-0046-z.

[33] A.-M. Makri Pistikou, G. A. O. Cremers, B. L. Nathalia, et al., “Engineering a scalable and orthogonal platform for synthetic communication in mammalian cells,” Nature Communications 14 (2023): 7001. 10.1038/s41467-023-42810-5.

[34] L. Morsut, K. T. Roybal, X. Xiong, et al., “Engineering customized cell sensing and response behaviors using synthetic Notch receptors,” Cell 164 (2016): 780–791. 10.1016/j.cell.2016.01.012.

[35] D. I. Piraner, M. H. Abedi, M. J. Duran Gonzalez, et al., “Engineered receptors for soluble cellular communication and disease sensing,” Nature 638 (2025): 805–813. 10.1038/s41586-024-08366-0.

[36] M. Garibyan, T. Hoffman, T. Makaske, et al., “Engineering programmable material-to-cell pathways via synthetic Notch receptors to spatially control differentiation in multicellular constructs,” Nature Communications 15 (2024): 5891. 10.1038/s41467-024-50126-1.

[37] Z.-R. Zhou, M.-S. Wu, Z. Yang, et al., “Synthetic transmembrane DNA receptors enable engineered sensing and actuation,” Nature Communications 16 (2025): 1464. 10.1038/s41467-025-56758-1.

[38] I. Nakase, S. Okumura, G. Tanaka, et al., “Signal transduction using an artificial receptor system that undergoes dimerization upon addition of a bivalent leucine-zipper ligand,” Angewandte Chemie International Edition 51 (2012): 7464–7467. 10.1002/anie.201201805.

[39] J. Manhas, H. I. Edelstein, J. N. Leonard, and L. Morsut, “The evolution of synthetic receptor systems,” Nature Chemical Biology 18 (2022): 244–255. 10.1038/s41589-021-00926-z.

[40] F. Teng, T. Cui, L. Zhou, et al., “Programmable synthetic receptors: The next generation of cell and gene therapies,” Signal Transduction and Targeted Therapy 9 (2024): 7. 10.1038/s41392-023-01680-5.

[41] Y. J. Chen, N. Dalchau, N. Srinivas, et al., “Programmable chemical controllers made from DNA,” Nature Nanotechnology 8 (2013): 755–762. 10.1038/nnano.2013.189.

[42] L. Qian, E. Winfree, “A simple DNA gate motif for synthesizing large-scale circuits,” Journal of the Royal Society Interface 8 (2011): 1281–1297. 10.1098/rsif.2010.0729.

[43] G. Seelig, D. Soloveichik, D. Y. Zhang, et al., “Enzyme-free nucleic acid logic circuits,” Science 314 (2006): 1585–1588. 10.1126/science.1132493.

[44] H. Gradišar, R. Jerala, “De novo design of orthogonal peptide pairs forming parallel coiled-coil heterodimers,” Journal of Peptide Science 17 (2011): 100–106. 10.1002/psc.1331.

[45] T. Plaper, E. Rihtar, T. Z. Ramuta, et al., “The art of designed coiled-coils for the regulation of mammalian cells,” Cell Chemical Biology 31 (2024): 1460–1472. 10.1016/j.chembiol.2024.06.001.

[46] T. Lebar, D. Lainšček, E. Merljak, et al., “A tunable orthogonal coiled-coil interaction toolbox for engineering mammalian cells,” Nature Chemical Biology 16 (2020): 513–519. 10.1038/s41589-019-0443-y.

[47] A. N. Lupas, M. Gruber, “The Structure of α-Helical Coiled Coils,” Advances in Protein Chemistry 70 (2005): 37–78. 10.1016/S0065-3233(05)70003-6.

[48] P. Burkhard, J. Stetefeld, S. V. Strelkov, “Coiled coils: a highly versatile protein folding motif,” Trends in Cell Biology 11 (2001): 82–88. 10.1016/S0962-8924(00)01898-5.

[49] Z. Chen, S. E. Boyken, M. Jia, et al., “Programmable design of orthogonal protein heterodimers,” Nature 565 (2019): 106–111. 10.1038/s41586-018-0802-y.

[50] D. Kashyap, M. J. Booth, “Nucleic Acid Conjugates: Unlocking Therapeutic Potential,” ACS Bio & Med Chem Au 5 (2025): 3–15. 10.1021/acsbiomedchemau.4c00092.

[51] B. A. R. Williams, J. C. Chaput, “Synthesis of peptide-oligonucleotide conjugates using a heterobifunctional crosslinker,” Current Protocols in Nucleic Acid Chemistry (2010). 10.1002/0471142700.nc0441s42.

[52] G. A. O. Cremers, B. J. H. M. Rosier, R. Riera Brillas, et al., “Efficient Small-Scale Conjugation of DNA to Primary Antibodies for Multiplexed Cellular Targeting,” Bioconjugate Chemistry 30 (2019): 2384–2392. 10.1021/acs.bioconjchem.9b00490.

[53] J. M. Chalker, G. J. L. Bernardes, Y. A. Lin, et al., “Chemical Modification of Proteins at Cysteine: Opportunities in Chemistry and Biology,” Chemistry – An Asian Journal 4 (2009): 630–640. 10.1002/asia.200800427.

[54] B. W. A. Bögels, B. H. Nguyen, D. Ward, et al., “DNA storage in thermoresponsive microcapsules for repeated random multiplexed data access,” Nature Nanotechnology 18 (2023): 912–921. 10.1038/s41565-023-01377-4.

[55] M. Gao, D. Wang, M. Wilsch-Bräuninger, et al., “Cell Free Expression in Proteinosomes Prepared from Native Protein–PNIPAAm Conjugates,” Macromolecular Bioscience 24 (2024): e2300464. 10.1002/mabi.202300464.

[56] A. Bocedi, G. Cattani, L. Stella, et al., “Thiol disulfide exchange reactions in human serum albumin: the apparent paradox of the redox transitions of Cys34,” FEBS Journal 285 (2018): 3225–3237. 10.1111/febs.14609.

[57] H. Khan, K. A. Khan, M. K. Khan, et al., “Thiol-disulfide exchange reactions occurring at modified bovine serum albumin detected using ellman’s reagent (5,5′-dithiobis (2-nitrobenzoic acid),” Pakistan Journal of Pharmaceutical Sciences 33 (2020): 2767–2772. 10.36721/PJPS.2020.33.6.SUP.2767-2772.1.

[58] J. Christodoulou, P. J. Sadler, A. Tucker, “A New Structural Transition of Serum Albumin Dependent on the State of Cys34: Detection by 1H-NMR Spectroscopy,” European Journal of Biochemistry 225 (1994): 363–368. 10.1111/j.1432-1033.1994.00363.x.

[59] D. Lysne, T. Hachigian, C. Thachuk, et al., “Leveraging Steric Moieties for Kinetic Control of DNA Strand Displacement Reactions,” Journal of the American Chemical Society 145 (2023): 16691–16703. 10.1021/jacs.3c04344.

[60] D. Yao, B. Wang, S. Xiao, et al., “What Controls the ‘Off/On Switch’ in the Toehold-Mediated Strand Displacement Reaction on DNA Conjugated Gold Nanoparticles?,” Langmuir 31 (2015): 7055–7061. 10.1021/acs.langmuir.5b01671.

[61] J. Hahn, S. F. J. Wickham, W. M. Shih, and S. D. Perrault, “Addressing the instability of DNA nanostructures in tissue culture,” ACS Nano 8 (2014): 8765–8775. 10.1021/nn503513p.

[62] S. Akhtar, R. Kole, and R. L. Juliano, “Stability of antisense DNA oligodeoxynucleotide analogs in cellular extracts and sera,” Life Sciences 49 (1991): 1793–1801. 10.1016/0024-3205(91)90480-Y.

[63] J. Fern and R. Schulman, “Design and characterization of DNA strand-displacement circuits in serum-supplemented cell medium,” ACS Synthetic Biology 6 (2017): 1774–1783. 10.1021/acssynbio.7b00105.

[64] H. Zhang, J. Vandesompele, K. Braeckmans, et al., “Nucleic acid degradation as barrier to gene delivery: a guide to understand and overcome nuclease activity,” Chemical Society Reviews 53 (2024): 317–360. 10.1039/D3CS00194F.

[65] A. R. Chandrasekaran, “Nuclease resistance of DNA nanostructures,” Nature Reviews Chemistry 5 (2021): 225–239. 10.1038/s41570-021-00251-y.

[66] H. Zhang, S. H. Kim, and I. T. S. Li, “Decoy DNA protects molecular tension probes from DNase degradation,” Angewandte Chemie International Edition 64 (2025): e202506590. 10.1002/anie.202506590.

[67] U. Piran and W. J. Riordan, “Dissociation rate constant of the biotin-streptavidin complex,” Journal of Immunological Methods 133 (1990): 141–143. 10.1016/0022-1759(90)90328-S.

[68] P. Tiwary, “Molecular Determinants and Bottlenecks in the Dissociation Dynamics of Biotin–Streptavidin,” The Journal of Physical Chemistry B 121 (2017): 10841–10849. 10.1021/acs.jpcb.7b09510.

[69] L. Deng, E. N. Kitova, and J. S. Klassen, “Dissociation kinetics of the streptavidin–biotin interaction measured using direct electrospray ionization mass spectrometry analysis,” Journal of the American Society for Mass Spectrometry 24 (2013): 49–56. 10.1007/s13361-012-0533-5.

[70] R. F. Delgadillo, T. C. Mueser, K. Zaleta-Rivera, et al., “Detailed characterization of the solution kinetics and thermodynamics of biotin, biocytin and HABA binding to avidin and streptavidin,” PLOS ONE 14 (2019): e0204194. 10.1371/journal.pone.0204194.

[71] C. E. Chivers, E. Crozat, C. Chu, et al., “A streptavidin variant with slower biotin dissociation and increased mechanostability,” Nature Methods 7 (2010): 391–393. 10.1038/nmeth.1450.

[72] T. W. van Veldhuisen, L. M. V. Raeven, N. G. L. van Herwijnen, et al., “Engineering a Transmembrane Receptor for Coacervate-Based Artificial Cells,” Journal of the American Chemical Society 148 (2026): 24926–24934. 10.1021/jacs.6c03433.

[73] M. A. Unger, H. P. Chou, T. Thorsen, A. Scherer, and S. R. Quake, “Monolithic microfabricated valves and pumps by multilayer soft lithography,” Science 288 (2000): 113–116. 10.1126/science.288.5463.113.

