## Supplementary Information for "DIALECT: DNA-Programmable Synthetic Cells Communicate with Mammalian Cells through Synthetic Receptors"

*Supporting information*

#### Table of contents


- Supplementary Methods
- Supplementary Movies
- Supplementary Figures
    - Figure S1. Design of cysteine-engineered **CC**$_{\textbf{ditopic}}$
    - Figure S2. Expression and purification of **CC**$_{\textbf{ditopic}}$
    - Figure S3. Purification of ssDNA–CC signalling ligand
    - Figure S4. Cy3 fluorescence calibration
    - Figure S5. Nonspecific association of signalling components
    - Figure S6. Two-layer microfluidic trapping array
    - Figure S7. Microfluidic monitoring of DNA-triggered ligand release
    - Figure S8. Intraproteinosomal ligand retention in culture media
    - Figure S9. Stability of proteinosome localization components
    - Figure S10. ssDNA–CC ligand integrity during receptor activation
    - Figure S11. DNA-programmable ligand release during cell co-incubation
    - Figure S12. Proteinosome–mammalian cell compatibility
    - Figure S13. Proteinosome sender population estimation
    - Figure S14. Design of cysteine-engineered **CC**$_{\textbf{AND1}}$
    - Figure S15. Design of cysteine-engineered **CC**$_{\textbf{AND2}}$
    - Figure S16. Expression and purification of **CC**$_{\textbf{AND1}}$ and **CC**$_{\textbf{AND2}}$
    - Figure S17. Conjugation and purification of AND-gate ligands
    - Figure S18. CC–GEMS activation by complementary ligand components
    - Figure S19. Orthogonal localization and release of AND-gate ligands
    - Figure S20. DNA-programmable communication regulates IL-10 secretion
    - Figure S21. SEAP calibration curve
    - Figure S22. Uncropped SDS–PAGE gels
    - Figure S23. Uncropped Native-PAGE gel
    - Figure S24. Uncropped Western blot membranes
- Supplementary Tables
    - Table S1. Plasmid constructs
    - Table S2. CC–GEMS receptor design and sequences
    - Table S3. DNA oligonucleotides
    - Table S4. CC proteins
    - Table S5. ssDNA–CC ligand constructs
    - Table S6. Fluorescence intensity and fold-change summary
    - Table S7. Unpaired two-tailed t-test summary
    - Table S8. One-way ANOVA and Tukey multiple-comparisons summary
- Supplementary References

#### Supplementary Methods

##### Materials

Plasmids encoding the CC–GEMS receptor constructs and associated expression and reporter components, together with bacterial expression plasmids encoding the original coiled-coil (CC) proteins, were obtained from Pistikou, et al.[33], whereas bacterial expression plasmids encoding the AND-gated CC proteins were obtained from Veldhuisen, et al.[72] DNA oligonucleotides were purchased from Integrated DNA Technologies (IDT). Modified oligonucleotides were purified by high-performance liquid chromatography (HPLC), whereas unmodified oligonucleotides were supplied desalted. Oligonucleotide stock solutions (1,000 or 100 µM) were prepared in nuclease-free water (Invitrogen, AM993) and stored at −30 °C. All plasmid constructs, DNA oligonucleotides, CC proteins and ssDNA–CC ligand constructs used in this study are listed in Supplementary Tables S1–S5. Unless otherwise stated, reagents and solvents were obtained from commercial suppliers and used without further purification.

##### Mammalian cell culture

HEK293T cells (ATCC, CRL3216) were cultured in standard culture medium consisting of Dulbecco's modified Eagle's medium (DMEM; Gibco, 41966) supplemented with 10% fetal bovine serum (FBS, Greiner Bio-One, 758093) and 1% (v/v) penicillin–streptomycin (10,000 U $mL^{-1}$; Gibco, 15140). Standard culture conditions consisted of incubation at 37 °C and 5% $CO_2$ in a humidified incubator. Cells were monitored routinely, passaged to maintain exponential growth and used up to a passage number of 30. Cell-line authentication and mycoplasma testing were performed by the supplier before distribution.

##### Expression and purification of cysteine-engineered coiled-coil proteins

A pET28a(+) vector encoding the cysteine-engineered coiled-coils (Fig. S1; Fig. S14; Fig. S15) was transformed into *Escherichia coli* BL21(DE3) (*E. coli,* Novagen, 69450). A single colony was cultured in a shaking incubator at 140 rpm and 37 °C in 1 L of LB medium, supplemented with 50 µg $mL^{-1}$ kanamycin (Carl Roth, T832). When the $OD_{600}$ (optical density measured at 600 nm) of the culture reached ~0.6–0.8, protein expression was induced by addition of β-D-1-thiogalactopyranoside (IPTG, AppliChem, A1008) to a final concentration of 100 µM, and the culture was incubated overnight at 140 rpm and 16 °C. Cells were harvested by centrifugation at 8,000× g for 10 min at 4 °C. Pelleted cells were resuspended in 13.33 mL BugBuster (Merck, 70584) supplemented with 5 µL benzonase (Merck, E1014) and one protease-inhibitor tablet (Thermo Scientific, A32965) and incubated for 1 h at room temperature on a shaking plate. The lysate was centrifuged at 40,000× g for 20 min at 4 °C and the supernatant containing soluble cysteine-engineered coiled coil (CC) proteins was collected for purification.
Protein purification was performed by $Ni^{2+}$-affinity chromatography using Ni-NTA resin (QIAGEN, 30230) on a gravity column with a column volume of 2 mL. The column was equilibrated with binding buffer (20 mM Tris-Cl, pH 7.9, 500 mM NaCl, 5 mM imidazole), after which the lysate was loaded onto the column. The column was subsequently washed with binding buffer followed by washing buffer (20 mM Tris-Cl, pH 7.9, 500 mM NaCl, 20 mM imidazole). The cysteine-engineered coiled coils were eluted using elution buffer (20 mM Tris-Cl, pH 7.9, 500 mM NaCl and 100 mM imidazole), and eluted protein fractions were dialysed using SnakeSkin Dialysis Tubing (3.5 kDa MWCO, Thermo Scientific, 68035) overnight at 4 °C against 2 L of dialysis buffer (100 mM Tris, 150 mM NaCl, pH 8.0) to remove imidazole. Following dialysis, protein samples were concentrated using an Amicon Ultra Centrifugal Filter (10 kDa MWCO, Millipore, UFC9010), snap-frozen and stored at −80 °C until further use. Protein concentration was determined by spectrophotometry by measuring absorption at 280 nm (NanoDrop One, Thermo Scientific), assuming an extinction coefficient calculated using the ProtParam tool on the ExPASy server. Protein purity was assessed by Coomassie-stained SDS–PAGE (Fig. S2; Fig. S16). Since the CC proteins were used for synthesis of ssDNA–CC ligands, they were stored with a final concentration of 2 mM TCEP (tris(2-carboxyethyl)phosphine), Merck, C4706) reducing agent added to the protein aliquots to prevent disulfide formation.

##### DNA–coiled-coil ligand conjugation and purification

Amine-modified ssDNA was functionalised with Sulfo-SMCC before conjugation to cysteine-engineered coiled-coil (CC) proteins (Fig. 1b). In a typical reaction, 30 µL of 125 µM amine-modified ssDNA (3.75 nmol) was mixed with 90 µL of PBS (pH 7.2) and 120 µL of 1,250 µM Sulfo-SMCC (Thermo Scientific, A39268) in DMSO. The reaction mixture was incubated at 850 rpm for 2 h at 20 °C. Excess Sulfo-SMCC was removed using two rounds of ethanol precipitation. SMCC-functionalized ssDNA was precipitated by adding 10% (v/v) 5 M NaCl and 300% (v/v) ice-cold ethanol, followed by incubation for 60 min at −30 °C. The reaction mixture was centrifuged at 14,000 rpm for 30 min at 4 °C, and the pellet was reconstituted in PBS (pH 7.2) before the precipitation was repeated. The final pellet was washed with 200 µL of 95% (v/v) ice-cold ethanol, centrifuged at 14,000 rpm for 15 min, and air-dried. SMCC-functionalized ssDNA was recovered at approximately 95% yield.
For the conjugation of cysteine-engineered coiled coil (CC) proteins to SMCC-functionalized ssDNA (Fig. 1b), an aliquot of CC protein was buffer-exchanged to 100 mM sodium phosphate (pH 7.0) containing 25 µM TCEP by repeated washing and centrifugation using an Amicon 10 kDa MWCO centrifugal filter (Merck Millipore, UFC5010). The desalted CC protein was subsequently concentrated to a final concentration of 25 µM using an Amicon 10 kDa MWCO centrifugal filter. SMCC-functionalized ssDNA (~3.56 nmol) was reconstituted directly in 50 µL of CC protein solution (25 µM, 1.25 nmol), corresponding to a DNA-to-CC molar ratio of approximately 2.85:1. The reaction mixture was shaken at 550 rpm for 1 h at 20 °C. The coupling efficiency was assessed using Coomassie-stained SDS–PAGE (Fig. 1b; Fig. S17).
The purification of ssDNA–CC ligand was performed using $Ni^{2+}$-affinity chromatography using Ni-NTA resin (QIAGEN, 30230) in a centrifugal filter (Merck Millipore, UFC30GV) with a column volume of 100 µL. The column was spun for 1 min at 500× g to remove residual fluid of the resin. The column was equilibrated with 500 µL of binding buffer (20 mM Tris-Cl, pH 7.9, 500 mM

NaCl, 5 mM imidazole), after which the column was centrifuged for 2 min at 500× g. The ssDNA–CC sample was prepared as a 500 µL sample by addition of binding buffer to the reaction mixture, loaded onto the column and incubated for 60 min with light agitation. The column was subsequently spun at 500 × *g* for 2 min to remove the unbound ssDNA. The column was subsequently washed with 500 µL of washing buffer (20 mM Tris-Cl, pH 7.9, 500 mM NaCl, 20 mM imidazole) and incubated for 2 min with gentle agitation, after which the column was spun for 2 min at 500 × *g*. The washing step was repeated three more times, after which the ssDNA–CC ligand was eluted by addition of 250 µL of elution buffer (20 mM Tris-Cl, pH 7.9, 500 mM NaCl, 100 mM imidazole) and incubation for 2 min with gentle agitation, after which the column was spun for 2 min at 500 × *g* to recover the purified protein. The elution step was repeated one more time and the pooled eluates were washed by Amicon 10 kDa MWCO centrifugal filter (Merck Millipore, UFC5010) to PBS to remove imidazole. Following washing, ssDNA–CC samples were concentrated using an Amicon centrifugal filter, snap-frozen and stored at −20 °C until further use. Ligand concentration was determined by spectrophotometry by measuring absorption at 260 nm (NanoDrop One, Thermo Scientific) using the oligonucleotide extinction coefficient supplied by Integrated DNA Technologies (IDT). Ligand purity was assessed by SYBR-gold-stained native-PAGE (Fig. S3).

For ligand with Cy3-labelled CC protein, the same procedure was executed except that labelled CC was used as the starting material. To prepare Cy3-labelled ligand with fluorescently labelled cysteine-engineered CC protein, the CC was labelled with Cy3 as follows: to 30 µM CC, 300 µM of sulfo-Cyanine 3 NHS ester (Lumiprobe, 11320) was added from a 10 mM stock dissolved in DMSO. The solution was incubated at 850 rpm for 2 h at 20 °C, and buffer-exchanged to 100 mM sodium phosphate (pH 7.0) with 25 µM TCEP using repetitive washing and centrifugation with an Amicon 10 kDa MWCO centrifugal filter (Merck Millipore, UFC5010) to remove residual unconjugated dye. The resulting Cy3-labelled CC was concentrated to a final concentration of 25 µM using an Amicon 10 kDa MWCO centrifugal filter and immediately used for conjugation to SMCC-functionalized ssDNA. The degree of labelling was determined spectrophotometrically from Cy3 absorbance at 548 nm, yielding a DoL of 4.5 Cy3 molecules per CC molecule.

**Proteinosome fabrication**

Cationized bovine serum albumin (BSA–$NH_2$) was synthesized using a previously reported method.[11] Diaminohexane (1.5 g, 12.9 mmol) was dissolved in 10 mL Milli-Q water and adjusted to pH 6.5 with 5 M HCl. The solution was added dropwise to BSA (200 mg, 3 µmol) dissolved in 10 mL Milli-Q water. 1-(3-Dimethylaminopropyl)-3-ethylcarbodiimide (EDC, Thermo Scientific, A10807; 100 mg) was added immediately, followed by a further 50 mg after 5 h. The pH was readjusted to 6.5 when necessary, and the reaction was stirred for a further 6 h. Precipitated material was removed by centrifugation and the supernatant was dialysed overnight against Milli-Q water using dialysis tubing with a 12–14 kDa MWCO (Medicell, DTV.12000) and then freeze-dried. End-capped mercaptothiazoline-activated PNIPAm ($M_n$ = 8,500 g mol⁻¹; 4 mg in 5 mL Milli-Q water), prepared as described by X. Huang, et al.[11], was added to BSA–$NH_2$ (10 mg in 5 mL PBS, pH 8.0). The mixture was stirred for 10 h. The resulting conjugates were purified using a 50 kDa MWCO centrifugal filter (Merck Millipore, UFC9050) and freeze-dried.

For FITC labelling, BSA (200 mg) was dissolved in 10 mL of 50 mM sodium carbonate buffer, pH 9.0. Fluorescein isothiocyanate (FITC, Merck, F7250; 2.36 mg in 590 µL DMSO) was added and the mixture was stirred for 5 h. The labelled BSA was dialysed overnight against Milli-Q water using 12–14 kDa MWCO dialysis tubing and freeze-dried. For DyLight405 labelling, BSA (30 mg) was dissolved in 6 mL of 50 mM sodium carbonate buffer, pH 9.0. DyLight405 NHS ester (Thermo Scientific, 46400; 1 mg in 100 µL DMF) was added, and the reaction was stirred for 2 h. The labelled BSA was dialysed overnight against Milli-Q water and freeze-dried. Fluorescently labelled BSA–$NH_2$/PNIPAm conjugates were prepared using the corresponding labelled BSA as the starting material.

Proteinosomes were assembled at a standard batch size of 15 µL using a previously reported method.[54] BSA–$NH_2$/PNIPAm conjugates, streptavidin and magnetic beads were combined in 7.5 µL of aqueous solution. Separately, PEG-bis(N-succinimidyl succinate) (2 mg; Merck, 713783) was dissolved in 7.5 µL of 50 mM sodium carbonate buffer (pH 9.0) and added to the mixture. The conjugates were included at a total concentration of 8 mg $mL^{-1}$, comprising 7 mg $mL^{-1}$ unlabelled conjugate and 1 mg $mL^{-1}$ fluorescently labelled conjugate, prepared from stock solutions of 32 mg $mL^{-1}$ and 16 mg $mL^{-1}$, respectively. Streptavidin (100 µM stock; Carl Roth, 6073) and Dynabeads MyOne Carboxylic Acid (10 mg $mL^{-1}$ stock; Thermo Scientific, 10563775) were included at final concentrations of 11.88 µM and 1 mg $mL^{-1}$, respectively. At a final aqueous-phase volume of 15 µL, the amounts corresponded to 120 µg total conjugate, including 15 µg fluorescently labelled conjugate, 178.2 pmol streptavidin and 15 µg magnetic beads. For the microfluidic strand-displacement release experiment (Fig. 1e–f; Fig. S7), Dynabeads M-270 Amine (30 mg $mL^{-1}$ stock; Thermo Scientific, 10151003) were used instead, at a final concentration of 3 mg $mL^{-1}$, corresponding to 45 µg of magnetic beads per 15 µL aqueous phase. Batch size was adjusted for proteinosome–cell co-incubation experiments by scaling all reagent quantities proportionally to the final aqueous-phase volume while maintaining the same final concentrations and reagent-to-volume ratios. After resuspension, 300 µL of 2-ethyl-1-hexanol was added, and the mixture was shaken vigorously by hand for 30 s to form a Pickering emulsion.

The emulsion was left at room temperature for 2 h to allow crosslinking and sedimentation. Transfer to the aqueous phase was performed using a washing procedure adapted from Gao et al.[55] The upper oil phase was removed and 400 µL ice-cold 75% ethanol was added. After resuspension, the sample was incubated at 4 °C for 2–3 h. Proteinosomes were collected by centrifugation at 3,000 × *g* for 5 min at room temperature and resuspended in 400 µL of ice-cold 50% ethanol. The suspension was centrifuged under the same conditions, followed by two washes with Milli-Q water. No additional incubation was performed during these subsequent washing steps. Proteinosomes were stored at 4 °C. Bright-field imaging using an EVOS XL microscope (Thermo Scientific) was used to visualize proteinosome membranes. Fluorescence imaging (Nikon, Eclipse Ti2-D-PD) was used to visualize labelled BSA–$NH_2$/PNIPAm conjugates. These imaging modes were used to assess proteinosome dimensions, morphology and abundance.

**DNA-programmable ligand localization and transfer between media**

Localization and wash buffer consisted of 10 mM Tris (pH 8.0; 1 M stock, Invitrogen), 3 mM $MgCl_2$ (1 M stock, Invitrogen), 430 mM NaCl (5 M stock, Invitrogen) and 0.1% (v/v) Tween 20 (Sigma). Proteinosome recovery during buffer exchange and washing was based on a previously reported method.[54] For standard batches, 15 µL of proteinosome suspension was resuspended in 500 µL of buffer or medium, after which the proteinosomes were recovered using a magnetic rack for 10–15 min and the supernatant was removed by pipetting. For larger batches, centrifugation at 3,000 × *g* for 1 min at room temperature was performed before magnetic separation to facilitate proteinosome recovery.

Streptavidin-containing proteinosomes were transferred from their initial aqueous suspension into buffer by two successive buffer exchanges. After removal of the supernatant, the proteinosomes were resuspended in 13.5 µL of buffer, and 1.5 µL of tenfold-diluted biotinylated ssDNA (10 µM working solution prepared from a 100 µM stock) was added to obtain a final concentration of 1 µM in a total volume of 15 µL. Samples were incubated at room temperature for 1 h to allow binding to streptavidin (Fig. 1c; Fig. S9). Unbound DNA was removed by resuspending the proteinosomes in 500 µL of buffer and allowing them to stand for 1 h before recovery. After removal of the supernatant, the proteinosomes were resuspended in 13.5 µL of buffer, and 1.5 µL of complementary ssDNA (10 µM working solution prepared by tenfold dilution of 100 µM stock Fig. S5; Fig. S9) or ssDNA–CC ligand (10 µM stock; Fig. 1c) was added to obtain a final concentration of 1 µM in a total volume of 15 µL. Samples were incubated overnight at 4 °C to allow DNA hybridization. Unbound DNA or ligand was removed by resuspending the proteinosomes in 500 µL of buffer and allowing them to stand for 3 h at 4 °C before recovery. Two additional washes were performed by immediate resuspension and recovery without further incubation. DNA- or ligand-loaded proteinosomes were stored at 4 °C until further use.

For experiments in culture medium, ligand-localized proteinosomes were transferred from buffer into the specified culture medium by sequential buffer exchange. The buffer was gradually replaced with culture medium using intermediate buffer-to-medium ratios of 75:25, 50:50 and 25:75 (v/v), followed by two washes in 100% culture medium. Proteinosomes were recovered after each exchange or wash as described above.

**Stability of ssDNA–CC ligand in cell-compatible media**

The stability of the ssDNA–CC signalling ligand ($\textbf{ligand}_{\textbf{ssDNA–CC}}$; Fig. 2a-b) was assessed following overnight incubation at 37 °C at a final concentration of 1 µM in the following media: standard culture medium; DMEM containing 10%, 5%, 2.5% or 1.25% heat-inactivated FBS (HI-FBS); serum-free DMEM; serum-free 293 SFM II medium (SFM, Gibco, 10503283); or supplemented DMEM containing 1.25% HI-FBS. Supplemented conditions included 1 µM non-complementary ssDNA ($\textbf{ssDNA}_{\textbf{scramble}}$), 50% (v/v) PBS, or both supplements. Stability medium consisted of DMEM and PBS mixed at a 1:1 (v/v) ratio, with a final HI-FBS concentration of 1.25% (v/v). To prepare 1 mL, 500 µL PBS, 487.5 µL DMEM and 12.5 µL HI-FBS were combined. Protection medium had the same final composition, supplemented with 10 µL of a 100 µM non-complementary ssDNA ($\textbf{ssDNA}_{\textbf{scramble}}$) stock solution to give a final concentration of 1 µM. HI-FBS was heat-inactivated at 56 °C for 30 min and stored at −20 °C until use. Both media were freshly prepared immediately before use. Following incubation, samples were diluted tenfold in PBS to a final ligand concentration of 100 nM to reduce interference from serum components during electrophoresis. Ligand integrity was assessed by SDS–PAGE followed by anti-SUMO Western blotting.

**SDS–PAGE and native-PAGE analysis**

Unless otherwise specified, samples for SDS–PAGE were prepared under reducing, non-heated conditions by mixing 6 µL of sample with 6 µL of 2× Laemmli sample buffer (Bio-Rad, 1610737) and adding 0.5 µL of 500 mM TCEP stock solution to achieve a final TCEP concentration of 20 mM. Where indicated, samples were heated at 95 °C for 5 min before loading (Fig. S2; Fig. S16). Samples were separated on precast 4–20% gradient polyacrylamide gels (Bio-Rad, 4561096) at 120 V for 60 min in SDS–PAGE running buffer (Tris/Glycine/SDS, Bio-Rad, 1610732). A total of 10 µL was loaded per lane.

For native-PAGE (Fig. S3), 8% polyacrylamide gels were prepared using standard gel-casting procedures. Samples were prepared under non-reducing, non-heated conditions in 10% glycerol and separated at 120 V for 60 min in 1× TAE running buffer (Tris/Acetic Acid/EDTA, Bio-Rad, 1610742). Following electrophoresis, gels were stained with SYBR Gold (Invitrogen, 10358492) for DNA detection or Coomassie Brilliant Blue (R-250, Bio-Rad, 1610436) for protein detection. Gels were imaged using an ImageQuant 800 system (Cytiva).

**Anti-SUMO Western blotting and densitometry**

Samples were separated by reducing, non-heated SDS–PAGE and transferred to a PVDF membrane using an iBlot 2 dry-transfer system and iBlot 2 Transfer Stacks (Thermo Scientific, IB21001 and IB24001), according to the manufacturer's instructions. Membranes were blocked for 1 h in 5% non-fat milk in TBS-T (0.1% Tween 20 in buffer containing 8 g NaCl, 0.2 g KCl and 3 g Tris base per litre, pH 7.4), then incubated overnight at 4 °C with anti-Smt3 rabbit antibody (Abcam, ab14405; 1:1,000) in blocking buffer. After three 10-min washes in TBS-T at room temperature, membranes were incubated for 1 h at room temperature with HRP-conjugated anti-rabbit IgG (Cell Signalling Technology, #7074; 1:5,000). Membranes were then washed three times for 10 min each. Chemiluminescence was developed using SuperSignal West Pico PLUS substrate (Thermo Scientific, 15659364) and detected with an ImageQuant 800 imager (Cytiva) (Fig. 2b; Fig. 3f; Fig. S10). Band densitometry was performed in ImageJ using the gel-analysis tool. Background-subtracted band densities were calculated from the area under the intensity profile. The upper band was assigned to intact DNA–CC conjugate and the lower band to a species consistent with DNA degradation. Percentage intact ligand was calculated as intact-band intensity divided by the sum of intact- and DNA-degraded-band intensities, multiplied by 100. Additional bands considered nonspecific were excluded from the calculation. Data are presented as a heat map showing the relative abundance of intact ssDNA–CC conjugate across samples (Fig. 2b; Fig. S10).

**Fluorescence microscopy and image analysis**

Fluorescence microscopy was performed using a high-content screening fluorescence microscope (Nikon, Eclipse Ti2-D-PD) equipped with solid-state lasers at 390 nm (DyLight405), 475 nm (FITC), 575 nm (Cy3/Alexa Fluor 546) and 637 nm (Cy5), with a 10× objective (NA 0.30) and a Prime 95B camera. Images were acquired at 12-bit depth, with acquisition settings held constant within each fluorescence channel across conditions and time points to enable quantitative comparisons. Images acquired before and after incubation represent different fields of view.

For suspension imaging, 2 µL of proteinosome suspension was placed on a microscope slide and allowed to settle for 1–2 min before image acquisition. Proteinosomes were identified using the FITC channel, and regions of interest (ROIs) were placed around individually resolved, approximately round objects. Debris, overlapping objects and proteinosomes intersecting image boundaries were excluded. Up to 30 resolvable proteinosomes were analysed per condition. For proteinosome–cell co-incubation experiments, images were acquired from 96-well plates (Greiner) (Fig. 3b; Fig. S11; Fig. S12). Because the wells contained dense proteinosome populations, fluorescence was quantified using a grid-based image-sampling approach rather than individual proteinosome segmentation. A total of 500 ROIs corresponding to image-sampling regions were analysed per biological replicate, with three independent biological replicates per condition (Fig. 3b; Fig. S11). For the DNA-programmable AND-gate experiments (Fig. S19), population **A** was identified using FITC fluorescence and population **B** using DyLight405 fluorescence. No upper limit of 30 proteinosomes per condition was imposed; all eligible proteinosomes identified in the respective fluorescence channels were quantified. For all experimental formats, corresponding ROIs were transferred to the relevant fluorescence channel, and mean fluorescence intensity was quantified using custom Python code. Mean fluorescence intensity was used for all subsequent quantitative analyses and data visualisations. Unless otherwise specified, no background subtraction was applied to the fluorescence measurements.

**Cy3 calibration and quantification of localized ligand**

A calibration curve was generated using known concentrations of Cy3-labelled ssDNA–CC ligand (**ligand$_{fluor}$**; Fig. S4). For fluorescence imaging, 2 µL droplets of each standard were placed on a microscope slide and imaged using a high-content screening fluorescence microscope (Nikon, Eclipse Ti2-D-PD). Three replicate measurements were acquired per concentration. For each calibration image, mean Cy3 fluorescence intensity was measured within a fixed 800 × 800-pixel region of interest (ROI) positioned in the central region of the droplet to avoid the droplet boundary. Mean Cy3 fluorescence intensity was plotted against ligand concentration to generate the calibration curve.

To quantify DNA-localized ligand, FITC-labelled proteinosomes containing **ligand$_{fluor}$** were imaged, and mean intraproteinosomal Cy3 fluorescence intensity was measured using the same intensity metric and acquisition settings as for the calibration standards. Ligand concentrations were estimated from the calibration curve (Fig. 1d; Fig. S13), and the size distribution of the selected proteinosomes was determined (Fig. S13). Data are presented as violin plots and histograms showing the distribution of intraproteinosomal ligand concentration and proteinosome size.

**Nonspecific ligand association**

To assess nonspecific association (Fig. S5), FITC-labelled proteinosomes lacking streptavidin were prepared following the proteinosome fabrication protocol. The proteinosomes were then buffer exchanged from water into localization buffer by two cycles of magnetic separation and resuspension. The proteinosomes were incubated overnight at 4 °C with Cy3-labelled, non-biotinylated ssDNA (**ssDNA$_{top_fluor}$**), CC protein or cysteine-engineered **CC$_{ditopic}$** (Fig. S1), or ssDNA–CC ligand (**ligand$_{fluor}$**). Following incubation, proteinosomes were washed thoroughly by magnetic separation to remove unbound components. The washed proteinosomes were imaged by fluorescence microscopy (Nikon, Eclipse Ti2-D-PD) to assess component association. FITC-labelled proteinosomes prepared with streptavidin and loaded with **ligand$_{fluor}$** using the standard DNA-mediated loading procedure served as the positive control. Data are presented as violin plots showing individual proteinosomes. Horizontal lines indicate the mean, and error bars denote s.d. Fold differences were calculated as ratios of mean fluorescence between the indicated conditions.

**Intraproteinosomal ligand retention in culture conditions**

FITC-labelled proteinosomes containing DNA-localized Cy3-labelled ssDNA–CC ligand (**ligand$_{fluor}$**) were transferred into serum-free DMEM (Gibco, 41966), serum-free 293 SFM II medium (SFM, Gibco, 10503283), or protection medium. Samples were imaged by fluorescence microscopy (Nikon, Eclipse Ti2-D-PD) before and after overnight incubation under standard culture conditions (Fig. S8). Cy3 fluorescence was normalized to the mean pre-incubation fluorescence for each condition. Data are presented as violin plots showing individual proteinosomes. Horizontal lines indicate the mean, and error bars denote s.d. (Fig. 2d). Fold reductions were calculated as the ratio of mean pre-incubation fluorescence to the mean post-incubation fluorescence.

**Localization-component stability in culture conditions**

To assess the retention of individual localization components (Fig. 2c; Fig. S9), FITC-labelled proteinosomes were prepared with either unlabelled streptavidin or Alexa Fluor 546-labelled streptavidin (Invitrogen, 10053112). For the labelled preparation, Alexa Fluor 546-labelled streptavidin was included during proteinosome assembly at a final concentration of 1 µM, with water used in place of the corresponding volume of unlabelled streptavidin. Proteinosomes containing unlabelled streptavidin were loaded separately with Cy3-labelled biotinylated ssDNA (**ssDNA$_{anchor_fluor}$**), a Cy3-labelled DNA duplex, or Cy3-labelled ssDNA–CC ligand (**ligand$_{fluor}$**). Loading was performed by immobilizing biotinylated ssDNA on streptavidin, followed, where applicable, by hybridization of the complementary strand. The DNA duplex was generated by immobilizing biotinylated ssDNA (**ssDNA$_{anchor}$**) and hybridizing complementary Cy3-labelled ssDNA (**ssDNA$_{top_fluor}$**). DNA-localized ligand was generated by hybridizing **ligand$_{fluor}$** to immobilized **ssDNA$_{anchor}$**. The samples were transferred into serum-free DMEM (Gibco, 41966), serum-free 293 SFM II medium (SFM, Gibco, 10503283), or protection medium. Samples were imaged by fluorescence microscopy (Nikon, Eclipse Ti2-D-PD) before and after overnight incubation under standard culture conditions (Fig. S9). Alexa Fluor 546 fluorescence was used to monitor streptavidin retention, while Cy3 fluorescence was used to monitor the retention of DNA-based components. Post-incubation fluorescence was normalized to the corresponding mean pre-incubation fluorescence. Data are presented as violin plots showing individual proteinosomes. Horizontal lines indicate the mean, and error bars denote s.d. Fold reductions were calculated as the ratio of mean pre-incubation fluorescence to the mean post-incubation fluorescence.

##### Fluorescence analysis of DNA-programmed ligand release during cell co-incubation

HEK293T cells (ATCC, CRL3216) were seeded at 8.0 × 10³ cells per well in a 96-well plate (Greiner) in standard culture medium and incubated overnight under standard culture conditions. The following day, FITC-labelled proteinosomes containing DNA-localized Cy3-labelled ssDNA–CC ligand (**ligand$_{fluor}$**) were transferred into stability medium. The cell culture medium was replaced with 100 µL of proteinosome suspension per well, after which either 1 µM non-complementary ssDNA (**ssDNA$_{scramble}$**) or 1 µM cognate ssDNA (**ssDNA$_{input}$**) was added to establish protection-medium or activation-medium conditions, respectively. Cells and proteinosomes were then co-incubated overnight under standard culture conditions (Fig. 3a-b; Fig. S11). Following co-incubation, intraproteinosomal Cy3 fluorescence was assessed by fluorescence microscopy (Nikon, Eclipse Ti2-D-PD) (Fig. S11). Cy3 fluorescence was normalized to the mean fluorescence intensity of the protection-medium condition. The fold reduction was calculated as the ratio of mean fluorescence in protection medium to the mean fluorescence in activation medium. Data are presented as lines showing the mean, with individual data points representing biological replicates (Fig. 3b). Statistical comparisons were performed using a two-sided Welch's t-test.

##### Orthogonal DNA-programmed localization and release of AND-gate ligands

Two proteinosome populations were prepared separately. Population **A** was FITC-labelled and contained Cy5-labelled **ligand$_{AND1}$**, whereas population **B** was DyLight405-labelled and contained Cy3-labelled **ligand$_{AND2}$**. Each ligand was localized using its corresponding biotinylated ssDNA anchor and complementary ssDNA–CC ligand strand (Table S3-S5). The two populations were mixed at a 1:1 ratio by proteinosome number and divided into four samples. The samples received either no input (00), 1 µM **ssDNA$_{inAND2}$** (01), 1 µM **ssDNA$_{inAND1}$** (10), or both inputs at 1 µM each (11). Samples were incubated overnight at room temperature and analysed by fluorescence microscopy (Nikon, Eclipse Ti2-D-PD) (Fig. S19). Cy5 fluorescence was used to assess **ligand$_{AND1}$** and Cy3 fluorescence to assess **ligand$_{AND2}$**. Fluorescence values were normalized to the 00 condition for the corresponding channel. Data are presented as violin plots showing individual proteinosomes. Horizontal lines indicate the mean, and error bars denote s.d. Fold reductions were calculated as the ratio of mean fluorescence in the 00 condition to the mean fluorescence each input condition.

##### Microfluidic device fabrication

A two-layer microfluidic device (Fig. S6) was used to facilitate the physical trapping and in situ imaging of ligand-loaded proteinosomes. The device consisted of a 1.5 × 2 mm or 1.5 × 1.5 mm localization chamber with polydimethylsiloxane (PDMS) pillars, a filtering chamber, inlet channels with pneumatically actuated Quake style push-up valves[73] and an outlet channel. Master moulds for the two layers were fabricated on separate silicon wafers using standard photolithography techniques.[73] The moulds for bottom and top layers were made by spincoating SU83050 to a height of 50 µm, and spincoating AZ 40xt to a height of 40 µm, respectively. After development, the AZ 40xt mould was reflowed, resulting in rounded channels with a height of ~60 µm at the centre. The microfluidic device was assembled from polydimethylsiloxane using standard multilayer soft lithography techniques[73] and plasma-bonded to 75 × 25 mm glass microscope slides. The microfluidic trapping device was mounted on the stage of a confocal laser scanning microscope (CLSM, Leica SP8) equipped with solid-state lasers at 488 nm (FITC) and 552 nm (Cy3), with a 10× objective (NA 0.40), and a hybrid detector in the photon counting mode. Control channels were filled with Milli-Q water and actuated using a pneumatic valve array (FESTO), which was in turn actuated using a programmable logic controller (PLC, WAGO Kontakttechnik). The PLC was connected to the Ethernet port of a PC and controlled using a custom MATLAB GUI. The pressure in the control channels was 2 bar. The pressure to the inlet channels was controlled using adjustable pressure regulators (FlowEZ, Fluigent).

##### Microfluidic strand-displacement release kinetics

Buffer solution was connected to the buffer inlet, and air bubbles were removed from the flow channels by pressurizing the buffer channel at 1 bar while closing the remaining inlet and outlet valves. The channels were subsequently flushed with buffer. Next ligand-containing proteinosomes were introduced into the trapping array through the proteinosome inlet at a pressure of 10 mbar. The inlet was then closed, and the trapped proteinosomes were gently washed with buffer for 5–10 min to equilibrate the system while minimizing displacement from the traps. The confocal microscope was focused on the trapping chamber, and time-lapse imaging was initiated (Fig. 1e-f; Fig. S7; Supplementary Movie 1). The experiment was performed at room temperature, and the initial steady-state fluorescence signals were recorded. Proteinosomes prepared using Dynabeads M-270 Amine were used for this experiment. During fabrication, the amine-functionalized M-270 beads accumulated centrally within the proteinosomes, resulting in a central non-fluorescent bead-rich region in the confocal fluorescence images (Fig. S7). The input ssDNA strand was

diluted and mixed to 100 nM in buffer solution and loaded to the DNA inlet port. The TMSD-reaction was started by flowing the input DNA into the trapping chamber at 10 mbar for 10 s. The inlet and outlet valves were then closed and remained closed throughout the experiment to ensure that TMSD-release occurred via diffusion.
Time-lapse fluorescence images were analysed using MATLAB. Individual proteinosomes were identified in the FITC channel, and their positions were assessed throughout the times series. Proteinosomes that moved substantially during the experiment were excluded to minimize errors arising from changes in ROI placement. For each remaining proteinosome, the corresponding ROI was transferred to the Cy3 channel. The central non-fluorescent bead-rich regions were excluded from fluorescence quantification, and Cy3 intensity was quantified from the surrounding regions at each time point (Fig. S7). Background fluorescence was subtracted from each measurement, and individual fluorescence trajectories were normalized to their respective baseline fluorescence before input addition (Fig. 1f).

##### Medium-dependent cellular metabolic activity

HEK293T cells (ATCC, CRL3216) were seeded at 8.0 × $10^3$ cells per well in a 96-well plate (Greiner) in standard culture medium and incubated overnight under standard culture conditions. The following day, the cell culture medium was replaced with 100 µL per well of either standard culture medium or protection medium. Cells were incubated overnight under standard culture conditions and cellular metabolic activity was assessed using the MTT assay (Fig. 2g). Metabolic activity in protection medium was normalized to the mean metabolic activity of cells incubated in standard culture medium. Data are presented as bar plots showing the mean, with individual data points representing biological replicates. Statistical comparisons were performed using a two-sided Welch's t-test.

##### Cellular metabolic activity of proteinosome-cell co-incubation

HEK293T cells (ATCC, CRL3216) were seeded at 8.0 × $10^3$ cells per well in a 96-well plate (Greiner) in standard culture medium and incubated overnight under standard culture conditions. The following day, FITC-labelled proteinosomes were transferred into stability medium. The cell culture medium was replaced with 100 µL per well of either stability medium alone or proteinosome suspension. Cells were incubated overnight under standard culture conditions. Following co-incubation, proteinosome abundance was visualized by fluorescence microscopy (Nikon, Eclipse Ti2-D-Pd) before assessment of cellular metabolic activity by MTT assay (Fig. S12; Fig. 3c). Metabolic activity in presence of proteinosomes was normalized to the mean metabolic activity of cells incubated in stability medium alone. Data are presented as bars showing the mean, with individual data points representing biological replicates. Statistical comparisons were performed using a two-sided Welch's t-test.

##### MTT metabolic activity assay

Cellular metabolic activity was measured using the MTT assay (3-(4,5-Dimethylthiazol-2-yl)-2,5-Diphenyltetrazolium Bromide, Thermo Scientific, M6494) (Fig. 2g; Fig. 3c). The cell culture medium was aspirated, and the cells were washed with PBS. Phenol-red-free culture medium (100 µL) and MTT reagent (10 µL; 5 mg $mL^{-1}$ in PBS) were added to each well, followed by incubation at 37 °C for 4 h. The medium was then removed, leaving 25 µL per well, and 50 µL DMSO was added to dissolve the formazan crystals. The contents of each well were thoroughly resuspended by pipetting, incubated at 37 °C for 10 min and mixed again. Absorbance was measured at 540 nm using a BioTek-Gen5 plate reader (BioTek) and corrected using a DMSO-only blank.

##### Mammalian transient transfection

For CC–GEMS receptor expression, HEK293T cells (ATCC, CRL3216) were seeded at 1.2 × $10^4$ cells per well in a 96-well plate (Greiner) in standard culture medium and incubated under standard culture conditions. The following day, cells were transfected for 5 h with 150 ng total plasmid DNA per well using Lipofectamine 2000 (0.25 µL per well; Life Technologies, 11668019) in 60 µL Opti-MEM I (Thermo Scientific, 31985062). The DNA mixture comprised 116.28 ng of the CC–GEMS receptor construct, 4.89 ng of the STAT3 expression vector pLS15 and 28.83 ng of either STAT3-inducible SEAP reporter plasmid pLS13 or the corresponding IL-10 reporter plasmid pLS13-IL-10. Following the 5 h transfection period, the transfection mixture was replaced with 100 µL of the medium specified for each experiment. Cells were then incubated overnight under standard culture conditions before analysis of SEAP activity (Fig. 2f; Fig. 3e; Fig. S18; Fig. 4c) or IL-10 secretion (Fig. S20).

##### Ligand-mediated CC–GEMS activation

HEK293T receiver cells were transiently transfected to express the cognate AA CC–GEMS receptor and the SEAP reporter. Following transfection, the transfection medium was removed and replaced with 100 µL per well of either standard culture medium or protection medium. Each medium was tested either alone or supplemented with 120 nM $\mathbf{CC_{ditopic}}$ (Fig. S1) or DNA-conjugated ligand ($\mathbf{ligand_{ssDNA–CC}}$). All conditions were incubated overnight under standard culture conditions. Following incubation, culture supernatants were collected for SEAP analysis, and cells were harvested for viable-cell counting (Fig. 2e-f).
Cells were detached by trypsinization, collected and resuspended in 200 µL of standard culture medium. Total and nonviable cell counts were determined using a NucleoCounter NC-100 (ChemoMetec, 900-0004), according to the manufacturer's A100/B reagent protocol. For total-cell counting, 50 µL of cell suspension was mixed with 50 µL of Reagent A100 lysis buffer (ChemoMetec, 910-0003) and 50 µL of Reagent B stabilizing buffer (ChemoMetec, 910-0002). The resulting mixture was loaded into a NucleoCassette (ChemoMetec, 941-0002) and analysed. Nonviable cells were counted separately using an untreated aliquot of the cell suspension. Total and nonviable cell numbers were calculated from the measured cell concentrations,

accounting for sample dilution and the total suspension volume. Viable-cell numbers were calculated by subtracting the nonviable cell number from the total cell number. SEAP activity was normalized to the corresponding viable-cell number and expressed relative to the response elicited by $\mathbf{CC}_{\mathrm{ditopic}}$ in standard culture medium. Data are presented as bar plots showing the mean, with individual data points representing biological replicates (Fig. 2f). Statistical comparisons were performed using one-way analysis of variance (ANOVA) across all experimental conditions, followed by Tukey's multiple-comparison test. To assess ligand integrity during CC–GEMS activation, supernatants from three independent biological replicates incubated overnight with 120 nM $\mathbf{ligand}_{\mathrm{ssDNA-CC}}$ in protection medium were analysed by anti-SUMO Western blotting (Fig. S10).

**Proteinosome counting and dose preparation for cell co-incubation experiments**

The target proteinosome dose for cell co-incubation experiments was estimated from the proteinosome size distribution and intraproteinosomal ligand-loading measurements shown in Fig. S13. Assuming complete release of the loaded ligand, approximately $4.69 \times 10^5$ proteinosomes per 100 µL corresponded to a nominal ligand dose equivalent of 120 nM. This represents a theoretical complete-release equivalent rather than a measured extracellular ligand concentration. Because ligand loading was not independently calibrated for each ligand preparation, these dose estimates should be regarded as approximate rather than as independently verified equivalent ligand amounts across preparations.
To determine the fabrication batch size required to achieve the target particle number after processing, batches prepared using different fabrication volumes were evaluated by high-content screening fluorescence microscopy (Nikon, Eclipse Ti2-D-PD). Samples were diluted, and 1 µL droplets of each diluted suspension were placed on a microscope slide and allowed to settle for 1–2 min. The entire droplet was imaged, and the proteinosomes were counted. Counts were converted to the corresponding numbers in the original suspensions using the dilution factor. The fabrication batch size yielding a proteinosome number closest to approximately $5.5 \times 10^5$ proteinosomes per 100 µL was selected to account for anticipated losses during water-to-buffer exchange, biotinylated DNA immobilization, washing, ligand loading, subsequent washing and buffer-to-medium exchange.
Following completion of these processing steps, proteinosome numbers were determined again using the same microscopy-based counting procedure to identify the fabrication batch size yielding a final proteinosome number closest to the target of approximately $4.69 \times 10^5$ proteinosomes per 100 µL. A fabrication batch volume in the range of 40–50 µL most closely met both the pre-processing and final proteinosome-number targets. This range was therefore selected as the standard for batch preparation. For experiments involving multiple wells or conditions, the batch volume was increased proportionally to the number of 100 µL aliquots required.

**DNA-programmable proteinosome–cell communication for SEAP activation**

FITC-labelled proteinosomes containing DNA-localized Cy3-labelled ssDNA–CC ligand ($\mathbf{ligand}_{\mathrm{activation}}$) were transferred into stability medium. HEK293T receiver cells were transiently transfected to express either the cognate AA or non-cognate BB CC–GEMS receptor together with the SEAP reporter. Following transfection, the transfection medium was removed and replaced with 100 µL of the experimental condition per well. Cognate receiver cells were incubated in either protection medium alone, protection medium supplemented with 120 nM free $\mathbf{ligand}_{\mathrm{activation}}$ or a suspension of ligand-containing proteinosomes in stability medium. For proteinosome-containing conditions, 100 µL of suspension containing approximately $4.69 \times 10^5$ proteinosomes was added per well. Subsequently, either 1 µM non-complementary ssDNA ($\mathbf{ssDNA}_{\mathrm{scramble}}$) or 1 µM cognate ssDNA ($\mathbf{ssDNA}_{\mathrm{input}}$) was added to the proteinosome suspension to establish protection-medium or activation-medium conditions, respectively. Non-cognate receiver cells were co-incubated with proteinosomes under the same protection and activation conditions as the cognate receiver cells. All conditions were incubated overnight under standard culture conditions. Following incubation, culture supernatants were collected for SEAP analysis (Fig. 1a; Fig. 3d-e). SEAP activity was normalized to the mean response elicited by 120 nM free $\mathbf{ligand}_{\mathrm{activation}}$. Data are presented as bar plots showing the mean, with individual data points representing biological replicates (Fig. 3e). Statistical comparisons were performed using one-way analysis of variance (ANOVA) across all experimental conditions, followed by Tukey's multiple-comparison test. To assess the presence and integrity of ligand-derived material in the culture supernatants following proteinosome–cell co-incubation, supernatants from conditions involving cognate receiver-cell were analysed by anti-SUMO Western blotting (Fig. 3f).

**CC–GEMS activation by complementary AND-gate ligand components**

HEK293T receiver cells were transiently transfected to express the cognate AA CC–GEMS receptor and the SEAP reporter. Following transfection, the transfection medium was removed and replaced with 100 µL per well of protection medium containing 2 µM non-complementary ssDNA ($\mathbf{ssDNA}_{\mathrm{scramble}}$). Four ligand-addition configurations were tested: no ligand components (00), 120 nM $\mathbf{ligand}_{\mathrm{AND2}}$ alone (01), 120 nM $\mathbf{ligand}_{\mathrm{AND1}}$ alone (10), or 120 nM of each ligand component (11) (Fig. S18). All conditions were incubated overnight under standard culture conditions. Following incubation, culture supernatants were collected for SEAP analysis. SEAP activity was normalized to the mean response of the dual-input condition (11). Data are presented as bar plots showing the mean, with individual data points representing biological replicates. Statistical comparisons were performed using one-way analysis of variance (ANOVA) across all experimental conditions, followed by Tukey's multiple-comparison test.

**Distributed DNA-programmable proteinosome–cell communication**

Two proteinosome populations were prepared separately. Population **A** was FITC-labelled and contained DNA-localized Cy5-labelled ssDNA–CC ligand ($\mathbf{ligand}_{\mathrm{AND1}}$), whereas population **B** was DyLight405-labelled and contained DNA-localized Cy3-labelled ssDNA–CC ligand ($\mathbf{ligand}_{\mathrm{AND2}}$). Each ligand was localized within its corresponding proteinosome population using the appropriate biotinylated DNA anchor and complementary ssDNA–CC ligand strand. The two populations were mixed at a 1:1

ratio by proteinosome number and transferred into stability medium. HEK293T receiver cells were transiently transfected to express the cognate AA CC–GEMS receptor and the SEAP reporter. Following transfection, the transfection medium was removed and replaced with 100 µL per well of either protection medium supplemented with 120 nM each of free **ligand**$_{\mathbf{AND1}}$ and **ligand**$_{\mathbf{AND2}}$, or a suspension of ligand-containing proteinosome populations **A** and **B** in stability medium. Protection medium consisted of stability medium supplemented with 2 µM non-complementary ssDNA (**ssDNA**$_{\mathbf{scramble}}$). For proteinosome-containing conditions, 100 µL of suspension containing approximately 4.69 × $10^5$ proteinosomes from each population was added per well, corresponding to approximately 9.38 × $10^5$ proteinosomes in total. After addition of the proteinosome suspension, DNA was added to establish one of four input configurations while maintaining a total DNA concentration of 2 µM: no activating input (00), 1 µM **ssDNA**$_{\mathbf{inAND2}}$ alone (01), 1 µM **ssDNA**$_{\mathbf{inAND1}}$ alone (10), or 1 µM of each input strand (11) (Fig. 4). The 00-condition contained 2 µM **ssDNA**$_{\mathbf{scramble}}$; in the single-input conditions, the relevant input strand was combined with **ssDNA**$_{\mathbf{scramble}}$ to maintain a total DNA concentration of 2 µM. All conditions were incubated overnight under standard culture conditions. Following incubation, culture supernatants were collected for SEAP analysis. SEAP activity was normalized to the mean response elicited by 120 nM free ligand-pair (120 nM each of **ligand**$_{\mathbf{AND1}}$ and **ligand**$_{\mathbf{AND2}}$). Data are presented as bar plots showing the mean, with individual data points representing biological replicates (Fig. 4c). Statistical comparisons were performed using one-way analysis of variance (ANOVA) across all experimental conditions, followed by Tukey's multiple-comparison test.

**SEAP reporter assay**

Culture supernatants were collected and clarified by centrifugation at 300 × *g* for 5 min to remove cell debris (Fig. 2f). Supernatants collected after proteinosome–cell co-incubation were clarified by centrifugation at 3,000 × *g* for 5 min to remove residual proteinosomes (Fig. 3e; Fig. 4c). Samples were analysed immediately or stored at −80 °C immediately after collection. Samples were not heat-inactivated upon collection, as heating was considered potentially incompatible with the ssDNA–CC conjugate and could compromise subsequent Western Blot analysis, where applicable. Instead, heat inactivation was performed during the SEAP assay, according to the manufacturer's protocol, as described below. SEAP activity in culture supernatants was quantified colorimetrically using the secreted alkaline phosphatase (SEAP) assay, according to the manufacturer's instructions. The assay measures the increase in absorbance resulting from hydrolysis of *p*-nitrophenyl phosphate (pNPP) to *p*-nitrophenol (pNP). Sample dilution buffer (1×) was prepared by diluting 100 µL of 10× Sample Dilution buffer (Novus Biologicals, NBP2-25285, IMK-515-6) in 900 µL of Milli-Q water. Samples were diluted 1:2 by mixing 6 µL of culture supernatant with 6 µL of 1× sample dilution buffer. For the assay, 10 µL of diluted sample was combined with 10 µL of MilliQ-water in a clear 96-well plate (Greiner). The plate was sealed and incubated at 65 ºC for 30 min for heat inactivation. The sealed plate was gently tapped to collect the liquid at the bottom of the wells. The pNPP substrate solution (1 mg $mL^{-1}$) was prepared by diluting 1 mL of 5× diethanolamine substrate buffer (Thermo Scientific, 34064) in 4 mL of Milli-Q water, followed by dissolution of a 5 mg pNPP tablet (Thermo Scientific, 34047). After heat inactivation, 100 µL of substrate solution was added to each well, yielding a final pNPP concentration of 0.83 mg $mL^{-1}$. Absorbance was measured at 405 nm every 30 s for 60 min at 25 °C using a BioTek-Gen5 plate reader (BioTek). SEAP activity was determined from a calibration curve generated using known concentrations of pNP (Fig. S21). Absorbance measurements were converted to pNP amounts and SEAP activity was calculated from the slope of the time trace, expressed in units per litre (U $L^{-1}$). One unit was defined as the amount of enzyme that converts 1 µmol of pNPP in 5 µL of culture supernatant per minute at 25 °C. Relative SEAP activity was calculated by normalizing the resulting reaction rates to the designated reference condition.

**DNA-programmable proteinosome–cell communication for IL-10 secretion**

FITC-labelled proteinosomes containing DNA-localized Cy3-labelled ssDNA–CC ligand (**ligand**$_{\mathbf{activation}}$) were transferred into stability medium. HEK293T receiver cells were transiently transfected to express the cognate AA CC–GEMS receptor together with the IL-10 reporter. Following transfection, the transfection medium was removed and replaced with 100 µL per well of either protection medium alone, protection medium supplemented with 120 nM free **ligand**$_{\mathbf{activation}}$, or a suspension of the ligand-containing proteinosomes in stability medium. For proteinosome-containing conditions, 100 µL of suspension containing approximately 4.69 × $10^5$ proteinosomes was added per well. Subsequently, either 1 µM non-complementary ssDNA (**ssDNA**$_{\mathbf{scramble}}$) or 1 µM cognate ssDNA (**ssDNA**$_{\mathbf{input}}$) was added to the proteinosome suspension to establish protection-medium or activation-medium conditions, respectively. All conditions were incubated overnight under standard culture conditions. Following incubation, culture supernatants were collected for IL-10 analysis using ELISA (Fig. S20). IL-10 secretion was normalized to the mean response elicited by 120 nM free **ligand**$_{\mathbf{activation}}$. Data are presented as bar plots showing the mean, with individual data points representing biological replicates. Statistical comparisons were performed using one-way analysis of variance (ANOVA) across all experimental conditions, followed by Tukey's multiple-comparison test.

**IL-10 quantification by ELISA**

Human IL-10 in culture supernatants was quantified using the Human IL-10 ELISA MAX Deluxe Set (BioLegend, 430604), according to the manufacturer's instructions (Fig. S20). The kit supplied the capture antibody (200×), detection antibody (200×), recombinant human IL-10 standard, coating buffer (5×), assay diluent (5×), avidin–HRP (1000×), and TMB substrate (a 1:1 mixture of solutions A and B). The 1× coating buffer was prepared by diluting 2.4 mL of 5× coating buffer in 9.6 mL deionized water. The 1× assay diluent was prepared by mixing 12 mL of 5× assay diluent with 48 mL PBS (pH 7.4). Wash buffer consisted of PBS containing 0.05% Tween 20 (Sigma), and 1 M $H_2SO_4$ was used as the stop solution. A Nunc MaxiSorp 96-well ELISA plate (Thermo Scientific, 442404) was coated with 100 µL per well of capture antibody diluted 1:200 in 1× coating buffer and incubated overnight at 4 °C. The plate was washed four times and blocked with 200 µL per well of 1× assay diluent for 1 h at room temperature with shaking. After four washes, culture supernatants diluted 100-fold were added and incubated for 2 h at room temperature with shaking. The plate was washed four times and incubated with 100 µL per well of detection antibody diluted

1:200 in 1x assay diluent for 1 h at room temperature with shaking. Following four washes, 100 µL per well of avidin–HRP diluted 1:1,000 in 1x assay diluent was added and incubated for 30 min at room temperature with shaking. The plate was then washed five times, leaving wash buffer in each well for 30–60 s per wash. TMB substrate (100 µL per well) was added, and the plate was incubated in the dark for 30 min. The reaction was stopped by adding 100 µL of stop solution per well. Absorbance was measured at 450 nm using a BioTek-Gen5 plate reader (BioTek), with reference correction at 570 nm. Sample IL-10 concentrations were determined from a calibration curve generated using known concentrations of recombinant human IL-10 (Fig. S20). The 100-fold-diluted samples fell within the usable quantification range of the standard curve, and the dilution factor was applied to calculate IL-10 concentrations in the original culture supernatants (ng/mL). Measurements were normalized to the specified reference condition.

**Statistical analysis**

The numbers of biological replicates and proteinosomes analysed are reported separately in the relevant experimental sections. Cell-based data are presented as mean values with individual biological replicates shown as dots. For proteinosome-level fluorescence measurements, individual proteinosomes were treated as measurements within the sampled population rather than as independent biological replicates. Data are presented as histograms or violin plots, with the mean ± s.d. indicated. Two-sided Welch's t-tests were used for comparisons between two groups where indicated. Comparisons involving more than two groups were analysed by one-way ANOVA across all experimental conditions, followed by Tukey's multiple-comparison test where indicated. Measurements were normalized to the specified reference condition. Where applicable, fold reductions were calculated as the ratio of mean fluorescence in the reference condition to mean fluorescence in the comparison condition. Statistical analyses and graph generation were performed using GraphPad Prism. Statistical significance was defined as follows: ns, $P > 0.05$; *, $P \leq 0.05$; **, $P \leq 0.01$; ***, $P \leq 0.001$; ****, $P \leq 0.0001$.

#### Supplementary Movies

***Supplementary Movie 1.*** Microfluidic ligand release from proteinosomes. Time-lapse confocal microscopy recording with the green and yellow channels showing the fluorescence of FITC-labelled BSA–$NH_2$/PNIPAm proteinosomes and Cy3-labelled localized fluorescent ligands (**ligand**$_{\mathbf{fluor}}$), respectively. This video corresponds to the reaction shown in Fig. 1f in the main text and Supplementary Fig. S7. At t = 8 min, input strand (**ssDNA**$_{\mathbf{input}}$) is flowed into the microfluidic trapping chamber and displaces the ssDNA–CC ligand (**ligand**$_{\mathbf{fluor}}$) from the DNA anchor strand (**ssDNA**$_{\mathbf{anchor}}$) within the proteinosomes, resulting in release of **ligand**$_{\mathbf{fluor}}$ from the proteinosome lumen and thereby a decrease in Cy3 fluorescence.

#### Supplementary Figures

**a.** **$CC_{ditopic}$ coding sequence**

ATG GGC AGC AGC CAT CAT CAT CAT CAT CAC AGC TGC GGC ACT AGT AGC GGC GGT GGT AGC GAT AGC GAA

M G S S H H H H H H S C G T S S G G G S D S E

GTC AAT CAA GAA GCG AAA CCG GAA GTC GAA CCG GAA GTC AAA CCG GAA ACG CAC ATC AAC CTG AAA GTT

V N Q E A K P E V E P E V K P E T H I N L K V

AGT GAT GGC AGC TCT GAA ATT TTC TTT AAA ATT AAG AAA ACC ACG CCG CTG CGT CGC CTG ATG GAA GCG

S D G S S E I F F K I K K T T P L R R L M E A

TTT GCC AAA CGC CAG GGC AAA GAA ATG GAT AGC CTG CGC TTC CTG TAT GAC GGT ATT CGT ATC CAG GCG

F A K R Q G K E M D S L R F L Y D G I R I Q A

GAT CAA GCC CCG GAA GAT CTG GAC ATG GAA GAT AAC GAC ATT ATC GAA GCG CAT CGC GAA CAG ATT GGC

D Q A P E D L D M E D N D I I E A H R E Q I G

GGC CAT ATG AGC GGT CCT GGA TCC TCT CCG GAA GAC AAA ATT GCG CAG TTG AAA CAA AAA ATT CAG GCG

G H M S G P G S S P E D K I A Q L K Q K I Q A

CTC AAG CAA GAA AAT CAA CAG TTG GAG GAG GAG AAT GCA GCC CTG GAG TAT GGT GGA AGT CGA ATT CGG

L K Q E N Q Q L E E E N A A L E Y G G S R I R

TGG GAA TTT CAT CAC GGG GGT TCG GGT GGT TCA GGC GGC TCA GGA GGC TCC GGG GGT TCC GGA GGG

W E F H H G G S G G S G G S G G S G G S G G

AGC GGT GCT GAA GCC GCA GCC AAG GAA GCA GCA GCT AAA GAG GCC GCT GCG AAG GAA GCT GCC GCA

S G A E A A A K E A A A K E A A A K E A A A

AAG GAG GCG GCG GCG AAA GAG GCG GCA GCA AAA GCC GGA TCT GGT GGC AGT GGT GGC TCC GGC GGG

K E A A A K E A A A K A G S G G S G G S G G

TCA GGT GGC AGC GGG GGA TCA GGA GGT ACC AGC GGT CCT GGA TCC TCT CCG GAA GAC AAA ATT GCG

S G G S G G S G G T S G P G S S P E D K I A

CAG TTG AAA CAA AAA ATT CAG GCG CTC AAG CAA GAA AAT CAA CAG TTG GAG GAG GAG AAT GCA GCC CTG

Q L K Q K I Q A L K Q E N Q Q L E E E N A A L

GAG TAT GGT GGA AGT AAG CTT TGG AGC CAC CCG CAG TTC GAA AAA TAA

E Y G G S K L W S H P Q F E K *

His-tag | Cysteine modification | SUMO-tag | A' CC | AEK linker | Strep-tag II

**b.** **N-terminal sequence highlighting the engineered cysteine**

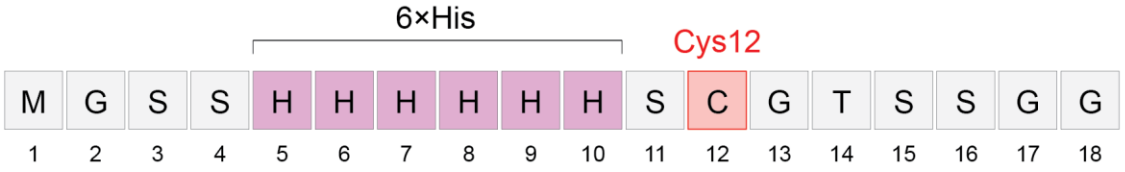


*SUMO domain begins immediately after this N-terminal region.*

***Supplementary Figure S1.*** Design of the cysteine-engineered **$CC_{ditopic}$** construct for cysteine-directed ssDNA conjugation. (a) Map of the **$CC_{ditopic}$** coding sequence showing the N-terminal 6×His tag, SUMO domain, coiled-coil domains, AEK linker and C-terminal Strep-tag II. An engineered cysteine was introduced within the N-terminal region to provide a thiol handle for subsequent ssDNA conjugation while positioning the conjugation site away from the receptor-binding coiled-coil domains. (b) N-terminal amino-acid sequence surrounding the engineered cysteine. Cys12 is positioned in the N-terminal region between the 6×His tag and the SUMO domain and provides an engineered thiol handle for subsequent thiol–maleimide coupling to ssDNA, enabling construction of the bifunctional ssDNA–CC signalling ligand.

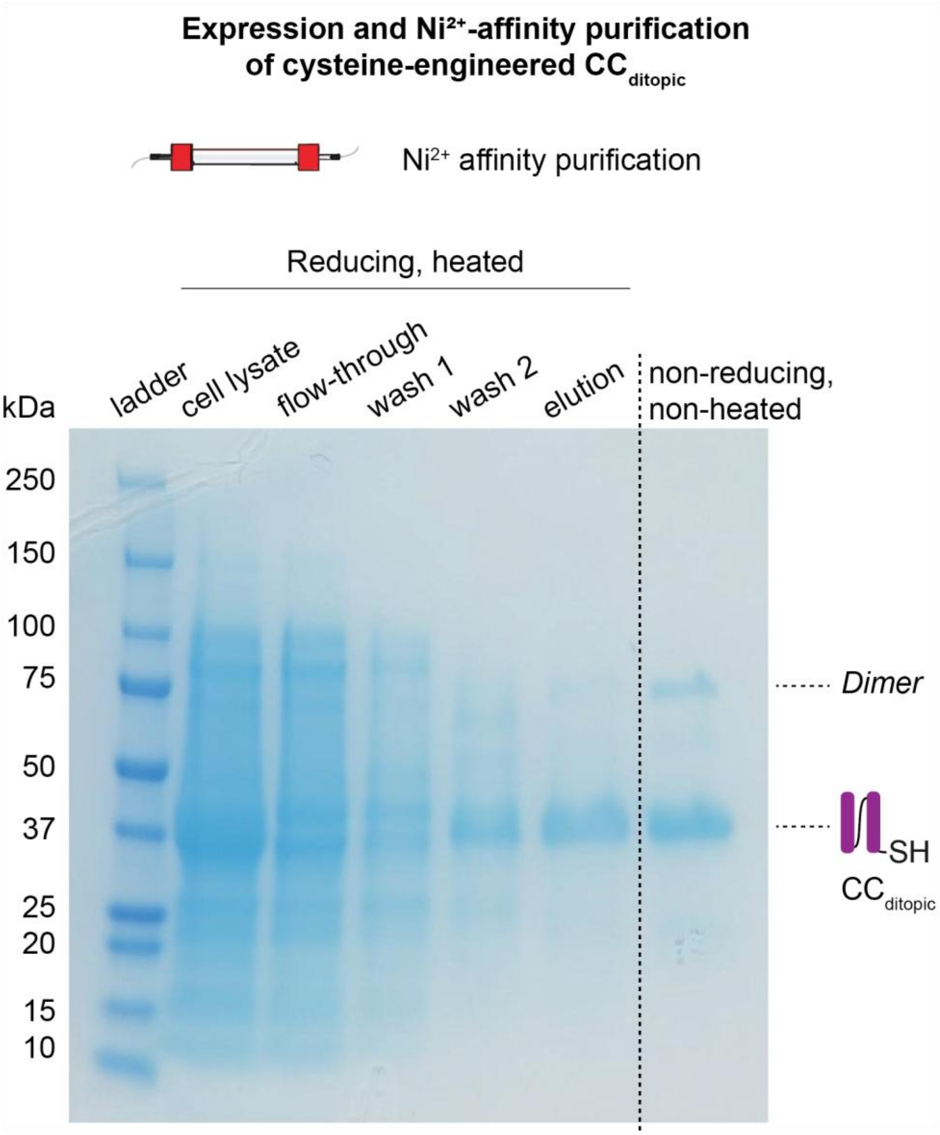


***Supplementary Figure S2.*** Expression and purification of cysteine-engineered **$CC_{ditopic}$**. SDS–PAGE analysis of **$CC_{ditopic}$** expression and $Ni^{2+}$-affinity purification (Methods). The cell lysate, flow-through, wash 1 and 2, and elution fractions were analysed under reducing, heated SDS–PAGE conditions, with **$CC_{ditopic}$** migrating at an apparent molecular mass of approximately 37 kDa. Under non-reducing, non-heated conditions, an additional species at approximately twice the monomeric molecular mass was observed, consistent with a putative disulfide-linked **$CC_{ditopic}$** dimer. Because Cys12 is the only cysteine residue in the construct, formation of this species supports accessibility of the engineered cysteine for thiol chemistry, consistent with its subsequent use for thiol–maleimide conjugation to generate the ssDNA–CC signalling ligand (Fig. 1b; Methods). The remaining purified protein was maintained under reducing conditions to preserve the engineered cysteine in its reactive thiol state. The corresponding uncropped image is provided in Supplementary Fig. S22.

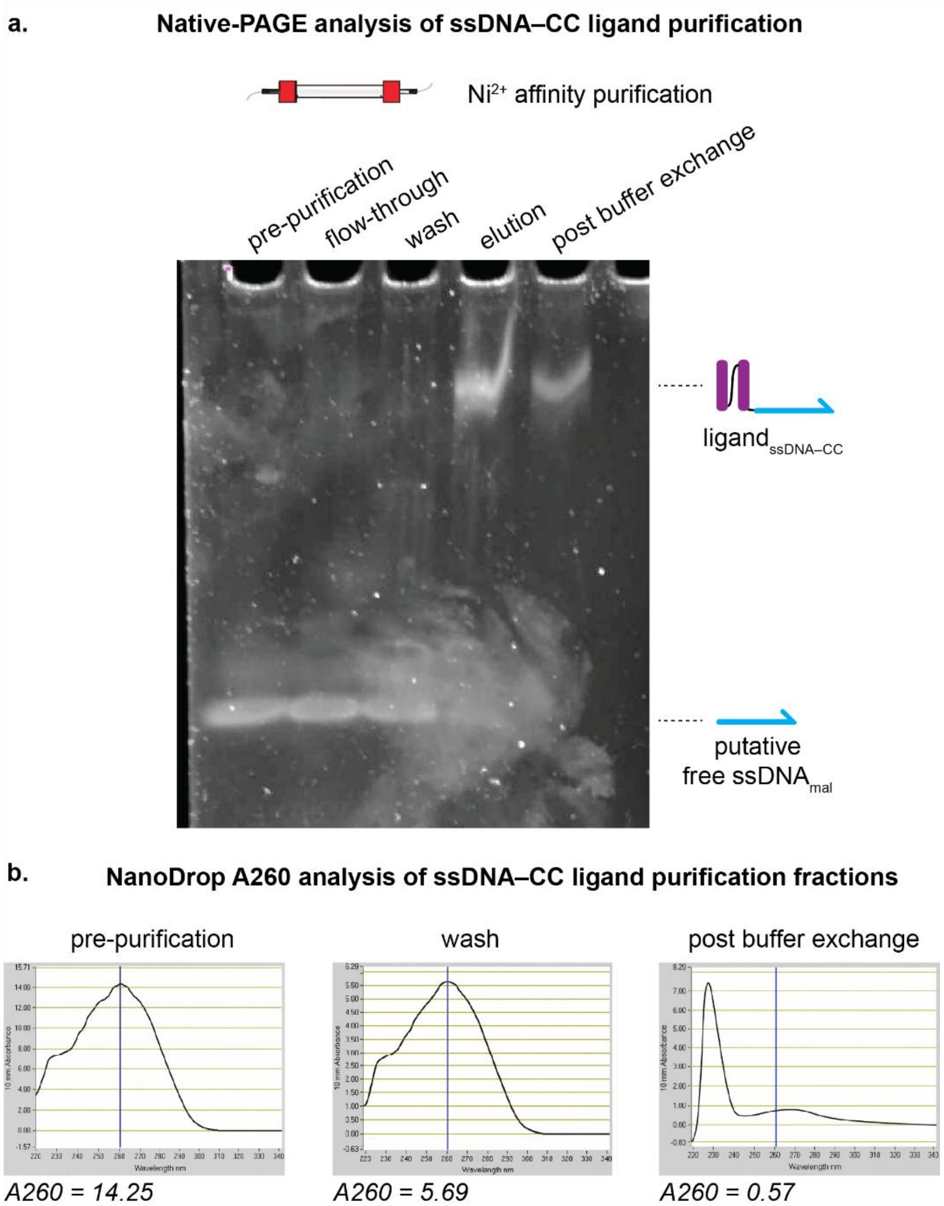


***Supplementary Figure S3.*** Purification of the ssDNA–CC signalling ligand. (a) Native-PAGE analysis of ssDNA–CC ligand purification fractions visualized by SYBR Gold staining (Methods). A slowly migrating nucleic-acid-containing species corresponding to the ssDNA–CC ligand is enriched in the elution and post buffer exchange fractions, whereas a rapidly migrating nucleic-acid species, putatively corresponding to unbound ssDNA, is progressively depleted during purification. (b) NanoDrop A260 spectra of the pre-purification, wash and post buffer exchange fractions. The measured A260 decreased from 14.25 in the pre-purification sample to 5.69 in the wash fraction and 0.57 in the post buffer exchange sample, corresponding to a 96% reduction in total nucleic-acid absorbance between the pre-purification and post buffer exchange samples. Together, the native-PAGE and absorbance measurements are consistent with enrichment of the ssDNA–CC conjugate and substantial depletion of excess nucleic acid during purification. Removal of excess ssDNA is important for the subsequent localization experiments in Fig. 1c,d, where free ssDNA could otherwise compete with programmed anchor-ligand hybridization. The corresponding uncropped image is provided in Supplementary Fig. S23.

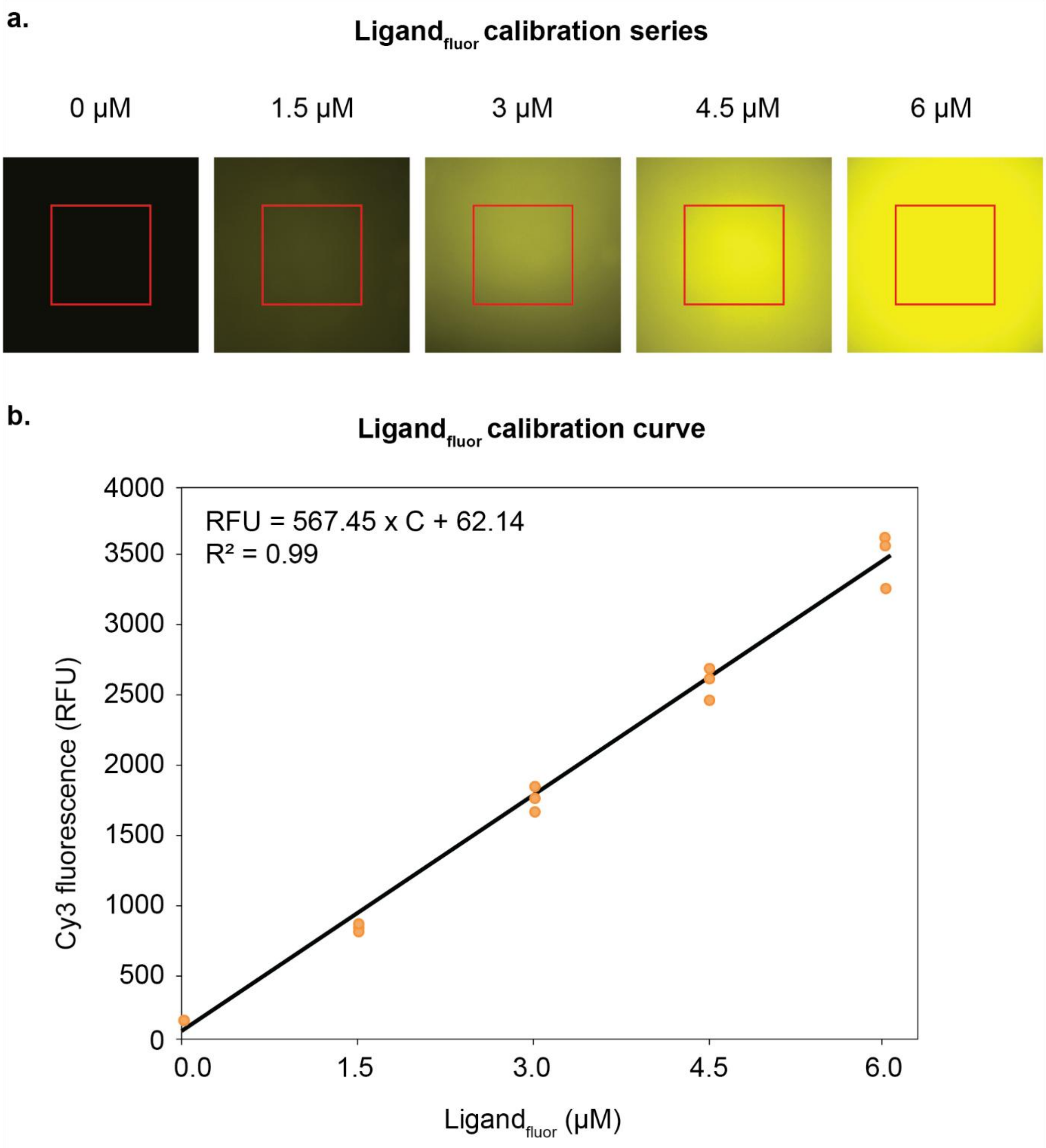


***Supplementary Figure S4.*** Cy3 fluorescence calibration for quantitative determination of intraproteinosomal ligand loading. (a) Representative Cy3 fluorescence images of **ligand$_{fluor}$** calibration standards containing 0, 1.5, 3, 4.5 and 6 µM **ligand$_{fluor}$**. Red squares indicate the fixed 800 × 800-pixel regions of interest (ROIs) used to measure mean Cy3 fluorescence intensity. (b) Calibration curve relating Cy3 fluorescence intensity to **ligand$_{fluor}$** concentration (Methods). Points represent individual replicate measurements ($n$ = 3 per concentration). Linear regression across the five concentrations gave RFU = 567.45 × C + 62.14 ($R^2$ = 0.99), where C is the **ligand$_{fluor}$** concentration in µM. The calibration was used to convert intraproteinosomal Cy3 fluorescence into the intraproteinosomal ligand concentrations reported in Fig. 1d.

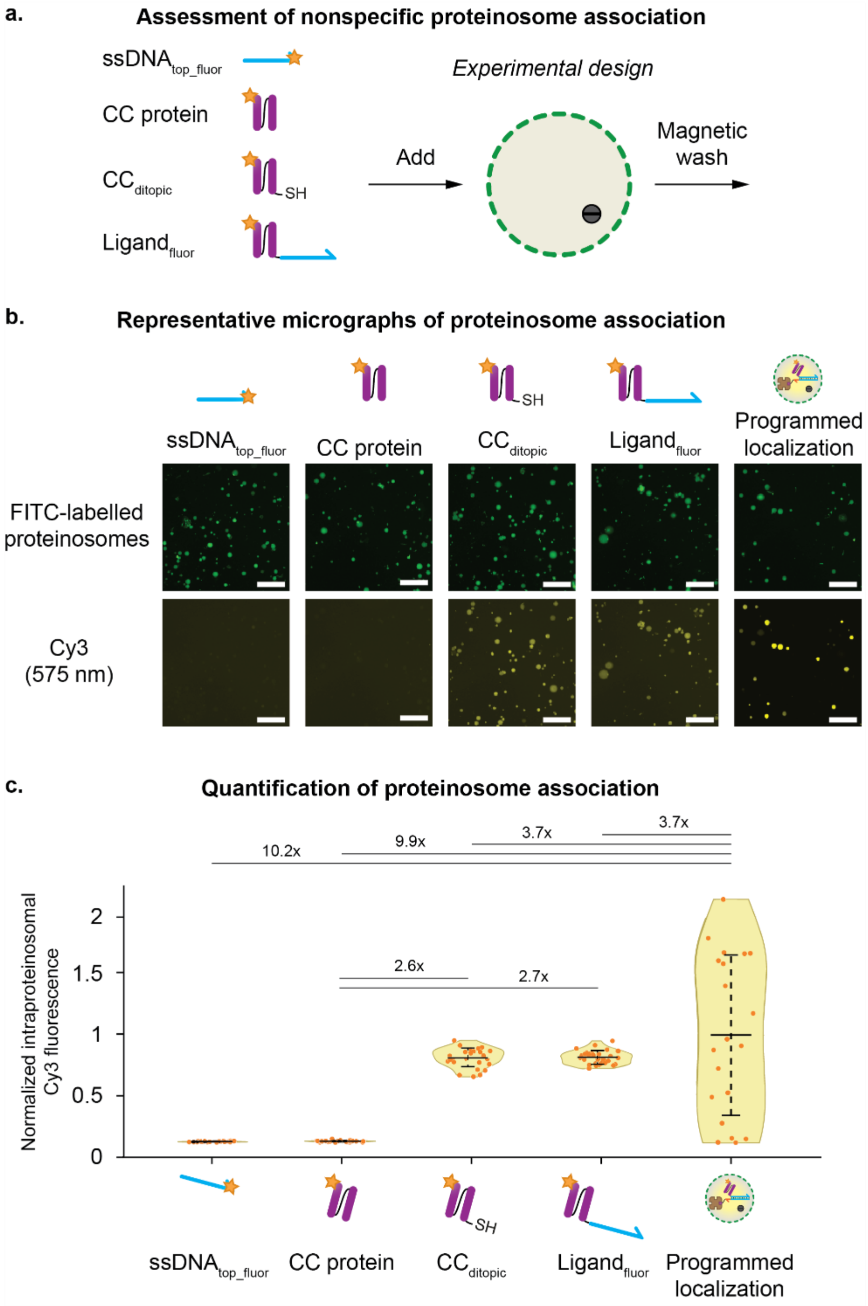


***Supplementary Figure S5.*** Nonspecific association of signalling components with proteinosomes in the absence of the programmed DNA-localization mechanism. (a) Experimental design for assessing proteinosome association in the absence of the cognate streptavidin–biotin–**ssDNA$_{anchor}$** localization system. FITC-labelled proteinosomes containing magnetic microparticles were incubated separately with Cy3-labelled ssDNA, CC protein, cysteine-engineered **CC$_{ditopic}$** or **ligand$_{fluor}$**, followed by magnetic washing (Methods). Proteinosomes containing **ligand$_{fluor}$** localized through the complete programmed DNA-localization mechanism were included as a positive control. (b) Representative fluorescence micrographs. Only low intraproteinosomal Cy3 fluorescence was observed for ssDNA and CC protein, whereas **CC$_{ditopic}$** and **ligand$_{fluor}$** showed greater association. The programmed-localization condition produced the strongest and most spatially confined intraproteinosomal Cy3 signal. A single independent experiment comprising all five association conditions is shown. Representative images in (b) were acquired using adjusted Cy3 settings to facilitate visualization of weak association signals and are therefore not intended for direct quantitative comparison between conditions. Scale bar, 250 µm. (c) Quantification of intraproteinosomal Cy3 fluorescence from images acquired using identical Cy3 intensity settings across all five conditions. Fluorescence values were normalized to the mean of the programmed-localization condition. Relative to CC protein, **CC$_{ditopic}$** and **ligand$_{fluor}$** showed 2.6-fold and 2.7-fold higher fluorescence, respectively. The programmed-localization condition showed substantially higher fluorescence than all conditions lacking the programmed localization mechanism. The increased association of **CC$_{ditopic}$** relative to CC protein is consistent with a contribution from the engineered cysteine, potentially involving thiol–disulfide exchange with BSA Cys34.[56–58] Residual association of **ligand$_{fluor}$** suggests that additional nonspecific interactions may also contribute. Individual proteinosome counts were 30 for ssDNA, 30 for CC protein, 30 for **CC$_{ditopic}$**, 30 for **ligand$_{fluor}$** and 20 for programmed localization. Up to 30 individually resolvable proteinosomes were analysed per condition. Data are presented as violin plots showing individual proteinosomes. Horizontal lines indicate the mean, and error bars denote s.d. Horizontal brackets denote the indicated fold differences, which were calculated as ratios of mean fluorescence between the indicated conditions.

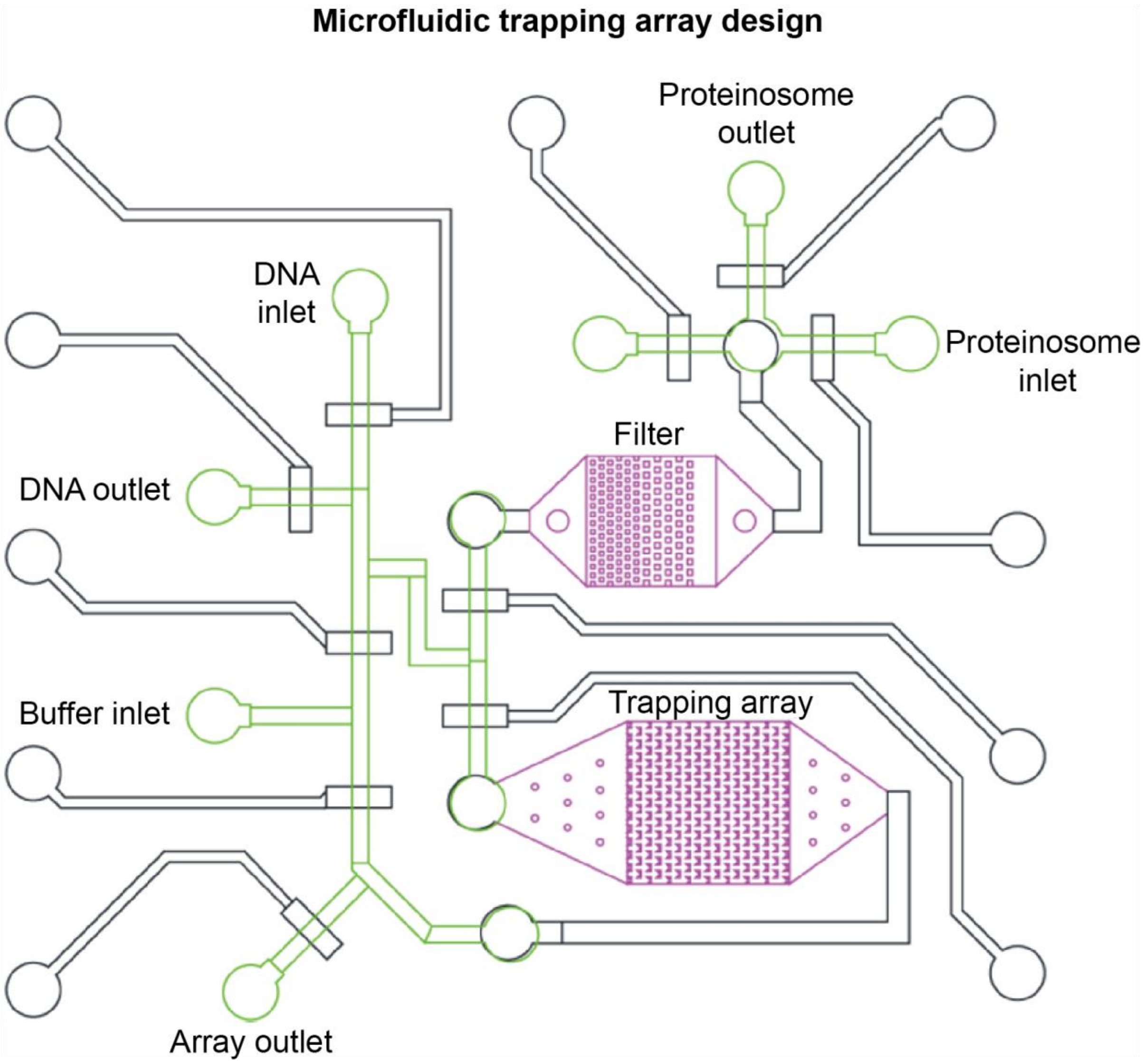


***Supplementary Figure S6.*** Two-layer microfluidic trapping array for controlled single-proteinosome imaging and reagent exchange. CAD schematic of the microfluidic device, adapted from previously developed designs.[10,12,13,54] The purple structures identify the filter and trapping array; black channels represent the bottom layer bonded to the glass slide, whereas green channels represent the rounded top-layer channels that can be actuated by pneumatic valves. At the layer-crossing regions, flow is transferred between the top and bottom layers because the rectangular bottom-layer geometry is required for the trapping array, while the rounded top-layer channels are compatible with pneumatic closure. During loading, proteinosomes enter through the proteinosome inlet and are directed through the filter towards the trapping array, where individual proteinosomes can be retained in defined trapping positions. After trapping, the surrounding solution can be exchanged by routing buffer or DNA-containing solution through the corresponding inlet and outlet while the proteinosomes remain spatially confined. This combination of physical trapping and controlled solution exchange provides the experimental basis for real-time single-proteinosome measurements.

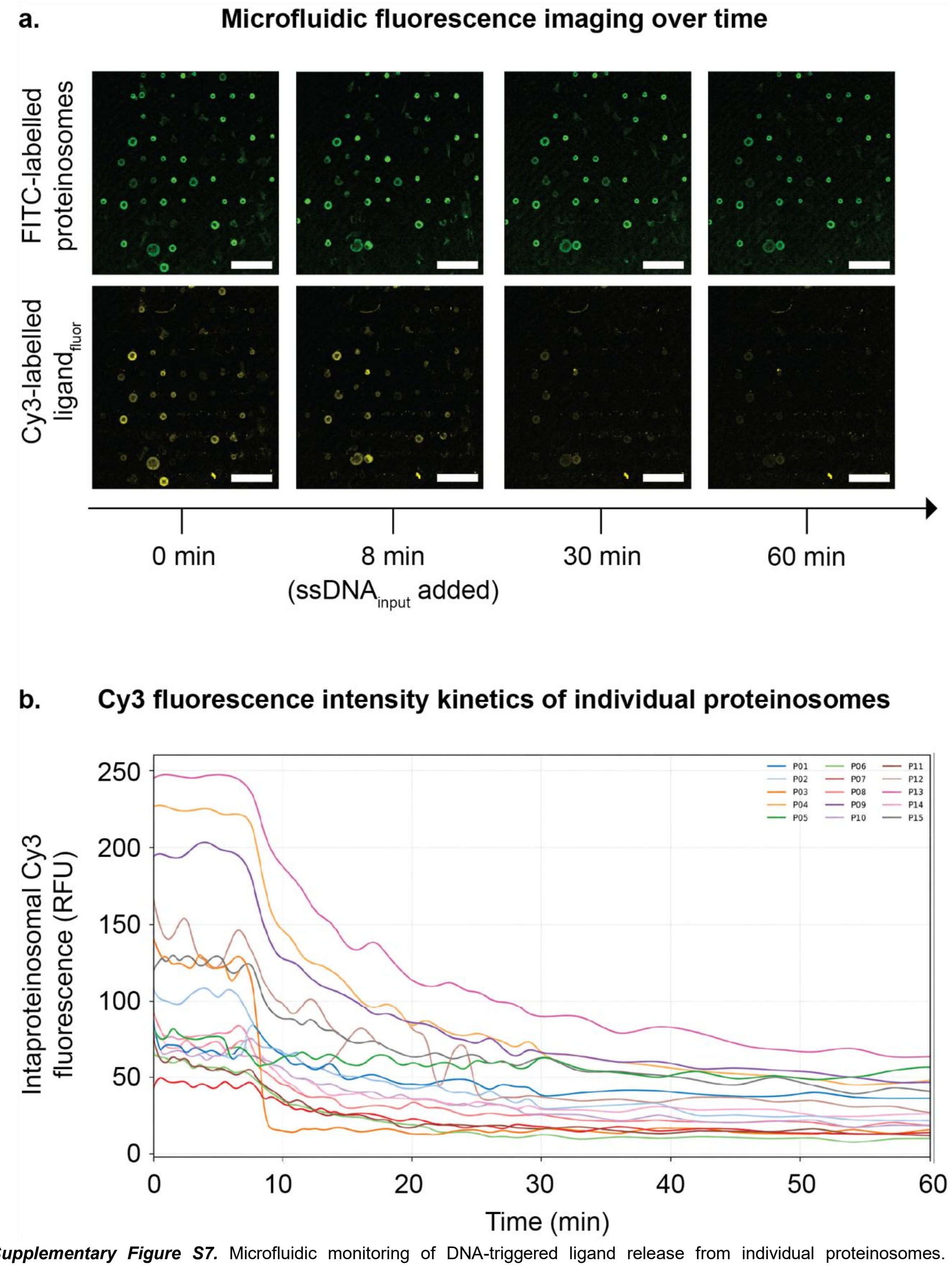


***Supplementary Figure S7.*** Microfluidic monitoring of DNA-triggered ligand release from individual proteinosomes. (a) Representative fluorescence micrographs. FITC-labelled proteinosomes containing Cy3-labelled ssDNA–CC signalling ligand (**ligand$_{fluor}$**) were imaged during the microfluidic release experiment (Fig. 1f; Supplementary Movie 1; Methods). The upper row shows FITC fluorescence used to identify proteinosomes, whereas the lower row shows Cy3 fluorescence corresponding to the localized ligand. The peripheral fluorescence pattern, with a central non-fluorescent region in both channels, reflects the internal bead architecture resulting from the central accumulation of non-fluorescent magnetic beads specifically used for this microfluidic experiment (Methods). Following addition of 100 nM complementary ssDNA (**ssDNA$_{input}$**) at $t$ = 8 min, intraproteinosomal Cy3 fluorescence decreased over time, consistent with DNA-triggered ligand release by toehold-mediated strand displacement (TMSD). (b) Individual fluorescence trajectories. Cy3 fluorescence trajectories are shown over time for the 15 proteinosomes included in the kinetic analysis. For quantification, Cy3 fluorescence was measured in the regions surrounding the central non-fluorescent bead-rich regions. Proteinosomes that moved substantially during the time series were excluded. Scale bars, 250 µm.

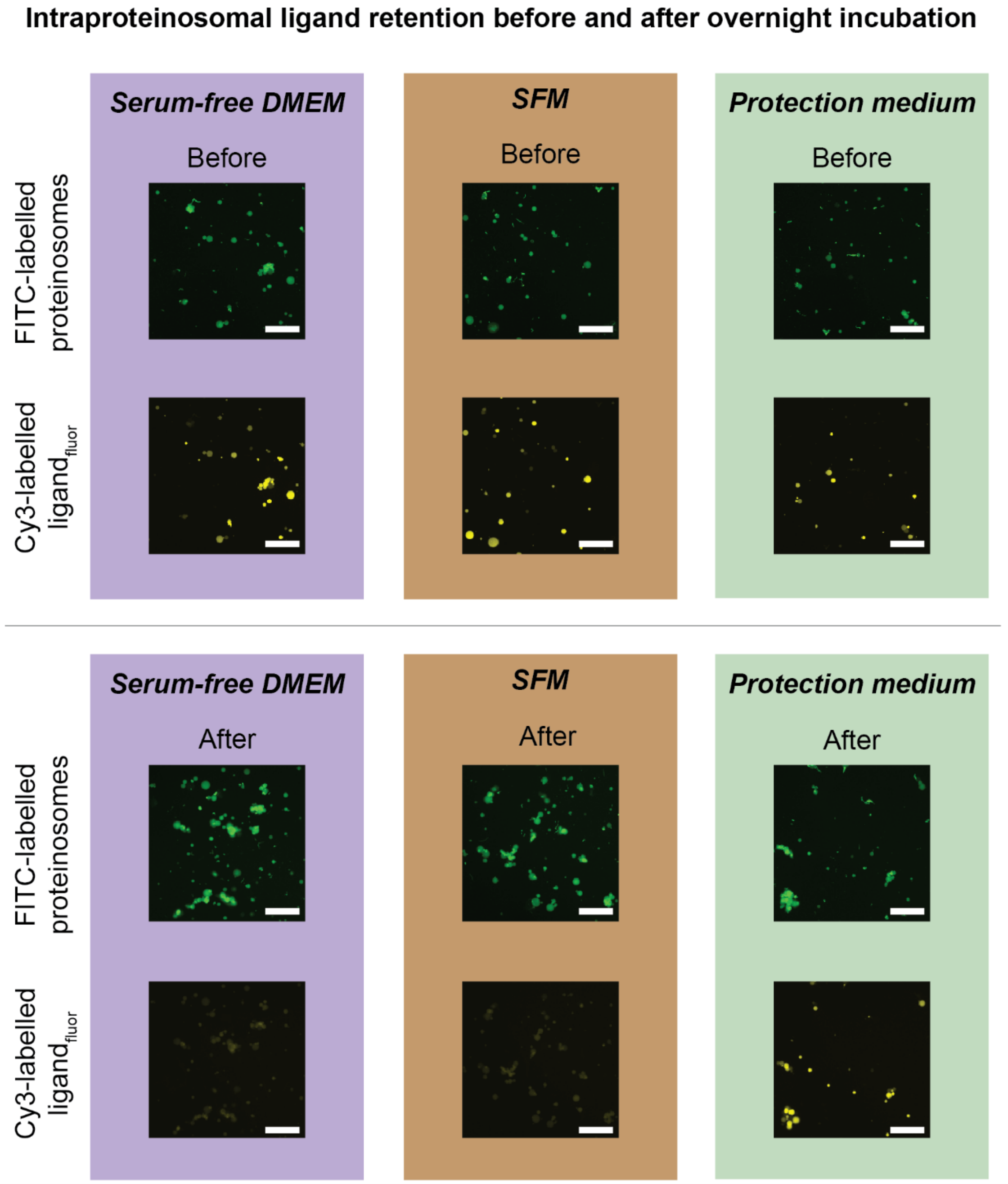


***Supplementary Figure S8.*** Representative fluorescence micrographs of intraproteinosomal ligand retention under mammalian cell culture conditions. FITC-labelled proteinosomes containing DNA-localized **ligand$_{fluor}$** were transferred into serum-free DMEM, serum-free 293 SFM II (SFM), or protection medium and imaged before and after overnight incubation under standard mammalian cell culture conditions (37 °C, 5% $CO_2$) (Methods). Protection medium consisted of DMEM containing 1.25% HI-FBS supplemented with 50% PBS and 1 µM **ssDNA$_{scramble}$**. Representative micrographs show FITC-labelled proteinosomes and intraproteinosomal Cy3 fluorescence from the DNA-localized ssDNA–CC signalling ligand before and after incubation. Before incubation, Cy3 fluorescence was readily detected within proteinosomes in all three media. Following overnight incubation, intraproteinosomal Cy3 fluorescence was strongly reduced in serum-free DMEM and SFM, whereas the Cy3 signal was largely retained in protection medium. Quantitative analysis in Fig. 2d showed a 3.2-fold and 3.4-fold reduction in intraproteinosomal Cy3 fluorescence in serum-free DMEM and SFM, respectively, whereas protection medium resulted in only a 1.1-fold decrease. The preservation of intraproteinosomal Cy3 fluorescence in protection medium supports maintenance of DNA-mediated ligand localization under mammalian cell culture conditions and further supports that **ssDNA$_{scramble}$** does not trigger substantial TMSD-mediated ligand release. A single independent experiment comprising all three medium conditions is shown, with the same dataset used for the quantitative analysis in Fig. 2d. The numbers of individual proteinosomes analysed were DMEM, n = 29 before and n = 30 after incubation; SFM, n = 23 before and n = 30 after incubation; and protection medium, n = 19 before and n = 30 after incubation. Images before and after incubation were acquired from different fields of view. Up to 30 individually resolvable proteinosomes were analysed per condition and time point. Fold reductions were calculated as the ratio of mean pre-incubation fluorescence to mean post-incubation fluorescence. Cy3 images were acquired and displayed using identical settings throughout. Scale bar, 250 µm.

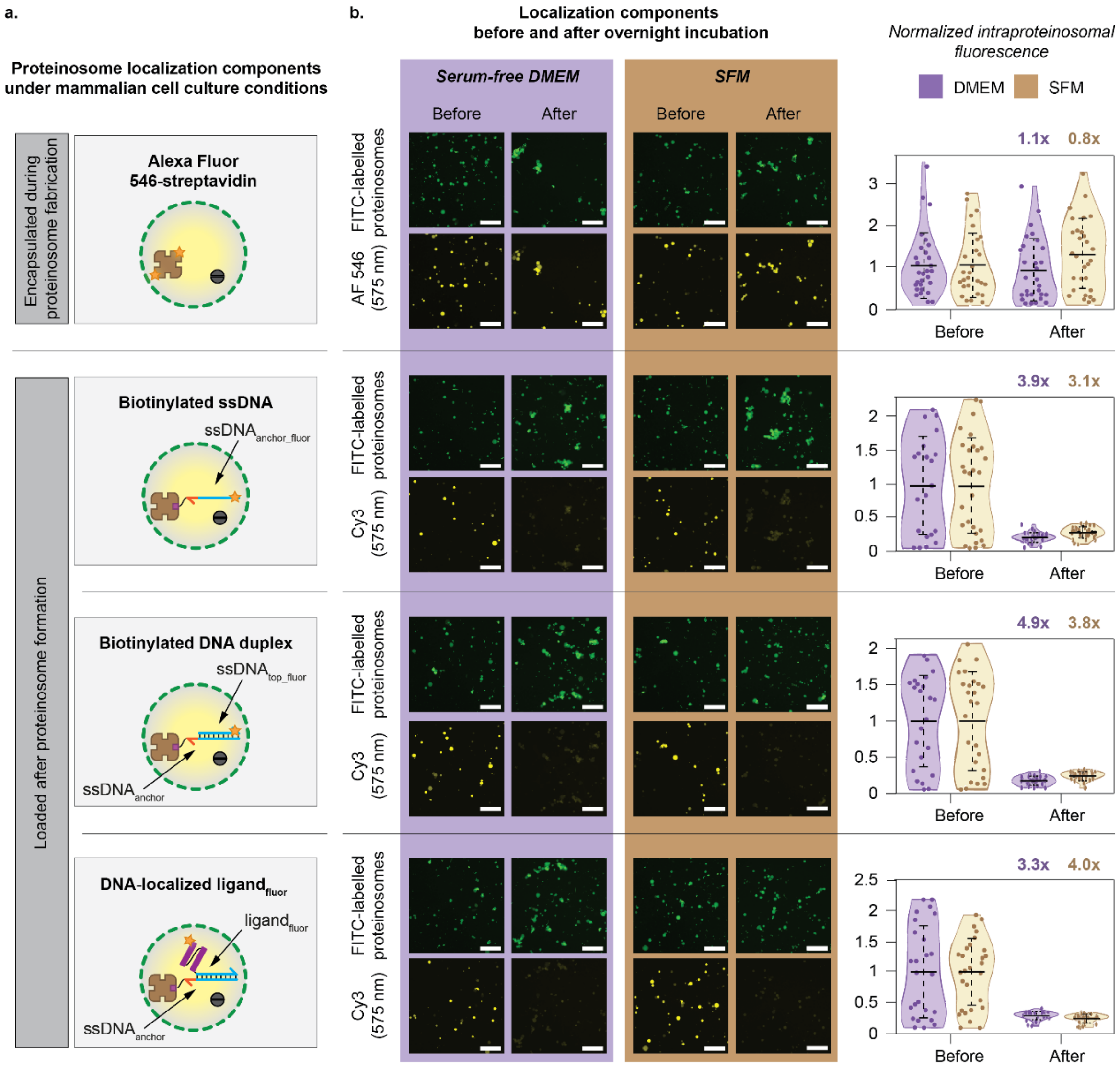


***Supplementary Figure S9.*** Characterization of proteinosome localization components under mammalian cell culture conditions. (a) Experimental design for characterizing retention of proteinosome localization components under mammalian cell culture conditions. Separate proteinosome populations were prepared to independently monitor retention of encapsulated streptavidin and the different components of the biotin- and DNA-mediated localization mechanism (Fig. 2c). Alexa Fluor 546 (AF 546)-labelled streptavidin was encapsulated during proteinosome formation to monitor retention of the streptavidin component (Methods). The other proteinosome populations contained unlabelled encapsulated streptavidin and were loaded with biotinylated DNA constructs to generate biotinylated ssDNA, biotinylated DNA duplex, or DNA-localized **ligand$_{fluor}$**. Biotinylated ssDNA consisted of Cy3-labelled 34-nt **ssDNA$_{anchor_fluor}$** immobilized within proteinosomes via streptavidin–biotin interactions, biotinylated DNA duplex was generated by immobilizing 34-nt **ssDNA$_{anchor}$** followed by hybridization with a Cy3-labelled 26-nt **ssDNA$_{top_fluor}$** strand, and DNA-localized **ligand$_{fluor}$** was generated similarly using **ligand$_{fluor}$** as the hybridization partner. The resulting populations were incubated overnight at 37 °C and 5% $CO_2$ in serum-free DMEM or serum-free 293 SFM II (SFM) (Methods). (b) Representative fluorescence micrographs showing FITC-labelled proteinosomes and corresponding intraproteinosomal fluorescence before and after overnight incubation in serum-free DMEM or SFM. Quantification at right shows the fluorescence of individual proteinosomes normalized to the corresponding mean pre-incubation fluorescence. Fluorescence of encapsulated, fluorescently labelled streptavidin remained relatively stable during overnight incubation, with pre-/post-incubation fluorescence ratios of 1.1 and 0.8 in serum-free DMEM and SFM, respectively. In contrast, fluorescence decreased 3.9-fold and 3.1-fold for biotinylated ssDNA, 4.9-fold and 3.8-fold for biotinylated DNA duplex, and 3.3-fold and 4.0-fold for **ligand$_{fluor}$** in serum-free DMEM and SFM, respectively. Thus, encapsulated streptavidin was largely retained, whereas biotinylated DNA cargo and the complete DNA-localized ligand showed substantially greater fluorescence loss. Retention of encapsulated streptavidin together with loss of biotinylated DNA cargo is consistent with destabilization of the biotin-dependent anchoring mechanism under these conditions, providing a component-level correlate of the reduced ligand retention observed in Fig. 2d and Fig. S8. A single independent experiment comprising all localization components in both media is shown. Individual proteinosome counts before/after incubation were 30/30, 21/30, 24/30, and 26/30 for streptavidin, ssDNA, dsDNA, and **ligand$_{fluor}$**, respectively, in serum-free DMEM, and 28/30, 30/30, 27/30, and 27/30, respectively, in SFM. Images before and after incubation were acquired from different fields of view. Up to 30 individually resolvable proteinosomes were analysed per condition and time point. Within each fluorescence channel, images were acquired and displayed using identical settings across conditions and time points. Data are presented as violin plots showing individual proteinosomes. Horizontal lines indicate the mean, and error bars denote s.d. Fold reductions were calculated as the ratio of mean pre-incubation fluorescence to mean post-incubation fluorescence. Scale bar, 250 μm.

#### Anti-SUMO Western blot of ssDNA–CC ligand integrity during receptor activation

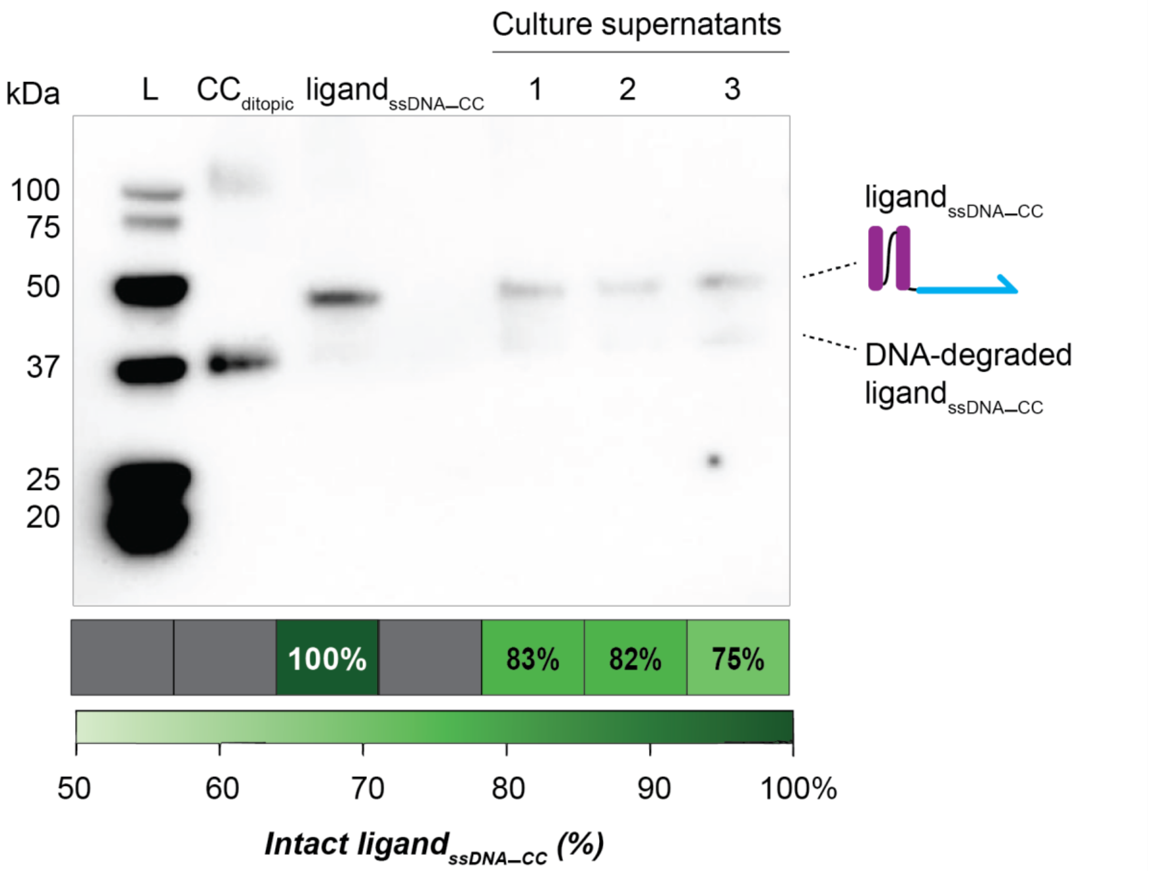


***Supplementary Figure S10.*** Anti-SUMO Western blot analysis of ssDNA–CC signalling ligand integrity during receptor activation. Culture supernatants from AA CC–GEMS-expressing HEK293T receiver cells incubated overnight with 120 nM **ligand**$_{\mathbf{ssDNA–CC}}$ in protection medium were analysed by anti-SUMO Western blotting to assess ligand integrity during receptor activation (Fig. 2e,f; Methods). Protection medium consisted of DMEM containing 1.25% HI-FBS supplemented with 50% PBS and 1 µM **ssDNA**$_{\mathbf{scramble}}$. L, molecular-mass marker; **CC**$_{\mathbf{ditopic}}$, protein stock; **ligand**$_{\mathbf{ssDNA–CC}}$, ssDNA–CC conjugate stock; 1–3, biological replicates from the receptor-activation experiment in Fig. 2f. The upper band corresponds to intact **ligand**$_{\mathbf{ssDNA–CC}}$, whereas the lower band is consistent with a DNA-degraded ligand species that co-migrates with **CC**$_{\mathbf{ditopic}}$. An additional band observed at approximately 25 kDa in replicate 3 was considered nonspecific background and was not included in the ligand-integrity analysis. Relative ligand integrity was determined by densitometric analysis of the intact and DNA-degraded ligand bands, calculated as intact ligand/(intact ligand + DNA-degraded ligand) × 100. The intact ligand fraction ranged from 75–83% across the three biological replicates. Greater loss of intact ligand was observed than in the corresponding cell-free stability experiment in Fig. 2b, consistent with increased nuclease exposure in the presence of HEK293T cells. Together with the receptor-activation measurements in Fig. 2f, these data show that a substantial fraction of the bifunctional ligand remains intact under the established communication-compatible conditions. The corresponding uncropped image is provided in Supplementary Fig. S24.

**Micrographs of DNA-programmable ligand release after overnight co-incubation**

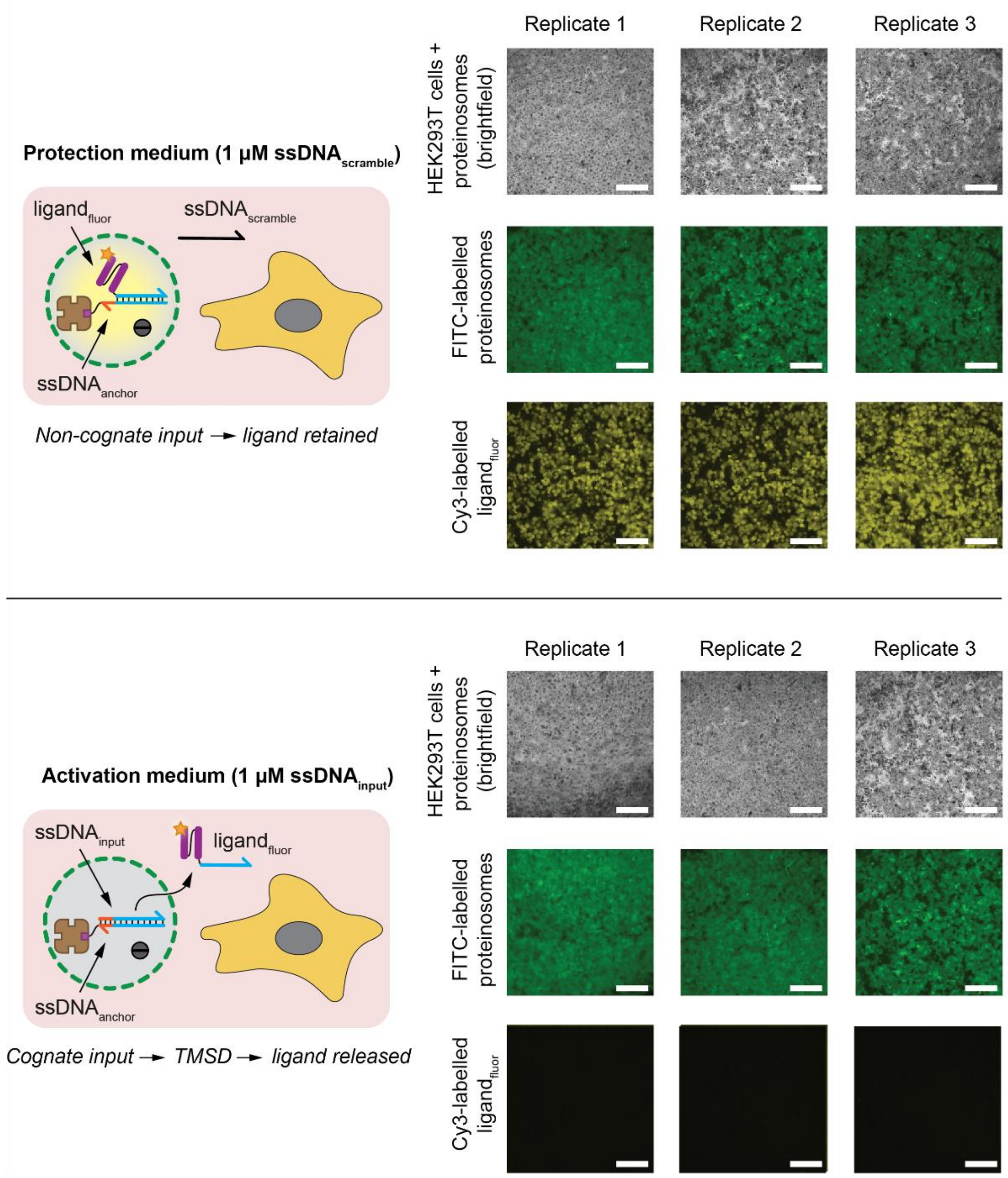


***Supplementary Figure S11.*** Representative fluorescence micrographs of DNA-programmable ligand release during proteinosome–mammalian cell co-incubation. HEK293T cells were co-incubated overnight with FITC-labelled proteinosomes containing DNA-localized **ligand$_{fluor}$** under standard mammalian cell culture conditions (37 °C, 5% $CO_2$) (Methods). Protection medium consisted of stability medium supplemented with 1 µM non-cognate **ssDNA$_{scramble}$**, whereas activation medium consisted of stability medium supplemented with 1 µM cognate **ssDNA$_{input}$**. The upper images show protection-medium co-incubation, in which **ssDNA$_{scramble}$** does not trigger TMSD and intraproteinosomal Cy3 fluorescence is retained. The lower images show activation-medium co-incubations, in which cognate **ssDNA$_{input}$** triggers TMSD-mediated ligand release and intraproteinosomal Cy3 fluorescence is strongly reduced. Brightfield images show the co-incubated HEK293T cells and proteinosomes, FITC fluorescence identifies the proteinosomes, and Cy3 fluorescence reports DNA-localized **ligand$_{fluor}$**. These observations support minimal ligand leakage during mammalian-cell co-incubation in protection medium while preserving cognate-input-triggered ligand release in activation medium. Quantitative analysis in Fig. 3b showed an 8.3-fold reduction in intraproteinosomal Cy3 fluorescence in activation medium relative to protection medium. Three independent biological replicates were analysed per condition, with fluorescence quantified in 500 grid-based image-sampling regions per replicate. The dense proteinosome population provided near-complete coverage of the sampled fields. Cy3 images were acquired and displayed using identical settings throughout. Scale bar, 250 µm.

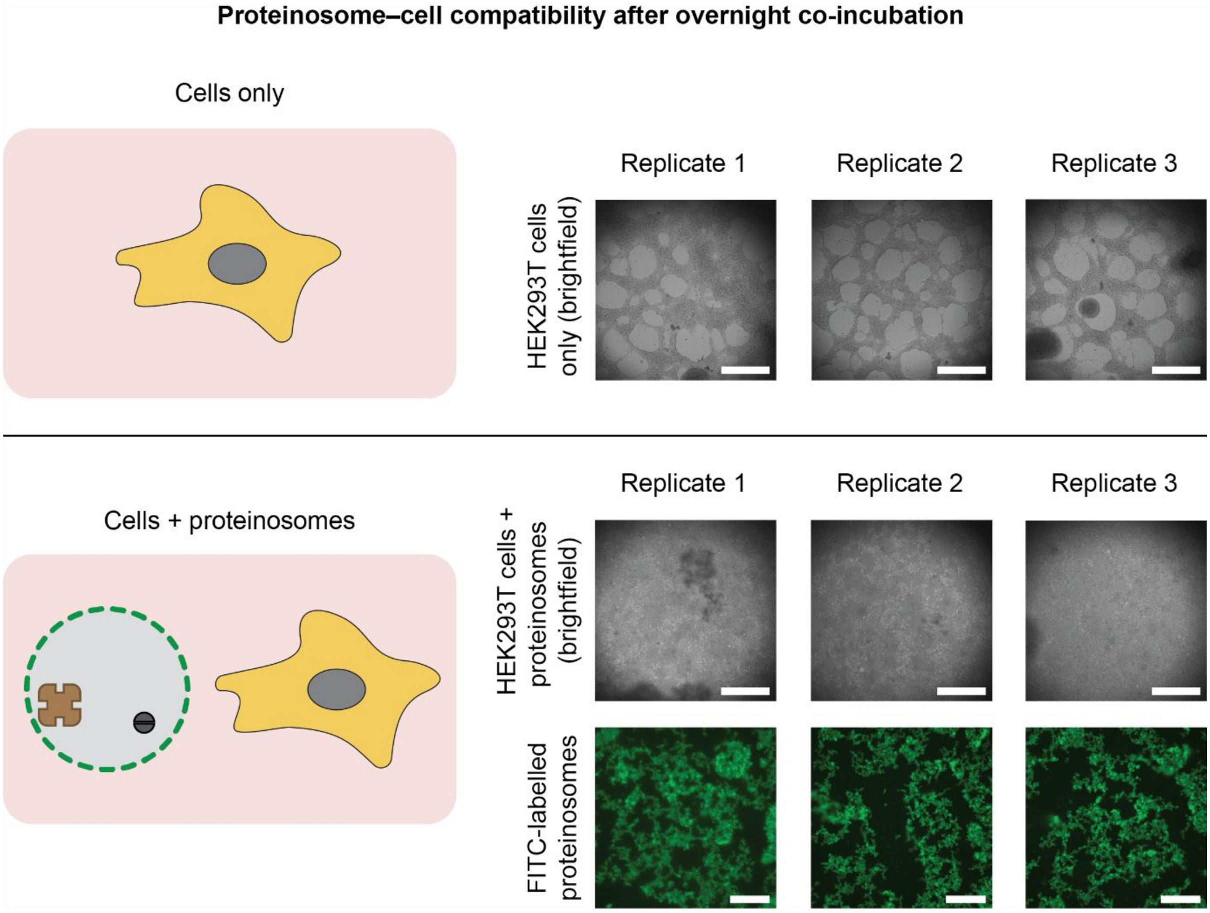


***Supplementary Figure S12.*** Representative micrographs corresponding to the proteinosome–mammalian cell compatibility analysis in Fig. 3c. HEK293T cells were incubated overnight in stability medium (DMEM containing 1.25% HI-FBS and 50% PBS), either alone or in the presence of FITC-labelled proteinosomes, under standard mammalian cell culture conditions (37 °C, 5% $CO_2$) (Methods). The upper images show HEK293T cells incubated in stability medium alone, whereas the lower images show HEK293T cells co-incubated with FITC-labelled proteinosomes. Brightfield images show the HEK293T cells under both conditions, while FITC fluorescence identifies the proteinosomes in the co-incubation condition. These representative micrographs provide visual context for the metabolic-activity measurements in Fig. 3c, where MTT analysis showed no substantial change in HEK293T cellular metabolic activity upon proteinosome co-incubation relative to cells incubated in stability medium alone. Three biological replicates are shown for each condition. Scale bar, 250 µm.

##### Estimation of proteinosome sender population based on target ligand concentration

**a.** ***Proteinosome size distribution***

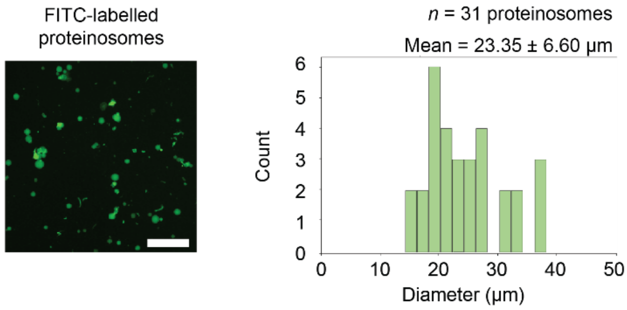


**b.** ***Intraproteinosomal ligand concentration***

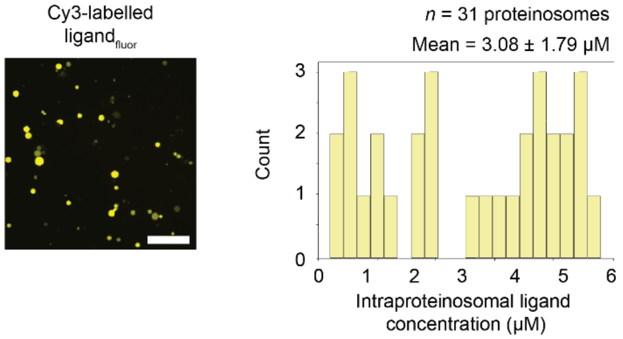


**c.** ***Calculation of required proteinosome number***

**Calculation approach**

1. Calculate individual proteinosome volumes and the mean proteinosome volume
   $V_i = {}^4/_3\pi \times (D_i / 2)^3$ ; $V_{proteinosome} = \text{mean}(V_i)$
2. Calculate ligand molecules per proteinosome
   $N_{ligand} = C_{ligand} \times V_{proteinosome} \times N_A$
3. Calculate total ligand molecules required
   $N_{total} = C_{target} \times V_{assay} \times N_A$
4. Calculate required number of proteinosomes
   $N_{proteinosomes} = N_{total} / (N_{ligand} \times \eta_{release})$

**Key parameters**

- Proteinosome dimensions
- Intraproteinosomal ligand concentration
- TMSD-release efficiency
- Target ligand concentration
- Assay volume

**Assumptions**

- Spherical proteinosomes
- Complete TMSD-release efficiency ($\eta_{release} = 1$)

***Supplementary Figure S13.*** Estimation of the proteinosome sender population required for DNA-programmable communication with mammalian receiver cells. (a) Proteinosome size distribution. Representative fluorescence micrograph of FITC-labelled proteinosomes (Fig. 1d) and corresponding distribution of proteinosome diameters, illustrating population heterogeneity. The population comprised n = *31* proteinosomes, with a mean diameter of 23.35 ± 6.60 µm (mean ± s.d.). (b) Intraproteinosomal ligand concentration. Representative fluorescence micrograph of **ligand$_{\mathbf{fluor}}$** localized within FITC-labelled proteinosomes (Fig. 1d; Methods) and corresponding distribution of intraproteinosomal ligand concentrations. Ligand concentration was determined from the Cy3 fluorescence signal using the calibration established in Fig. S4 and was measured as 3.08 ± 1.79 µM (mean ± s.d.; *n* = 31 proteinosomes). (c) Estimation of the required proteinosome sender population. Individual proteinosome volumes were calculated from the measured diameters assuming spherical geometry, and the resulting mean proteinosome volume was used to estimate ligand content per proteinosome. Individual proteinosome volumes were calculated according to $V_i = \frac{4}{3}\pi(D_i/2)^3$, where $D_i$ is the measured diameter of an individual proteinosome. The mean number of ligand molecules contained within a proteinosome was then calculated according to $N_{ligand} = C_{ligand} \times V_{proteinosome} \times N_A$, where $C_{ligand}$ is the mean intraproteinosomal ligand concentration, $V_{proteinosome}$ is the mean proteinosome volume calculated from the individual proteinosome volumes ($V_i$), and $N_A$ is Avogadro's constant. For a target ligand concentration of 120 nM in an assay volume of 100 µL, the total number of ligand molecules required was calculated according to $N_{total} = C_{target} \times V_{assay} \times N_A$, where $C_{target}$ is the target extracellular ligand concentration and $V_{assay}$ is the assay volume. The required number of ligand-containing proteinosomes was then calculated according to $N_{proteinosomes} = \frac{N_{total}}{(N_{ligand} \, x \, \eta_{release})}$, where $\eta_{release}$ is the TMSD-mediated ligand-release efficiency. The calculation assumes complete TMSD-mediated release of the intraproteinosomal ligand and therefore represents a nominal estimate of the sender population required to achieve the target extracellular ligand concentration. These parameters yielded an estimated requirement of approximately 4.69 × 10$^5$ ligand-containing proteinosomes, defining the sender population used for the subsequent proteinosome–mammalian cell communication experiments. Scale bar, 250 µm.

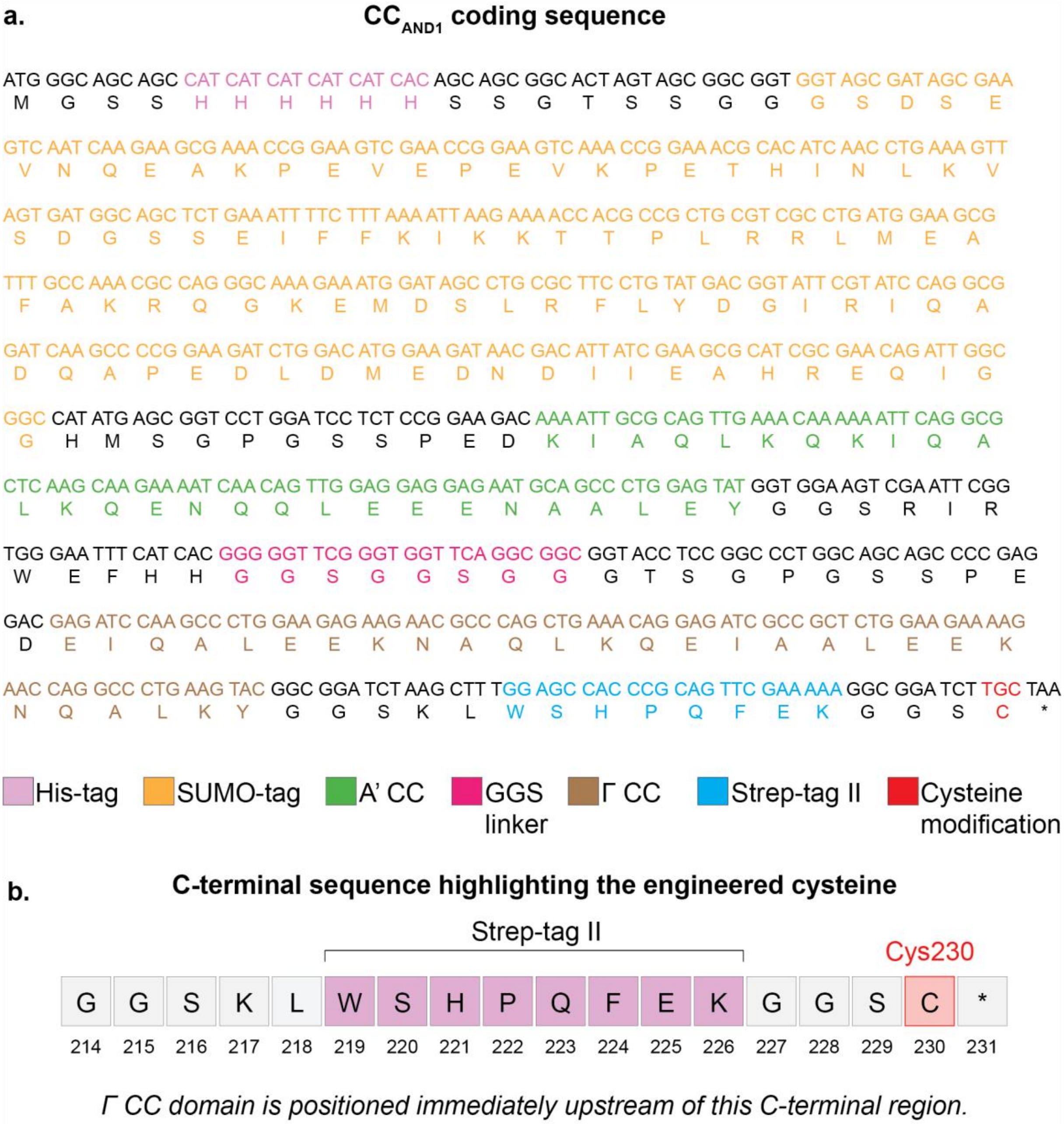


***Supplementary Figure S14.*** Design of the cysteine-engineered A'Γ $\textbf{CC}_{\textbf{AND1}}$ construct for cysteine-directed ssDNA conjugation. (a) Map of the A'Γ $\textbf{CC}_{\textbf{AND1}}$ coding sequence showing the N-terminal 6×His tag, SUMO domain, coiled-coil domains, GGS linker and C-terminal Strep-tag II. An engineered cysteine was introduced within the C-terminal region to provide a thiol handle for subsequent ssDNA conjugation. (b) C-terminal amino-acid sequence surrounding the engineered cysteine. Cys230 is positioned in the C-terminal region downstream of the Strep-tag II and provides a thiol handle for subsequent conjugation to ssDNA, generating $\textbf{ligand}_{\textbf{AND1}}$.

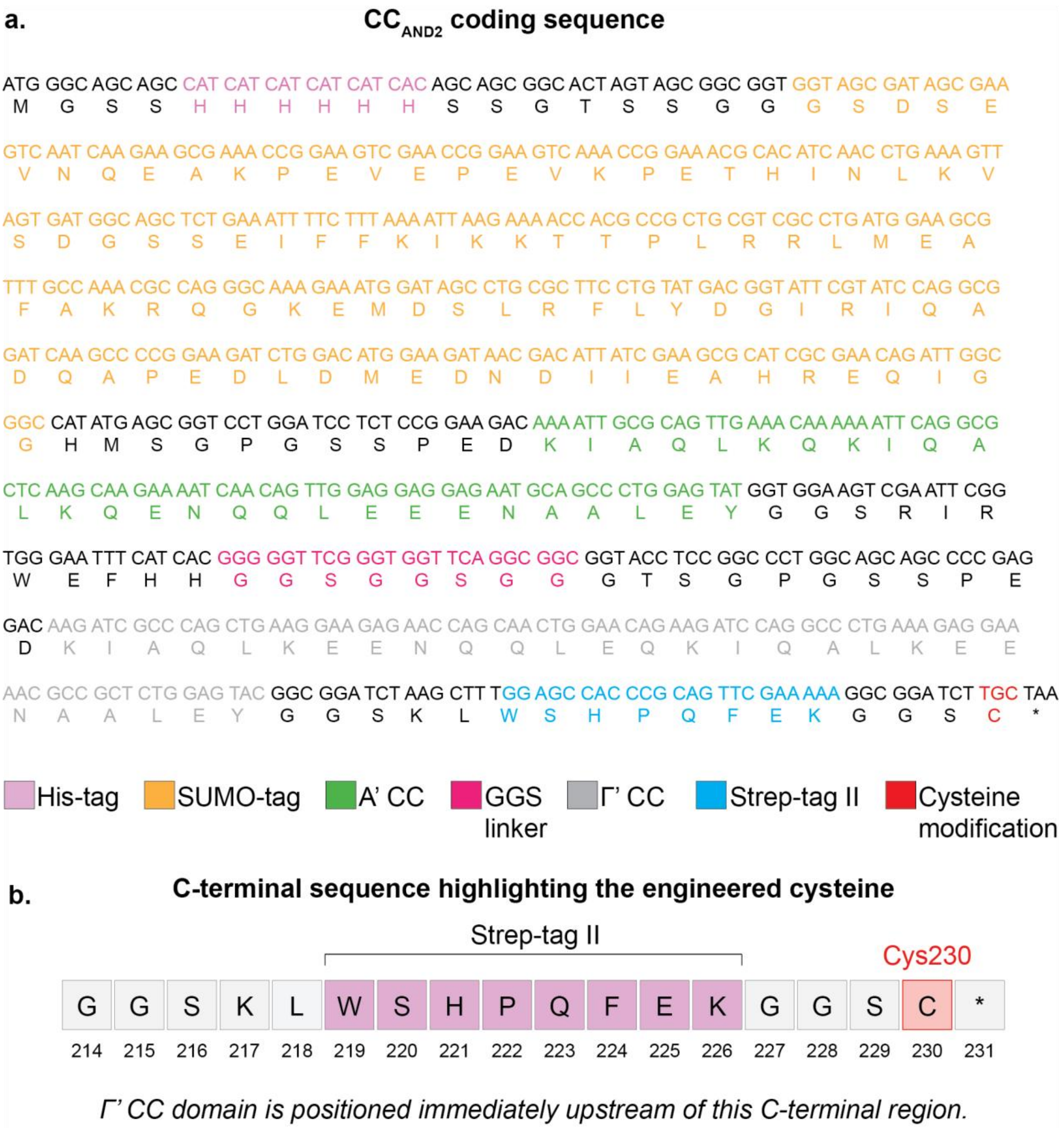


***Supplementary Figure S15.*** Design of the cysteine-engineered A'Γ' $\mathbf{CC_{AND2}}$ construct for cysteine-directed ssDNA conjugation. (a) Map of the A'Γ' $\mathbf{CC_{AND2}}$ coding sequence showing the N-terminal 6×His tag, SUMO domain, coiled-coil domains, GGS linker and C-terminal Strep-tag II. An engineered cysteine was introduced within the C-terminal region to provide a thiol handle for subsequent ssDNA conjugation. (b) C-terminal amino-acid sequence surrounding the engineered cysteine. Cys230 is positioned in the C-terminal region downstream of the Strep-tag II and provides a thiol handle for subsequent conjugation to ssDNA, generating $\mathbf{ligand_{AND2}}$.

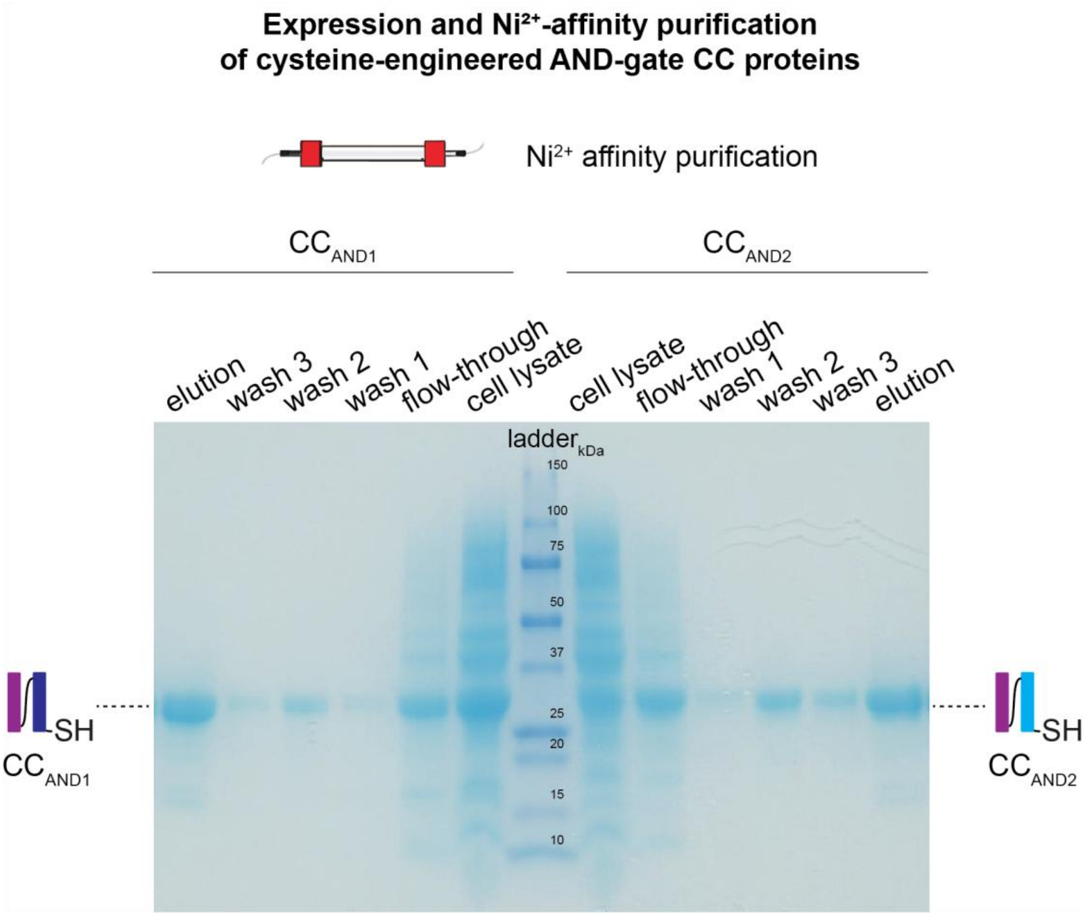


***Supplementary Figure S16.*** Expression and purification of cysteine-engineered **$CC_{AND1}$** and **$CC_{AND2}$** coiled-coil constructs. SDS–PAGE analysis of expression and $Ni^{2+}$-affinity purification of the cysteine-engineered **$CC_{AND1}$** and **$CC_{AND2}$** coiled-coil constructs under reducing, heated conditions (Methods). The expression and purification fractions for **$CC_{AND1}$** are shown to the left and those for **$CC_{AND2}$** to the right of the molecular-mass marker. The presence of protein bands in the elution fractions is consistent with recovery of the cysteine-engineered coiled-coil constructs following $Ni^{2+}$-affinity purification, with both coiled-coil proteins migrating at an apparent molecular mass of approximately 30 kDa. The purified constructs were used for subsequent conjugation to their corresponding ssDNA strands to generate the ligand components required for distributed AND-gated signalling (Fig. S17; Methods). The corresponding uncropped image is provided in Supplementary Fig. S22.

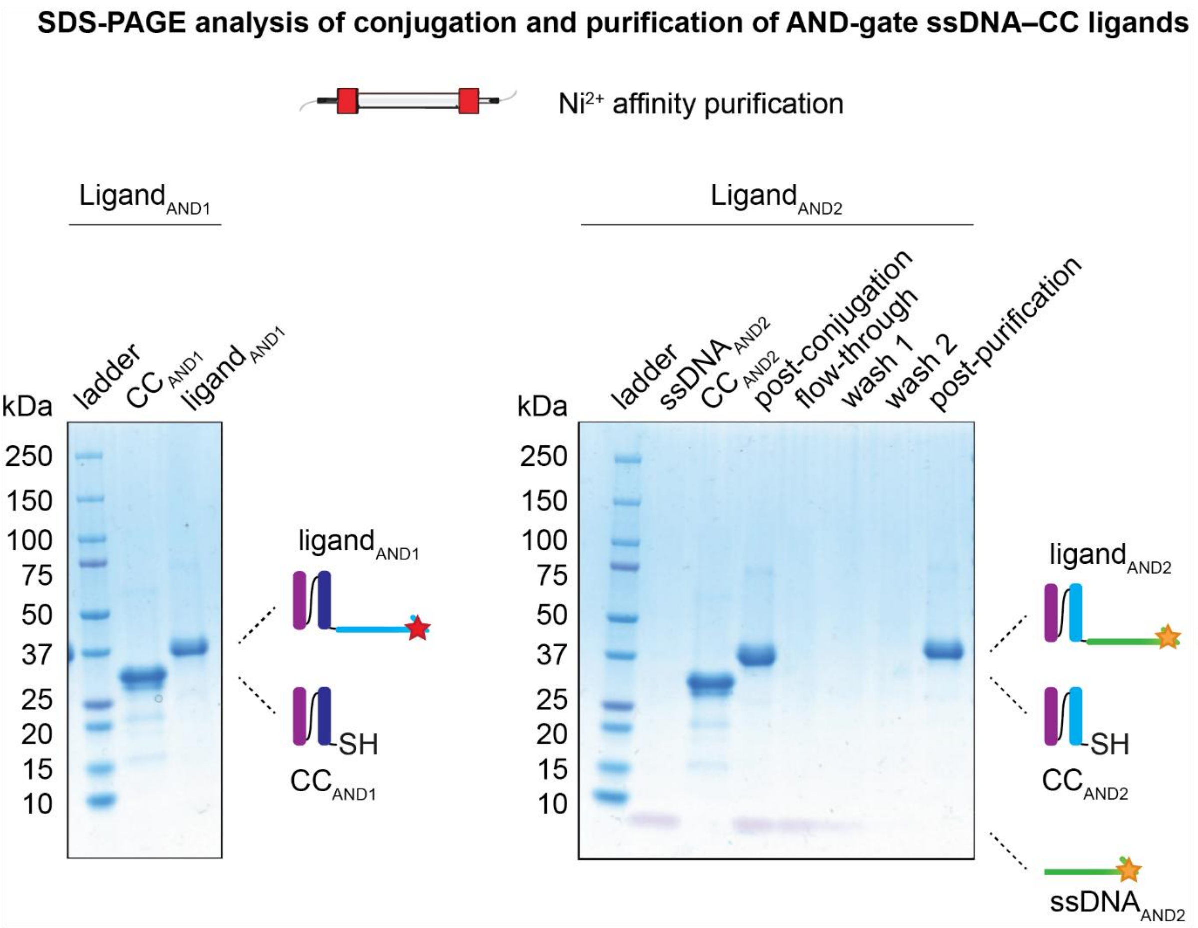


***Supplementary Figure S17.*** Validation of ssDNA–CC conjugation and purification of the AND-gate ligand components. SDS-PAGE analysis of the conjugation and purification of the ssDNA–CC ligand components **ligand$_{AND1}$** and **ligand$_{AND2}$** (Methods). The respective cysteine-engineered coiled-coil proteins were conjugated to their corresponding maleimide-functionalized ssDNA strands through thiol–maleimide coupling. The ssDNA component of **ligand$_{AND1}$** was Cy5-labelled (26-nt, **ssDNA$_{AND1}$**), whereas the ssDNA component of **ligand$_{AND2}$** was Cy3-labelled (26-nt, **ssDNA$_{AND2}$**). For **ligand$_{AND1}$** (left), **CC$_{AND1}$** protein before conjugation (**CC$_{AND1}$**) and the **ligand$_{AND1}$** elution fraction following ssDNA conjugation and $Ni^{2+}$-affinity purification are shown (**Ligand$_{AND1}$**). For **ligand$_{AND2}$** (right), the Cy3-labelled **ssDNA$_{AND2}$** and **CC$_{AND2}$** starting materials, the post-conjugation sample, flow-through, wash fractions 1 and 2, and purified elution fraction (post-purification) are shown. The changes in electrophoretic migration following conjugation are consistent with formation of the corresponding ssDNA–CC ligand components, while the purification fractions show enrichment of the conjugated products and depletion of unbound ssDNA. The corresponding uncropped image is provided in Supplementary Fig. S22.

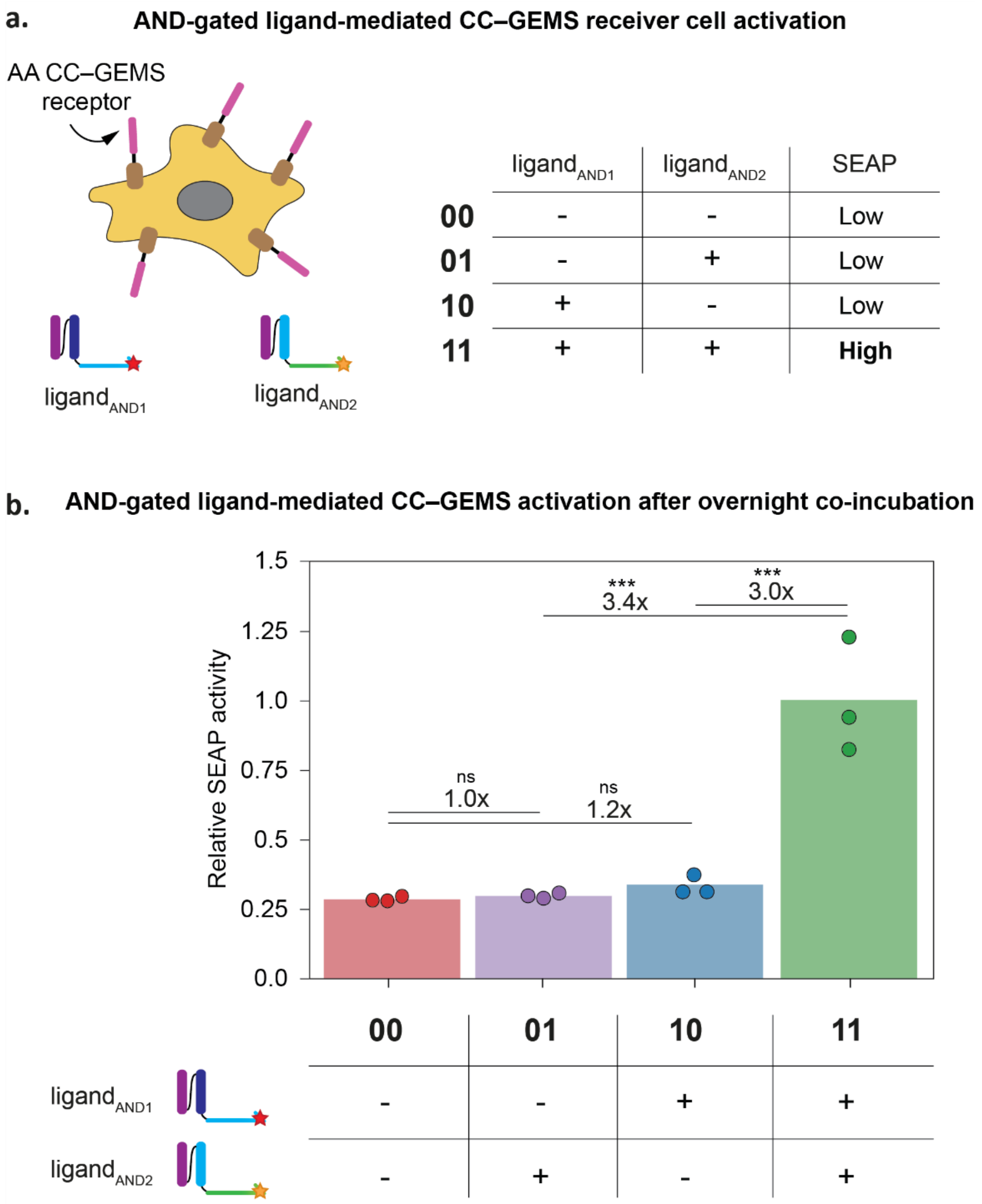


***Supplementary Figure S18.*** CC–GEMS activation by complementary ligand components in an AND-gate configuration. (a) Schematic representation of AND-gated ligand-mediated CC–GEMS receptor activation. The complementary ligand components **ligand$_{AND1}$** and **ligand$_{AND2}$** engage receptor signalling only when both components are simultaneously present, allowing formation of the functional ligand pair required for receptor activation.[33] (b) Quantification of CC–GEMS activation following overnight incubation of HEK293T receiver cells expressing AA CC–GEMS receptors with the complementary ligand components in four configurations in protection medium with 2 µM **ssDNA$_{scramble}$**: absence of both ligand components (00), 120 nM **ligand$_{AND2}$** alone (01), 120 nM **ligand$_{AND1}$** alone (10), or 120 nM each of **ligand$_{AND1}$** and **ligand$_{AND2}$** (11) (Methods). SEAP activity was measured in the culture supernatants and normalized to the 11-condition. The individual ligand components produced low SEAP activity when administered separately, comparable to the condition lacking both components, with 01 and 10 showing 1.0-fold and 1.2-fold changes relative to 00, respectively. Simultaneous addition of **ligand$_{AND1}$** and **ligand$_{AND2}$** resulted in increased SEAP activity, corresponding to 3.4-fold and 3.0-fold increases relative to the 01 and 10 conditions, respectively. These results demonstrate that complementary ligand components are required for functional CC–GEMS activation and establish the basis for implementing distributed AND-gated signalling using independently programmed proteinosome populations. Data represent biological triplicates; bars indicate the mean and dots indicate individual replicates. Statistical significance was determined by one-way ANOVA across all experimental conditions, with Tukey's multiple-comparison test. ns: $P > 0.05$, *$P \leq 0.05$, **$P \leq 0.01$, ***$P \leq 0.001$, ****$P \leq 0.0001$.

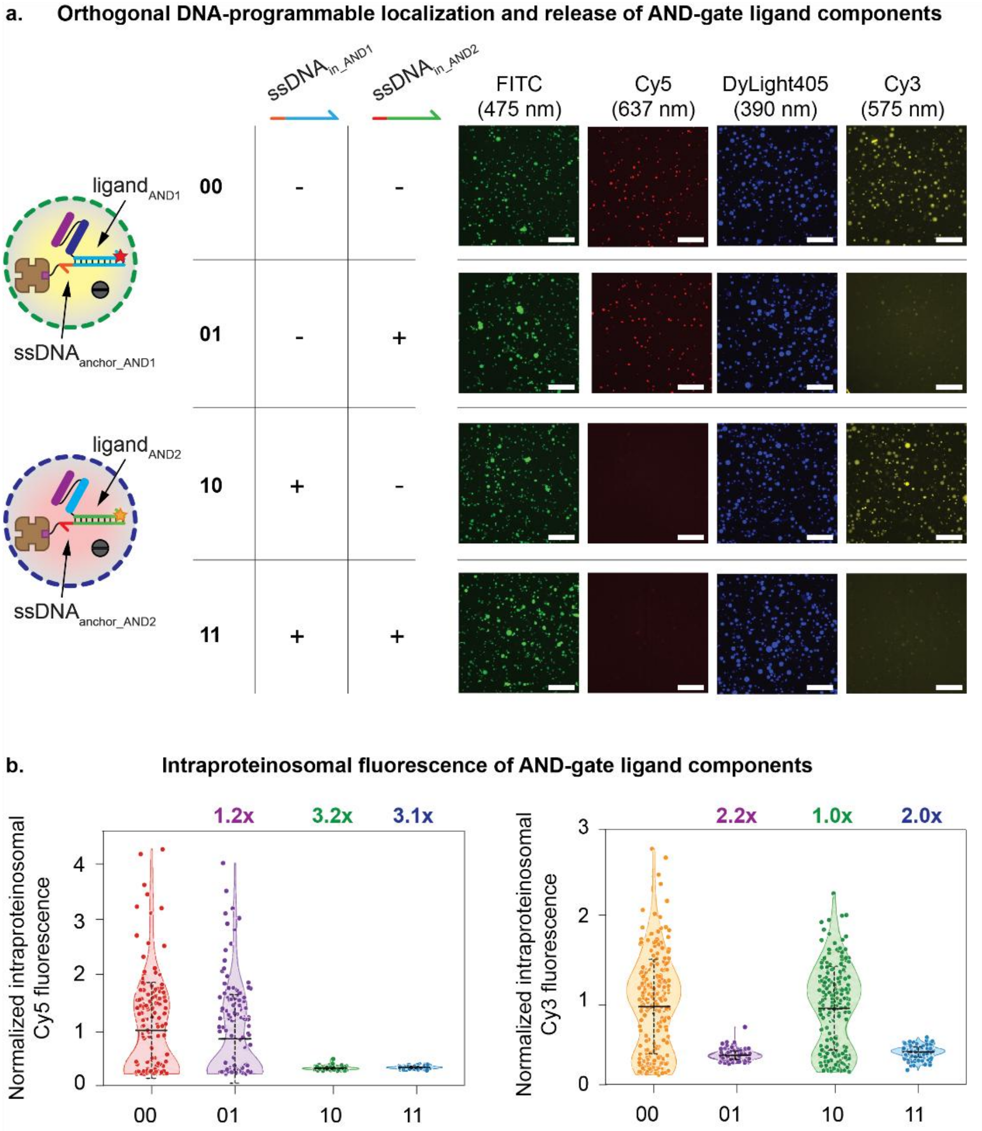


***Supplementary Figure S19.*** Orthogonal DNA-programmable localization and release of AND-gate ligand components from proteinosomes. (a) Experimental design and representative fluorescence micrographs illustrating orthogonal DNA-programmable localization and release of **ligand$_{AND1}$** and **ligand$_{AND2}$** from separate proteinosome populations. Population **A**, labelled with FITC, contained the Cy5-labelled **ligand$_{AND1}$**, whereas population **B**, labelled with DyLight405, contained the Cy3-labelled **ligand$_{AND2}$**, both localized through their corresponding ssDNA anchors (Methods). The four input configurations consisted of the absence of both DNA inputs (00), addition of **ssDNA$_{in_AND2}$** alone (01), addition of **ssDNA$_{in_AND1}$** alone (10), or simultaneous addition of both DNA inputs (11). Representative fluorescence micrographs show the FITC- and DyLight405-labelled proteinosomes and the corresponding intraproteinosomal Cy5 and Cy3 fluorescence signals. In the absence of input DNA, both ligand components remained localized within their respective proteinosome populations. Addition of **ssDNA$_{in_AND1}$** or **ssDNA$_{in_AND2}$** selectively reduced the corresponding intraproteinosomal ligand fluorescence, whereas the non-cognate input did not produce a detectable reduction in fluorescence in the orthogonal proteinosome population, consistent with orthogonal, sequence-specific TMSD-mediated ligand release. Simultaneous addition of both inputs resulted in reduced intraproteinosomal fluorescence of both ligand components. (b) Quantification of intraproteinosomal Cy5 and Cy3 fluorescence associated with **ligand$_{AND1}$** and **ligand$_{AND2}$**, respectively, across the four input configurations. Fluorescence values were normalized to the 00 condition within each channel. Addition of **ssDNA$_{in_AND1}$** selectively reduced intraproteinosomal Cy5 fluorescence associated with **ligand$_{AND1}$**, corresponding to a 3.2-fold reduction relative to the 00 condition, whereas the non-cognate **ssDNA$_{in_AND2}$** resulted in a 1.2-fold change; simultaneous addition of both DNA inputs resulted in a 3.1-fold reduction. Similarly, addition of **ssDNA$_{in_AND2}$** selectively reduced intraproteinosomal Cy3 fluorescence associated with **ligand$_{AND2}$**, corresponding to a 2.2-fold change relative to the 00 condition, whereas the non-cognate **ssDNA$_{in_AND1}$** resulted in a 1.0-fold change; simultaneous addition of both DNA inputs resulted in a 2.0-fold reduction in Cy3 fluorescence. Together, these results demonstrate orthogonal, sequence-specific localization and TMSD-mediated release of the ligand components, providing the basis for distributed AND-gated communication between proteinosome sender populations and mammalian receiver cells (Fig. 4c). For Cy5-labelled **ligand$_{AND1}$**, 170, 186, 219 and 155 proteinosomes were analysed in the 00, 01, 10 and 11 conditions, respectively. For Cy3-labelled **ligand$_{AND2}$**, the corresponding numbers were 235, 204, 254 and 197 proteinosomes. Within each fluorescence channel, images were acquired and displayed using identical settings across input configurations. Data are presented as violin plots showing individual proteinosomes. Horizontal lines indicate the mean, and error bars denote s.d. Fold reductions were calculated as the ratio of mean fluorescence in the 00 condition to mean fluorescence in each input condition. Scale bar, 250 μm.

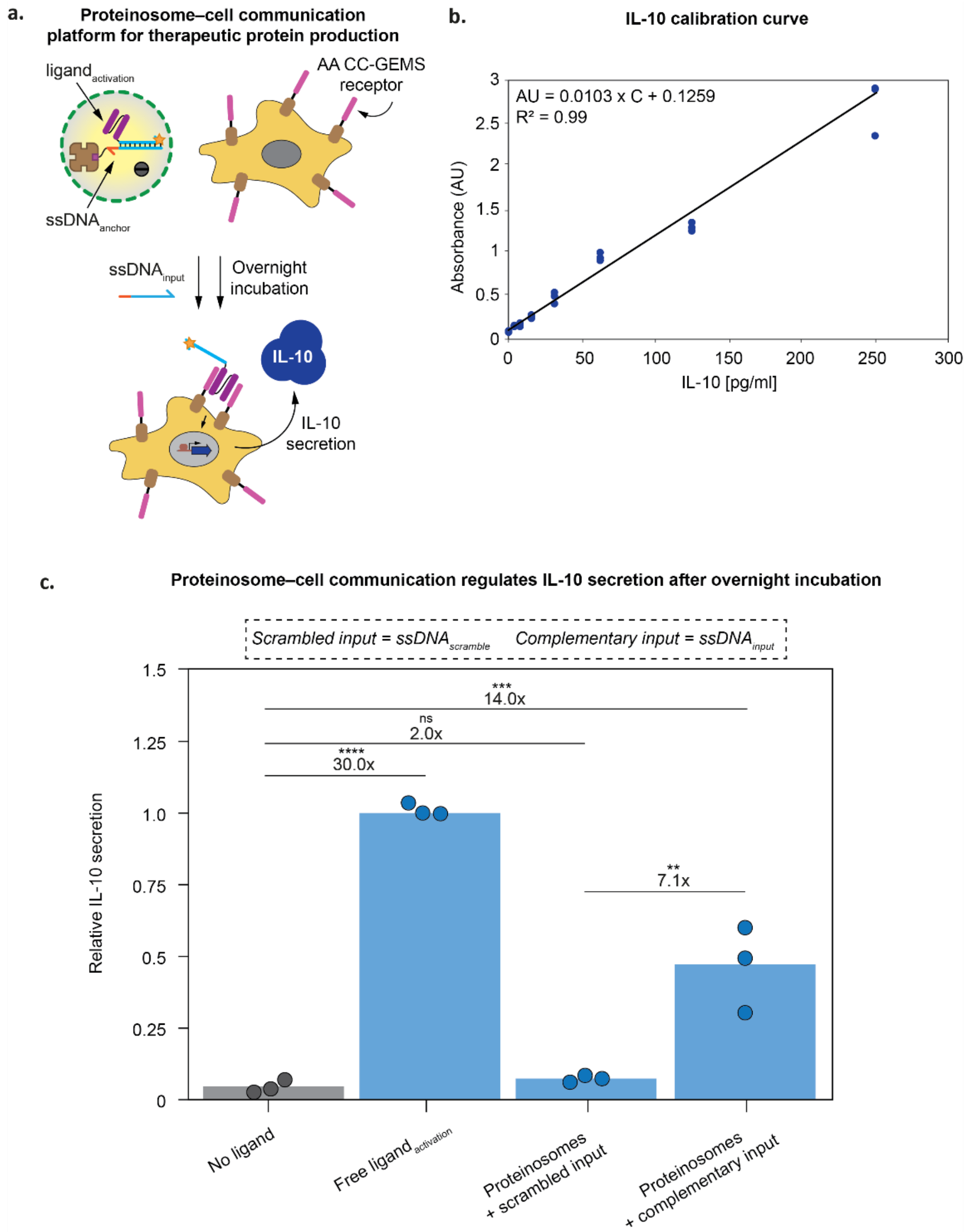


***Supplementary Figure S20.*** DNA-programmable proteinosome–mammalian cell communication regulates IL-10 secretion. (a) Assay for DNA-programmable proteinosome–mammalian cell communication for the regulation of interleukin-10 (IL-10) secretion. AA CC–GEMS-expressing HEK293T receiver cells, with an IL-10 output, were co-incubated overnight with ligand-containing proteinosomes in the presence of input DNA. Culture supernatants were subsequently collected, and IL-10 secretion was quantified colorimetrically by ELISA (Methods). (b) IL-10 calibration curve used to quantify IL-10 in culture supernatants by ELISA. Absorbance at 450 nm, with reference correction at 570 nm, was plotted against known IL-10 concentrations (0, 3.90625, 7.8125, 15.625, 31.25, 62.5, 125, and 250 pg/mL), and the data were fitted by linear regression. The resulting calibration equation was AU = 0.0103 × C + 0.1259 ($R^2$ = 0.99), where AU represents absorbance and C denotes IL-10 concentration (pg/mL). Individual datapoints represent independent technical replicates (*n* = 3). (c) DNA-programmable proteinosome–cell communication after overnight co-incubation. IL-10 secretion was measured in CC–GEMS receiver cells incubated without ligand or with 120 nM Cy3-labelled ssDNA–CC ligand (**ligand$_{\text{activation}}$**) under scrambled-input conditions, or with FITC-labelled proteinosomes containing **ligand$_{\text{activation}}$** under either scrambled-input or complementary-input conditions. IL-10 secretion was normalized to the free-ligand condition. Free ligand induced a 30.0-fold increase in IL-10 secretion relative to the no-ligand control, whereas proteinosomes under complementary-input conditions elicited a 14.0-fold increase relative to the no-ligand control. Complementary-input conditions produced significantly higher IL-10 secretion than scrambled-input conditions, with IL-10 secretion increasing 7.1-fold. The lower IL-10 secretion obtained with DNA-triggered proteinosomes relative to free ligand may reflect heterogeneous ligand loading within the proteinosome population (Fig. 1d; Fig. S13). Nevertheless, these results demonstrate that DNA-triggered ligand release from proteinosomes can regulate an alternative mammalian signalling output in addition to SEAP. Data represent biological triplicates; bars indicate the mean and dots indicate individual replicates. Statistical significance was determined by one-way ANOVA across all experimental conditions, with Tukey's multiple-comparison test. ns: P> 0.05, *P ≤ 0.05, **P ≤ 0.01, ***P ≤ 0.001, ****P ≤ 0.0001.

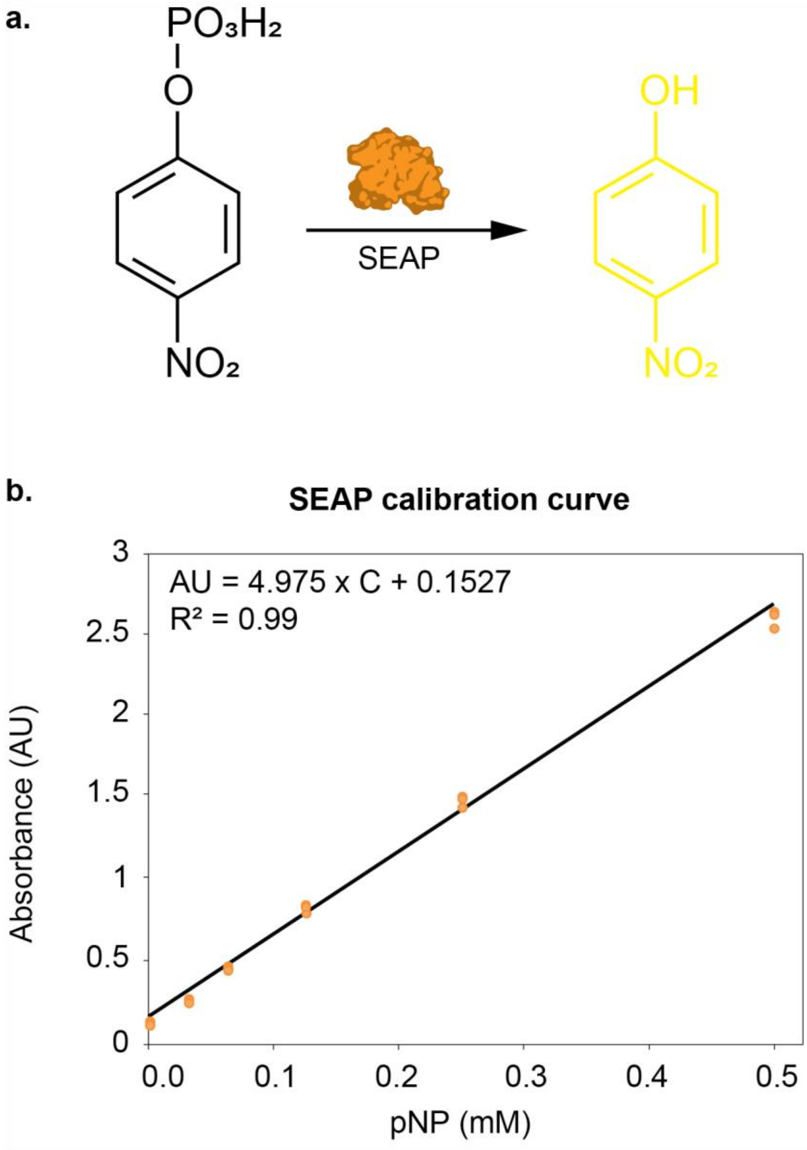


***Supplementary Figure S21***. Calibration curve used to quantify SEAP activity in culture supernatants. (a) SEAP catalyses the hydrolysis of the chromogenic substrate *p*-nitrophenyl phosphate (pNPP) to *p*-nitrophenol (pNP), which can be quantified by measuring absorbance at 405 nm. (b) Absorbance at 405 nm was plotted against known pNP concentrations (0, 0.0315, 0.0625, 0.125, 0.25, and 0.5 mM), and the data were fitted by linear regression. The resulting calibration equation was AU = 4.975 × C + 0.1527 ($R^2$ = 0.99), where AU represents absorbance and C denotes pNP concentration (mM). Individual data points represent independent technical replicates (*n* = 3).

### Uncropped SDS–PAGE gels

**a.** Gel for Figure 1b

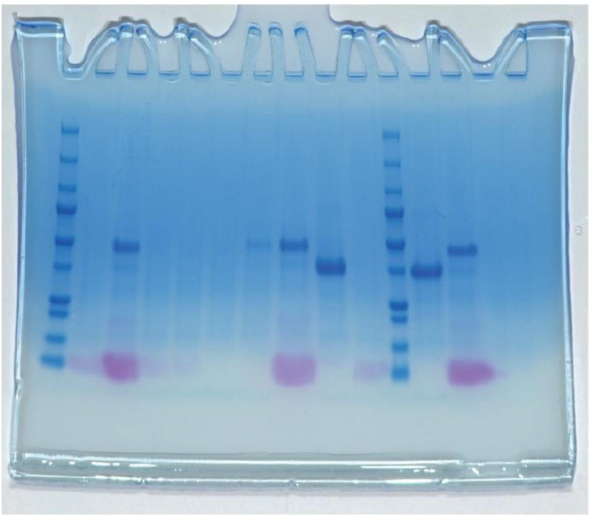

**b.** Gel for Figure S2

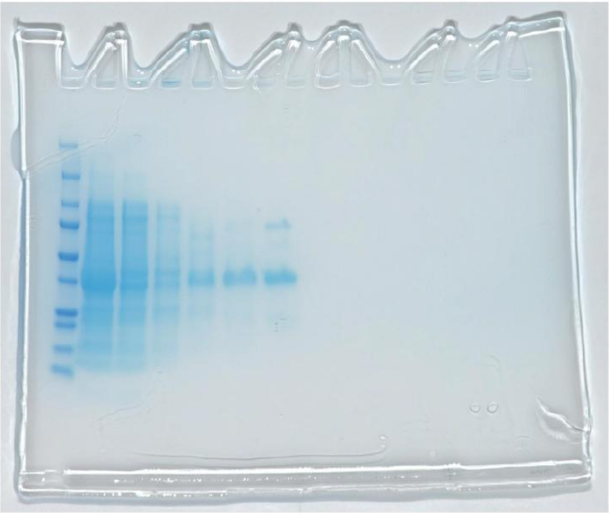

**c.** Gel for Figure S16

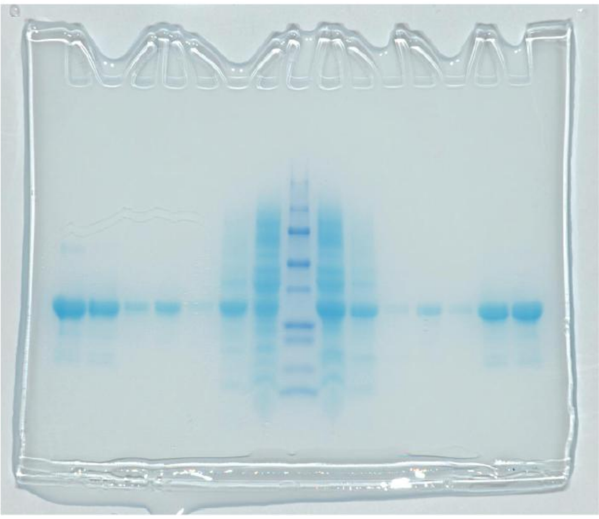

**d.** Gel for Figure S17

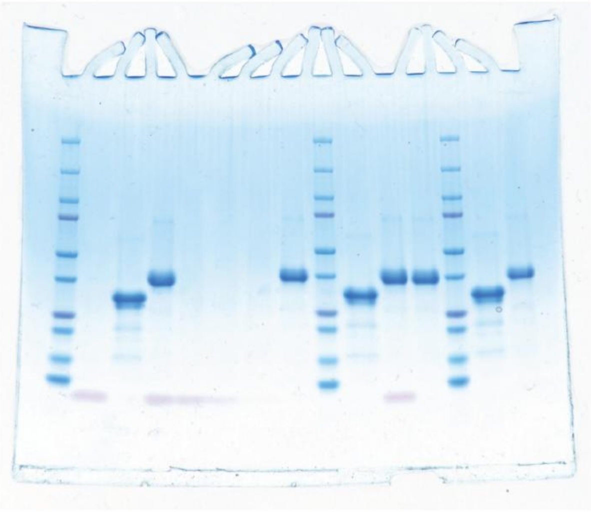

***Supplementary Figure S22***. Uncropped SDS–PAGE gels corresponding to Figure 1b and Supplementary Figures S2, S16 and S17.

#### Uncropped Native-PAGE gel for Figure S3

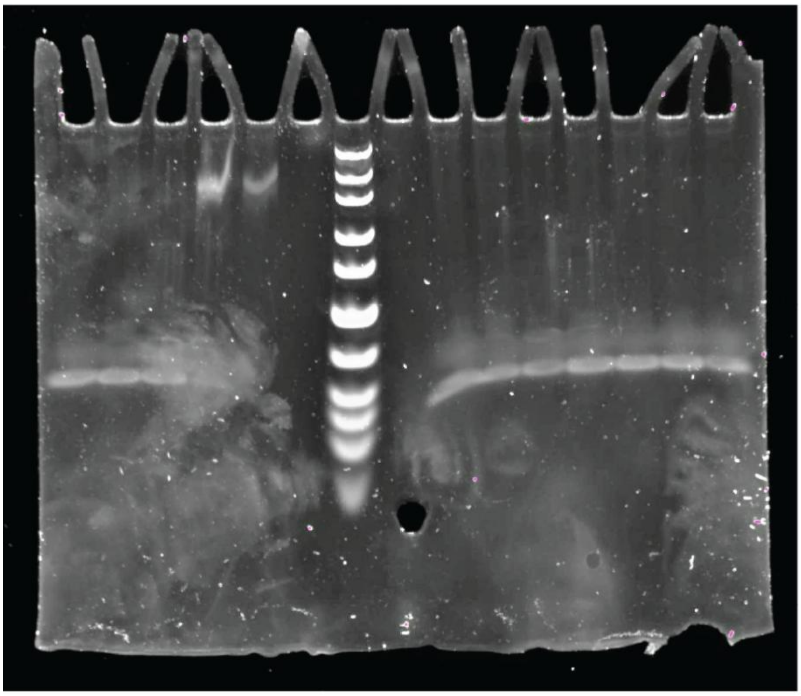

***Supplementary Figure S23***. Uncropped Native-PAGE gel corresponding to Supplementary Figure S3.

#### Uncropped Western blot membranes

**a.** Western blot for Figure 2b

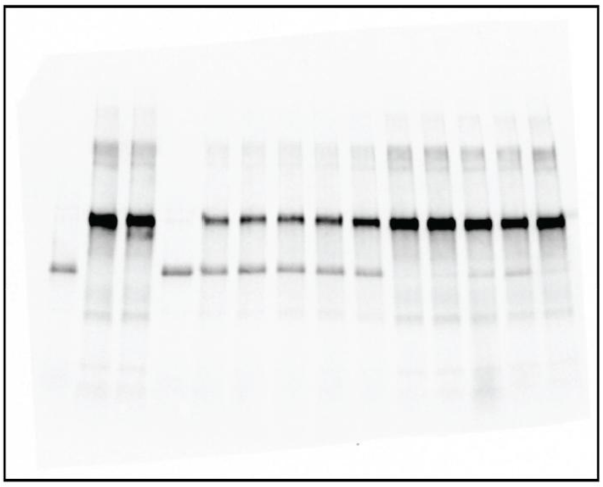

**b.** Western blot for Figure 3f

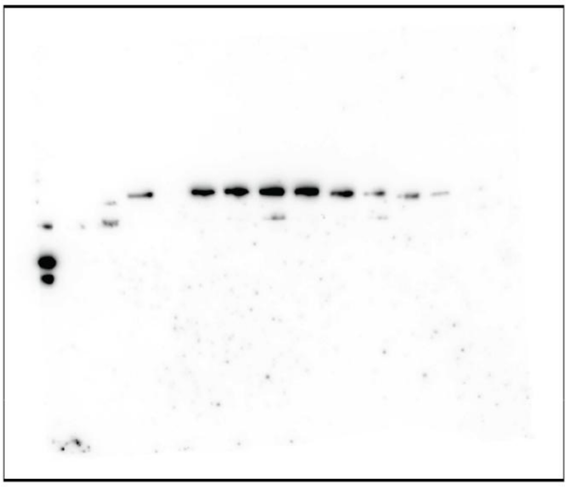

**c.** Western blot for Figure S10

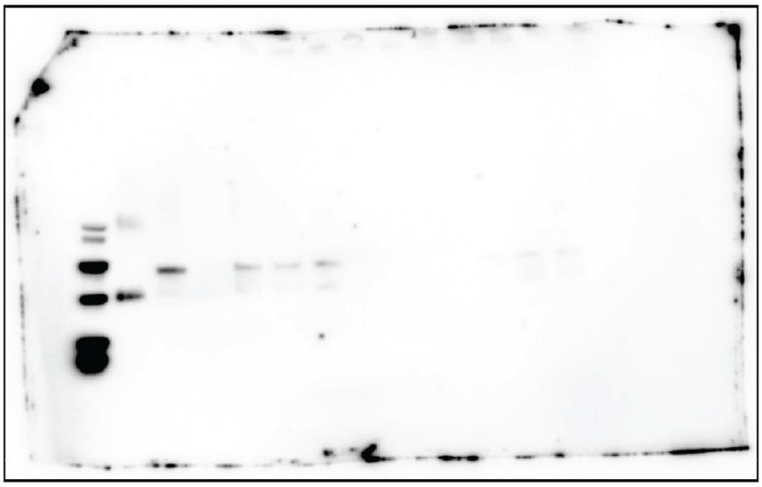

***Supplementary Figure S24***. Uncropped Western blot membranes corresponding to Figure 2b, Figure 3f and Supplementary Figure S10.

#### Supplementary tables

**Supplementary Table S1. Plasmid constructs.** Plasmids used in this study, including their backbone and encoded insert or function.

| Name | Description |
|---|---|
| pET28a(+)-$P_{T7}$ | Bacterial pET28a(+) expression vector used to express variants of SUMO–CC fusion proteins in *Escherichia coli (E. coli)*. |
| pLeo619-$P_{SV40}$ | Mammalian pLeo619 expression vector encoding the $EpoR_m$-IL-$6RB_m$-pA receptor region fused to the selected CC module. |
| pLeo-A-$\alpha_2$-$GEMS_{JAK/STAT}$ | Cognate AA CC–GEMS receptor construct. |
| pLeo-B-$\alpha_2$-$GEMS_{JAK/STAT}$ | Non-cognate BB CC–GEMS receptor construct. |
| pLS15 | Mammalian STAT3 expression vector ($P_{hCMV}$-STAT3-pA). |
| pLS13 | Mammalian reporter plasmid for STAT3-induced SEAP expression ($O_{Stat3}$-$P_{hCMVmin}$-SEAP-pA). |
| pLS13-IL-10 | Mammalian reporter plasmid for STAT3-induced IL-10 expression ($pLS13_{STAT3\text{-}RE}$-IL-10). |

**Supplementary Table S2. CC–GEMS receptor design and sequences.** DNA and amino-acid sequences of the cognate and non-cognate CC–GEMS receptor constructs listed in Table S1, including the A and B CC modules and the $\alpha_2$ linker connecting the CC module to the EpoR-derived receptor region. DNA and amino-acid sequences of the CC ligands are provided in Supplementary Fig. S1 for cysteine-engineered A′A′ CC ligands, including CC and $\mathbf{CC_{ditopic}}$, and in Supplementary Figs. S14 and S15 for AND-gated cysteine-engineered CC ligands.

| Component | DNA Sequence (5' to 3') | Amino acid sequence (N-term to C-term) |
|---|---|---|
| $\alpha_2$ linker | GGATCAGGGTCTGGAAGTGGGAGT | GSGSGSGS |
| A CC | GAGATCCAGCAACTGGAAGAGGAAATCGCCCAGCTGGA GCAGAAAAACGCCGCTCTGAAGGAAAAGAACCAGGCCC TGAAGTAC | EIQQLEEEIAQLEQKNAALKEKNQALKY |
| B CC | GAGAACGCCGCTCTGGAAGAGAAGATCGCCCAGCTGAA ACAGAAGAACGCCGCCCTGAAGGAAGAGATCCAGGCCC TGGAATAC | ENAALEEKIAQLKQKNAALKEEIQALEY |

**Supplementary Table S3. DNA oligonucleotides.** Oligonucleotides used in this study, including sequence, terminal modifications and intended function. Figure references indicate experiments in which each oligonucleotide was used.

| Name | Length | Sequence (5′→3′) | 5' modification | 3' modification | Role |
|---|---|---|---|---|---|
| $ssDNA_{scramble}$ | 34-nt | GAT AGG TGT ATT ACT TCC CGG ACA ACC TGC CCG T | | | Non-complementary control / sacrificial nuclease substrate. Fig. 2b; Fig. 2d; Fig. S8; Fig. 2f-g; Fig. S10; Fig. 3b; Fig. S11; Fig. 3e-f; Fig. S18; Fig. 4c; Fig. S20. |
| $ssDNA_{anchor}$ | 34-nt | TGG TGG TGG TTA GAC TCG TAT GTA GAA TGG TGG C | Biotin | | Streptavidin-binding anchor for $ssDNA_{top\_fluor}$ and $ligand_{ssDNA\text{-}CC}$, $ligand_{fluor}$, and $ligand_{activation}$. Fig. 1d; Fig. S13; Fig. 1f; Fig. S7; Fig. 2d; Fig. S8; Fig. S9; Fig. 3b; Fig. S11; Fig. 3e. |
| $ssDNA_{anchor\_fluor}$ | 34-nt | TGG TGG TGG TTA GAC TCG TAT GTA GAA TGG TGG C | Biotin | Cy3 | Cy3-labelled streptavidin-binding anchor. Fig. S9. |
| $ssDNA_{top\_fluor}$ | 26-nt | GCC ACC ATT CTA CAT ACG AGT CTA AC | Cy3 | | Cy3-labelled $ssDNA_{anchor}$ duplex strand. Fig. S5; Fig. S9. |
| $ssDNA_{Am}$ | 26-nt | GCC ACC ATT CTA CAT ACG AGT CTA AC | | AmMO | Amine-modified precursor used to prepare $ligand_{ssDNA\text{-}CC}$. Fig. 1b. |

| | | | | | |
|---|---|---|---|---|---|
| $ssDNA_{activation}$ | 26-nt | GCC ACC ATT CTA CAT ACG AGT CTA AC | Cy3 | AmMO | Cy3-labelled amine-modified precursor used to prepare $ligand_{activation}$. |
| $ssDNA_{input}$ | 34-nt | GCC ACC ATT CTA CAT ACG AGT CTA ACC ACC ACC A | | | $ssDNA_{anchor}$-cognate input strand. Fig. 1f; Fig. S7; Fig. 3b; Fig. S11; Fig. 3e-f. |
| $ssDNA_{AND1}$ | 26-nt | CAA GCG CTA TAT AGC TAT TCG CAA CG | Cy5 | AmMO | Amine-modified precursor used to prepare $ligand_{AND1}$. |
| $ssDNA_{AND2}$ | 26-nt | TGA TGT TAG TCG GAT GAA CGT CCT CT | Cy3 | AmMO | Amine-modified precursor used to prepare $ligand_{AND2}$. Fig. S17. |
| $ssDNA_{anchor\_AND1}$ | 34-nt | GCG TAT GCC GTT GCG AAT AGC TAT ATA GCG CTT G | Biotin | | Streptavidin-binding anchor for $ligand_{AND1}$. Fig. S19; Fig. 4c. |
| $ssDNA_{anchor\_AND2}$ | 34-nt | CAG TCC TGA GAG GAC GTT CAT CCG ACT AAC ATC A | Biotin | | Streptavidin-binding anchor for $ligand_{AND2}$. Fig. S19; Fig. 4c. |
| $ssDNA_{in\_AND1}$ | 34-nt | CAA GCG CTA TAT AGC TAT TCG CAA CGG CAT ACG C | | | $ssDNA_{anchor\_AND1}$-cognate input strand. Fig. S19; Fig. 4c. |
| $ssDNA_{in\_AND2}$ | 34-nt | TGA TGT TAG TCG GAT GAA CGT CCT CTC AGG ACT G | | | $ssDNA_{anchor\_AND2}$-cognate input strand. Fig. S19; Fig. 4c. |

**Supplementary Table S4. CC proteins.** CC proteins used in this study, with the corresponding expression construct and intended function. Figure references indicate experiments in which each CC protein was used.

| Name | Plasmid ID | Role |
|---|---|---|
| CC | pET28a(+)-SUMO-A' CC-$I_2$-A' CC | Original coiled-coil protein. Fig. S5. |
| $CC_{ditopic}$ | pET28a(+)-Cys12-SUMO-A' CC-$I_2$-A' CC | Cysteine-engineered coiled-coil protein precursor used to prepare $ligand_{ssDNA–CC}$, $ligand_{fluor}$, and $ligand_{activation}$. Fig. S2; Fig. 1b; Fig. S5; Fig. 2f. Fig. S10. |
| $CC_{AND1}$ | pET28a(+)-SUMO-A' CC-$I_4$-Γ CC-Cys230 | Cysteine-engineered AND-gate coiled-coil protein used to prepare $ligand_{AND1}$. Fig. S16; Fig. S17. |
| $CC_{AND2}$ | pET28a(+)-SUMO-A' CC-$I_4$-Γ' CC-Cys230 | Cysteine-engineered AND-gate coiled-coil protein used to prepare $ligand_{AND2}$. Fig. S16; Fig. S17. |

**Supplementary Table S5. ssDNA–CC ligand constructs.** ssDNA–CC ligand constructs used in this study, including their DNA and CC components, fluorophore labelling and intended function. Figure references indicate experiments in which each CC protein was used.

| Ligand name | DNA component | CC component | Fluorophore | Role |
|---|---|---|---|---|
| $Ligand_{ssDNA–CC}$ | $ssDNA_{Am}$ | $CC_{ditopic}$ | Unlabelled. | Base construct for DNA-programmed localization and release. Fig. 1b; Fig. S3; Fig. 2b; Fig. 2f; Fig. S10. |
| $Ligand_{fluor}$ | $ssDNA_{Am}$ | $CC_{ditopic}$ | Cy3-labelled; Cy3 was introduced by conjugation of Cy3–NHS ester to $CC_{ditopic}$. | Fluorescent ligand used in calibration, localization, retention and release experiments. Fig. 1d; Fig. S4; Fig. S13; Fig. S5; Fig. 1f; Fig. S7; Fig. 2d; Fig. S8; Fig. S9; Fig. 3b. Fig. S11. |
| $Ligand_{activation}$ | $ssDNA_{activation}$ | $CC_{ditopic}$ | Cy3-labelled oligonucleotide. | Used for SEAP and IL-10 proteinosome–cell communication. Fig. 3e-f; Fig. S20. |
| $Ligand_{AND1}$ | $ssDNA_{AND1}$ | $CC_{AND1}$ | Cy5-labelled oligonucleotide. | Used for AND-gated proteinosome–cell communication; ligand pairs with $ligand_{AND2}$. Fig. S17; Fig. S18; Fig. 4c. |
| $Ligand_{AND2}$ | $ssDNA_{AND2}$ | $CC_{AND2}$ | Cy3-labelled oligonucleotide. | Used for AND-gated proteinosome–cell communication; ligand pairs with $ligand_{AND1}$. Fig. S17; Fig. S18; Fig. 4c. |

**Supplementary Table S6. Fluorescence intensity and fold-change summary.** Overview of the number of proteinosomes analysed, mean fluorescence intensity ± standard deviation (SD). Fold changes for the indicated comparisons and corresponding figure.

| Figure | Compared groups | Proteinosomes (n) | Mean ± SD (RFU) | Compared conditions | Fold change | Fold change type |
|---|---|---|---|---|---|---|

| | | | | | | |
|---|---|---|---|---|---|---|
| 2d (RFU) | DMEM before (A) | 29 | 1,475.40 ± 892.80 | A-B | 3.24 | Before/ after fluorescence ratios |
| | DMEM after (B) | 30 | 455.22 ± 120.35 | | | |
| | SFM before (C) | 23 | 1,595.55 ± 942.10 | C-D | 3.39 | |
| | SFM after (D) | 30 | 470.52 ± 83.89 | | | |
| | Protection medium, before (E) | 19 | 1,313.00 ± 1,097.70 | E-F | 1.12 | |
| | Protection medium, after (F) | 30 | 1,169.25 ± 764.12 | | | |
| S5c (RFU) | ssDNA$_{top_fluor}$ (A) | 30 | 161.72 ± 5.56 | A-B | 1.03 | Reference / comparison fluorescence ratios |
| | | | | A-C | 2.75 | |
| | CC protein (B) | 30 | 167.87 ± 10.31 | A-D | 2.78 | |
| | | | | A-E | 10.22 | |
| | CC$_{ditopic}$ (C) | 30 | 443.69 ± 130.61 | B-C | 2.64 | |
| | | | | B-D | 2.70 | |
| | Ligand$_{fluor}$ (D) | 30 | 449.92 ± 94.19 | B-E | 9.90 | |
| | | | | C-D | 1.01 | |
| | Programmed (E) | 20 | 1,651.63 ± 1,121.75 | C-E | 3.72 | |
| | | | | D-E | 3.67 | |
| S9b (RFU) | DMEM before, AF 546-streptavidin (A) | 30 | 1,299.90 ± 1,011.89 | A-B | 1.11 | Before/ after fluorescence ratios |
| | DMEM after, AF 546-streptavidin (B) | 30 | 1,171.75 ± 930.64 | | | |
| | SFM before, AF 546-streptavidin (C) | 28 | 1,000.70 ± 756.20 | C-D | 0.78 | |
| | SFM after, AF 546-streptavidin (D) | 30 | 1,281.35 ± 815.78 | | | |
| | DMEM before, ssDNA$_{anchor_fluor}$ (E) | 21 | 1,756.74 ± 1,252.48 | E-F | 3.94 | |
| | DMEM after, ssDNA$_{anchor_fluor}$ (F) | 30 | 445.67 ± 121.38 | | | |
| | SFM before, ssDNA$_{anchor_fluor}$ (G) | 30 | 1,563.06 ± 1,084.62 | G-H | 3.08 | |
| | SFM after, ssDNA$_{anchor_fluor}$ (H) | 30 | 507.38 ± 127.85 | | | |
| | DMEM before, ssDNA$_{top_fluor}$ (I) | 24 | 2,040.76 ± 1,268.14 | I-J | 4.86 | |
| | DMEM after, ssDNA$_{top_fluor}$ (J) | 30 | 419.64 ± 118.99 | | | |
| | SFM before, ssDNA$_{top_fluor}$ (K) | 27 | 1,681.74 ± 1,084.57 | K-L | 3.75 | |
| | SFM after, ssDNA$_{top_fluor}$ (L) | 30 | 448.23 ± 100.22 | | | |
| | DMEM before, ligandfluor (M) | 26 | 1,485.16 ± 1,107.75 | M-N | 3.32 | |
| | DMEM after, ligand$_{fluor}$ (N) | 30 | 447.41 ± 94.14 | | | |
| | SFM before, ligand$_{fluor}$ (O) | 27 | 1,631.29 ± 905.82 | O-P | 3.90 | |
| | SFM after, ligand$_{fluor}$ (P) | 30 | 417.97 ± 117.34 | | | |
| S19b (RFU) | Ligand$_{AND1}$, 00 (A) | 170 | 926.57 ± 804.37 | A-B | 1.18 | Reference / comparison fluorescence ratios |
| | | | | A-C | 3.20 | |
| | Ligand$_{AND1}$, 01 (B) | 186 | 783.45 ± 747.50 | A-D | 3.07 | |
| | | | | B-C | 2.70 | |
| | Ligand$_{AND1}$, 10 (C) | 219 | 290.01 ± 22.33 | B-D | 2.60 | |
| | | | | C-D | 1.04 | |
| | Ligand$_{AND1}$, 11 (D) | 155 | 301.68 ± 19.60 | E-F | 2.17 | |
| | | | | E-G | 1.02 | |
| | Ligand$_{AND2}$, 00 (E) | 235 | 719.69 ± 376.43 | E-H | 2.00 | |
| | Ligand$_{AND2}$, 01 (F) | 204 | 332.22 ± 33.20 | F-G | 2.12 | |
| | Ligand$_{AND2}$, 10 (G) | 254 | 705.71 ± 333.90 | F-H | 1.08 | |
| | Ligand$_{AND2}$, 11 (H) | 197 | 358.22 ± 42.14 | G-H | 1.97 | |

**Supplementary Table S7. Unpaired two-tailed t-test summary.** Overview of triplicate values, fold change, p-values, t-values and degrees of freedom (df) for unpaired, two-tailed t-tests, indicating the compared groups and corresponding Figure. ns, P > 0.05; *, P ≤ 0.05; **, P ≤ 0.01; ***, P ≤ 0.001; ****, P ≤ 0.0001.

| Figure | Compared groups | Triplicate values | Fold change | p-value summary | p-value | t-value | df |
|---|---|---|---|---|---|---|---|
| 2g | Standard medium | 1.607490; 1.701429; 1.540102 | 3.23 | *** | 0.0009895 | 23.13 | 2.258 |
| (AU, 540nm) | Protection medium | 0.522531; 0.495163; 0.482082 | | | | | |
| 3b | Scrambled input | 1,803.31; 1,925.78; 2,048.25 | 8.30 | ** | 0.00164 | 23.88 | 2.025 |
| (RFU) | Complementary input | 222.22; 232.02; 241.82 | | | | | |
| 3c | Cell incubation | 0.100756; 0.119634; 0.081878 | 1.40 | ns | 0.1886 | 1.697 | 2.991 |
| (AU 540nm) | Proteinosome-cell co-incubation | 0.141159; 0.177829; 0.104488 | | | | | |

**Supplementary Table S8. One-way ANOVA and Tukey multiple-comparisons summary.** Overview of triplicate values, F-values, p-values and degrees of freedom (df) for one-way ANOVAs, with Tukey's multiple comparison test, indicating the compared groups and corresponding Figure. ns, P > 0.05; *, P ≤ 0.05; **, P ≤ 0.01; ***, P ≤ 0.001; ****, P ≤ 0.0001.

| Fig. | Compared groups (ANOVA) | Triplicate values | F-value | p-value summary (ANOVA) | Compared groups (multiple comparisons) | Fold change | Adjusted p-value (multiple comparisons) | df |
|---|---|---|---|---|---|---|---|---|
| 2f (U/L per viable cell) | Standard medium, no-ligand (A) | $1.08378 \times 10^{-4}$; $9.5294 \times 10^{-5}$; $1.08155 \times 10^{-4}$ | 73.42 | **** (p < 0.0001) | A-B | 23.92 | **** (p < 0.0001) | 12 |
| | | | | | A-C | 20.33 | **** (p < 0.0001) | |
| | | | | | A-D | 1.12 | ns (p = 1.0) | |
| | Standard medium, CC$_{ditopic}$ (B) | $2.04098 \times 10^{-3}$; $2.73347 \times 10^{-3}$; $2.68469 \times 10^{-3}$ | | | A-E | 7.05 | * (p = 0.0253) | |
| | | | | | A-F | 7.55 | * (p = 0.0150) | |
| | | | | | B-C | 1.18 | ns (p = 0.2919) | |
| | Standard medium, ligand$_{ssDNA-CC}$ (C) | $2.18431 \times 10^{-3}$; $1.77389 \times 10^{-3}$; $2.38154 \times 10^{-3}$ | | | B-D | 21.40 | **** (p < 0.0001) | |
| | | | | | B-E | 3.39 | **** (p < 0.0001) | |
| | | | | | B-F | 3.17 | **** (p < 0.0001) | |
| | Protection medium, no-ligand (D) | $1.25075 \times 10^{-4}$; $1.23676 \times 10^{-4}$; $9.97675 \times 10^{-5}$ | | | C-D | 18.19 | **** (p < 0.0001) | |
| | | | | | C-E | 2.88 | **** (p < 0.0001) | |
| | Protection medium, CC$_{ditopic}$ (E) | $7.70206 \times 10^{-4}$; $7.75622 \times 10^{-4}$; $6.53641 \times 10^{-4}$ | | | C-F | 2.70 | **** (p < 0.0001) | |
| | | | | | D-E | 6.31 | **** (p < 0.0001) | |
| | Protection medium, ligand$_{ssDNA-CC}$ (F) | $8.02149 \times 10^{-4}$; $8.06394 \times 10^{-4}$; $7.46764 \times 10^{-4}$ | | | D-F | 6.76 | **** (p < 0.0001) | |
| | | | | | E-F | 1.07 | ns (p = 0.9995) | |
| 3e (U/L) | Cognate receptor, no ligand (A) | 80.7974704; 55.62036972; 248.454612 | 71.36 | **** (p < 0.0001) | A-B | 6.18 | **** (p < 0.0001) | 12 |
| | | | | | A-C | 1.27 | ns (p = 0.9917) | |
| | | | | | A-D | 6.51 | **** (p < 0.0001) | |
| | Cognate receptor, free ligand (B) | 831.0718602; 815.2925492; 732.0626604 | | | A-E | 3.46 | ns (p = 0.6935) | |
| | | | | | A-F | 2.22 | ns (p = 0.8612) | |
| | | | | | B-C | 4.85 | **** (p < 0.0001) | |
| | Cognate receptor, proteinosomes + scrambled DNA (C) | 228.4962179; 76.72552525; 185.3495828 | | | B-D | 1.05 | ns (p = 0.9820) | |
| | | | | | B-E | 21.41 | **** (p < 0.0001) | |
| | | | | | B-F | 13.70 | **** (p < 0.0001) | |
| | Cognate receptor, proteinosomes + complementary DNA (D) | 721.1499312; 922.2449734; 860.7854037 | | | C-D | 5.10 | **** (p < 0.0001) | |
| | | | | | C-E | 4.42 | ns (p = 0.3840) | |
| | Noncognate receptor, proteinosomes + | 45.08349623; 29.77042791; 36.2604831 | | | C-F | 2.83 | ns (p = 0.5615) | |
| | | | | | D-E | 22.54 | **** (p < 0.0001) | |

| | | | | | | | | |
|---|---|---|---|---|---|---|---|---|
| | scrambled DNA (E) | | | | | | | |
| | Noncognate receptor, proteinosomes + complementary DNA (F) | 19.17457613; 135.7593073; 18.62318143 | | | D-F | 14.43 | **** (p < 0.0001) | |
| | | | | | E-F | 1.56 | ns (p = 0.9993) | |
| 4c (U/L) | 00 | 133.2062721; 141.3078495; 145.1952451 | 1224.78 | **** (p < 0.0001) | 00-01 | 1.04 | ns (p = 0.9314) | |
| | | | | | 00-10 | 1.12 | ns (p = 0.2463) | |
| | 01 | 138.6867509; 148.0682414; 150.0698076 | | | 00-11 | 3.28 | **** (p < 0.0001) | |
| | | | | | 00-A | 3.55 | **** (p < 0.0001) | |
| | 10 | 150.2127766; 157.3408031; 161.0920377 | | | 01-10 | 1.07 | ns (p = 0.6151) | |
| | | | | | 01-11 | 3.15 | **** (p < 0.0001) | |
| | 11 | 445.8046156; 461.4086622; 469.9051064 | | | 01-A | 3.41 | **** (p < 0.0001) | |
| | | | | | 10-11 | 2.94 | **** (p < 0.0001) | |
| | Free ligand pair (A) | 482.125553; 502.5020414; 503.993004 | | | 10-A | 3.18 | **** (p < 0.0001) | |
| | | | | | 11-A | 1.08 | ** (p = 0.0035) | |
| S18b (U/L) | 00 | 172.80; 163.28; 162.66 | 32.88 | **** (p < 0.0001) | 00-01 | 1.05 | ns (p = 0.9987) | 8 |
| | | | | | 00-10 | 1.20 | ns (p = 0.9166) | |
| | 01 | 165.15; 176.53; 179.83 | | | 00-11 | 3.56 | *** (p = 0.0001476) | |
| | | | | | 01-10 | 1.14 | ns (p = 0.9592) | |
| | 10 | 185.98; 185.61; 224.91 | | | 01-11 | 3.41 | *** (p = 0.000168) | |
| | 11 | 492.12; 556.41; 729.15 | | | 10-11 | 2.98 | *** (0.0002604) | |
| S20c (ng/mL) | No-ligand (A) | 0.50259; 1.41521; 0.31165 | 100.38 | **** (p < 0.0001) | A-B | 29.99 | **** (p < 0.0001) | 8 |
| | | | | | A-C | 1.98 | ns (p = 0.9536) | |
| | Free ligand (B) | 23.06893; 21.80356; 21.98479 | | | A-D | 13.96 | *** (p = 0.0006) | |
| | | | | | B-C | 15.15 | **** (p < 0.0001) | |
| | Proteinosomes + scrambled DNA (C) | 1.23074; 1.74531; 1.43786 | | | B-D | 2.15 | *** (p = 0.0001) | |
| | Proteinosomes + complementary DNA (D) | 13.45081; 6.81327; 10.85534 | | | C-D | 7.05 | ** (p = 0.0011) | |

#### Supplementary References


[73] M. A. Unger, H. P. Chou, T. Thorsen, A. Scherer, and S. R. Quake, “Monolithic microfabricated valves and pumps by multilayer soft lithography,” Science 288 (2000): 113–116. https://doi.org/10.1126/science.288.5463.113.
